# Pathological meprin α expression associated with degradation of dermokine drives a psoriasis-like skin phenotype in a genetic mouse model

**DOI:** 10.1101/2025.05.23.655724

**Authors:** Vasco Köhling, Florian Peters, Inez Götting, Emil Fries, Niklas Beck, Fred Armbrust, Silje Beckinger, Cynthia Bülck, Vahap Canbay, Inken Harder, Marion Mengel, Malina Rüffer, Kira Bickenbach, Konstantinos Kalogeropoulos, Michaela Schweizer, Neele Schumacher, Michael Haase, Ronald Naumann, Ulrich auf dem Keller, Christoph Becker-Pauly, Sascha Rüffer

## Abstract

Keratinocyte proliferation and differentiation is regulated via proteolytic networks. Dysregulation of proteases within these networks can cause hyperproliferative and inflammatory skin disorders. In healthy skin the metalloprotease meprin α is localized in the *stratum basale*. In contrast, in wound healing tissue and psoriatic lesions increased meprin α levels are found in the upper epidermal layers. We developed a transgenic mouse model for inducible expression of pathological meprin α levels (K5Mα) to investigate its epidermal degradome and identify molecular links to keratinocyte proliferation and skin inflammation.

K5Mα mice developed a severe skin phenotype characterized by hyperkeratosis, acanthosis and parakeratosis accompanied by increased transepidermal water loss and a strong inflammatory response within six days after induction of meprin α overexpression. Histological and molecular analyses showed that increasing meprin α expression correlates with meprin α activity and keratinocyte hyperproliferation. Proteomics analyses revealed massive changes in proteins associated with keratinocyte differentiation and epidermal barrier integrity already three days after induction. N-terminomics data indicated a dominant chymotryptic activity with elevated proteolytic turnover of proteins associated with the cytoskeleton, cellular stress responses and cell adhesion. By filtering for cleavage sites that match with the specificity of meprin α, we identified highly elevated dermokine-derived peptides. Subsequent mass spectrometric analyses validated dermokine as a novel substrate of meprin α and identified the cleavage site, which is highly conserved in mammals.

Based on the striking similarities with the phenotype reported for dermokine αβγ^-/-^ mice, we propose meprin α as a central regulator of keratinocyte proliferation and leukocyte recruitment by proteolytic inactivation of dermokine. Hence, pathological meprin α activity could be a driver of hyperproliferative, inflammatory skin disorders like psoriasis vulgaris.

## Introduction

Epidermal homeostasis and barrier function is regulated by a complex proteolytic network that controls keratinocyte proliferation and differentiation. Proteases process proteins that form the cornified envelope and regulate the proliferation and differentiation of keratinocytes by the release and degradation of growth factors as well as cleavage of cell-cell and cell-matrix interactions (reviewed in references^1–3^). The expression of proteases in the epidermis is tightly regulated by the differentiation status of keratinocytes and their activity is controlled by the protease inhibitors as well as other proteases within the network. Thereby, it is ensured that the activity of each protease is restricted to distinct layers of the epidermis^1^. This local restriction is very important because mislocalized protease activities can cause severe defects in keratinocyte differentiation, which in turn can lead to loss of barrier integrity, keratinocyte hyperproliferation and/or initiation of harmful inflammatory responses. Netherton syndrome (NS) represents a prominent example of such a skin disease that is based on epidermal protease dysregulation^4^. However, there is increasing evidence that dysregulation of the proteolytic network also plays a role in highly prevalent skin diseases like acute dermatitis (AD) and psoriasis vulgaris (PV)^5–7^.

The metalloproteases meprin α and meprin β are part of the epidermal protease network. Although both exhibit similar cleavage specificities with a preference for negatively charged amino acids in P1, P1’ and P2’ position, there are major differences in the synthesis, regulation and localization of meprins^8^. Both meprin α and meprin β are synthesized as zymogens activated by tryptic proteases^9–12^. We showed that several members of the kallikrein-related peptidases family function as activators of meprins while others are activated by meprins^9^. Meprin β is synthesized as a type I transmembrane protein that can be released from the cell surface *via* ectodomain shedding^13–15^. The ectodomain of meprin α is canonically released by furin already within the secretory pathway and, therefore, meprin α is constitutively secreted^10,16,17^. Moreover, meprin β expression is restricted to the *stratum granulosum* while meprin α is synthesized in the *stratum basale*, indicating that meprin β regulates terminal keratinocyte differentiation, whereas meprin α might control keratinocyte proliferation^18,19^. Recently, we showed that meprin β regulates terminal keratinocyte differentiation by shedding of the proteoglycan syndecan-1 and provided evidence that excessive interleukin-18 activation by meprin β leads to an upregulated melanogenesis causing hyperpigmentation in mice^20^. However, the epidermal degradome of meprin α is poorly characterized. Currently, the only hints towards the epidermal function of meprin α arise from its levels and localization detected in healthy skin compared to skin samples from PV patients.

Here, in psoriatic lesions increased meprin α levels have been found mislocalized to the *stratum spinosum*^18^. However, it is not known whether dysregulated meprin α activity can be causative for the onset or progression of hyperproliferative and/or inflammatory skin disorders.

In order to address these questions, we developed a transgenic mouse model on C57BL/6 background that allows epidermis-specific induction of meprin α overexpression by tamoxifen treatment, thereby, mimicking increased meprin α levels observed in PV. In case these mice develop a pathological skin phenotype, our aim was to characterize the phenotype on histological and molecular level, investigate alterations within the proteolytic network and to identify new epidermal substrates of meprin α using proteomics approaches.

## Results

### K5Mα mice develop a severe ichthyosis throughout the body upon induction of meprin α overexpression

K5Mα mice were generated by insertion of a construct comprising the C-terminal HA-tagged murine meprin α cDNA sequence with an upstream stop cassette flanked by loxP sites and a CAG promotor into the Rosa26 locus of C57BL/6J mice. These mice were crossed with a driver strain that harbors the information for the CreER^T2^ fusion protein downstream of the Krt5 promoter, which is highly active in keratinocytes of the *stratum basale*. Respective control mice did not express CreER^T2^ but harbored the transgenic Rosa26 locus (**Figure 1 A**). For induction of CreER^T2^ translocation, tamoxifen was applied by supplemented food pellets (**Figure 1 B**). Within the first week, we noticed reddening of the ears as well as small scabby lesions in the ears, around the muzzle and in the neck accompanied by regular scratching behavior indicating mild itching (**Figure 1 C**). In the following ten days, these initially small lesions spread across the entire body. Both intense redness and scaling as well as skin stiffening were observed. We also noted very frequent scratching and prominent digging behavior indicating severe itching leading to hair loss and wounding in respective skin areas (**Figure 1 C**). At this stage of the phenotype, mice were euthanized for further analyses (**Figure 1 C**). Based on our observations we developed a cumulative scoring system to rate and track the skin phenotype development (**Figure 1 C**, red numbers). Correlation analyses highlighted an exponential amplification of the skin phenotype severity with a maximum score of 20 reached within 17 days after the start of tamoxifen treatment (**Figure 1 D**). Of note, control mice developed none of the described phenotypic changes over the whole experimental course. Between day 10 to 16 K5Mα mice developed a very bad overall condition, so we had to remove them from the experiment before day 17 and some K5Mα mice even unexpectedly died. Subsequent histological examinations revealed a prominent ichthyosis in the skin of K5Mα mice (**Figure 1 E**) accompanied by significantly higher meprin α activity in the skin of K5Mα compared to control mice (0.89 ± 0.08 *vs.* 0.35 ± 0.09; p=0.004) (**Figure 1 F**).

**Figure 1:**
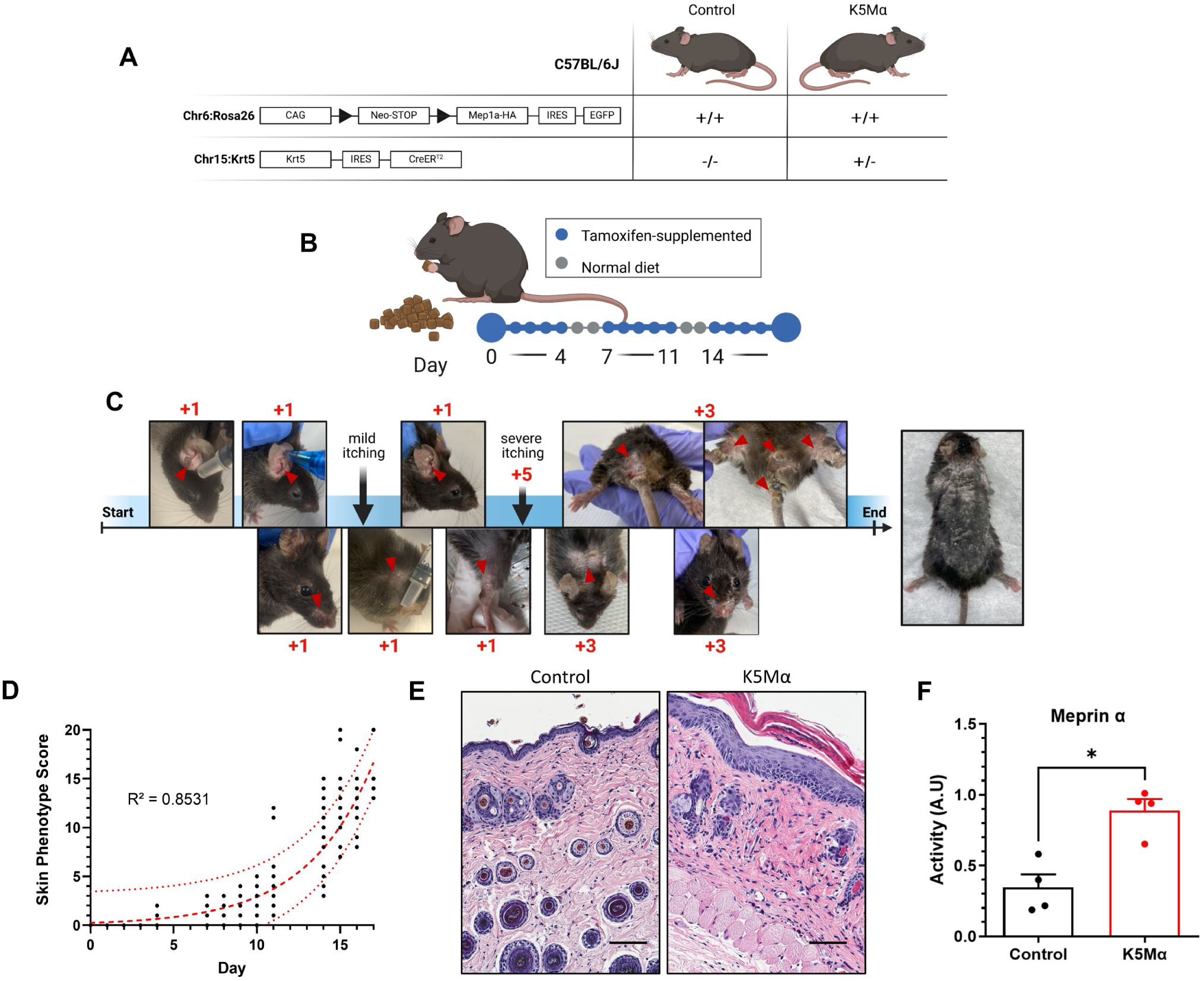
K5Mα mice develop a severe ichthyosis throughout the body upon induction of meprin α overexpression. **(A)** Scheme illustrates the genotype of transgenic mice designated as “control” and “K5Mα”. Both strains were bred on a C57BL7/6J background harboring a homozygous (+/+) insertion in the Rosa26 locus composed of a CAG-promotor (CAG), a loxP site (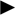)- flanked neomycin resistance (Neo)/STOP cassette, a cDNA sequence encoding HA-tagged murine meprin α (*Mep1a*-HA), an internal ribosomal entry site (IRES) and a cDNA sequence encoding enhanced green fluorescent protein (*EGFP*). K5Mα mice, but not control mice, also carry heterozygous (+/-) downstream of the *Krt5* gene an IRES followed by a gene encoding the fusion protein CreER^T2^ composed of Cre recombinase and a mutant estrogen receptor ligand-binding domain that specifically binds tamoxifen. **(B)** Illustration of the feeding scheme for tamoxifen treatment of control and K5Mα mice. Tamoxifen-supplemented food pellets were fed *ad libitum* for five days followed by two days of normal diet. **(C)** Development of the skin phenotype in K5Mα mice after start of the tamoxifen treatment. Representative images of the macroscopic skin examination. Red arrowheads (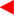) mark the examined skin area and red number (1-5) the assigned scores. **(D)** Cumulative skin phenotype scores calculated from macroscopic skin examination of K5Mα mice (n=22) from day zero to day 17 after start of tamoxifen treatment. R-square (R^2^) value was calculated by non-linear regression based on an exponential growth equation. **(E)** Representative microscopical images of hematoxylin and eosin stained skin sections from control and K5Mα mice after 17 days of tamoxifen treatment. Scale bar = 75 µm. **(F)** Meprin α activity in skin lysates of control (n=4) and K5Mα (n=4) mice after 17 days of tamoxifen treatment. Activity is plotted in arbitrary units (A.U.). Bar charts and error bars display mean and standard error of the mean. Statistically significant differences (p < 0.05) are labeled by an asterisk (*). Graphical illustrations in **(A)**, **(B)** and **(C)** were created with BioRender.com.

### K5Mα mice develop a severe hyperkeratosis, acanthosis and parakeratosis accompanied by barrier integrity defects

In order to track the development of the skin phenotype more precisely, we treated shaved skin areas at the flank of control and K5Mα mice by a single local, cutaneous application of 4-hydroxytamoxifen (TAM, 125 µg/cm² in DMSO) and sampled respective skin areas daily from euthanized mice over a time course of six days for subsequent analyses (**Figure 2 A**). Within these six days, we observed development of the full phenotype as described for diet-based tamoxifen treatment. While control mice developed no histopathological changes in the skin, hyperplastic epidermal lesions were observed in K5Mα mice already on day two after TAM treatment (**Figure 2 B**). Thickness of the epidermis from K5Mα but not control mice significantly increased over the experimental period (**Figure 2 C**), while dermal thickness was not altered in both K5Mα and control mice (**Figure 2 D**). A closer look at the epidermis showed a physiological architecture and proportion of each epidermal layer in the skin of control mice (**Figure 2 E**, Control), while prominent hyperkeratosis and parakeratosis was observed in the epidermis of K5Mα mice (**Figure 2 E**, K5Mα (I)). Moreover, we identified unusual strongly granulated keratinocytes within the *stratum spinosum* (**Figure 2 E**, K5Mα (II)) and found evidence for proliferating keratinocytes in the *stratum spinosum* (**Figure 2 E**, K5Mα (III)+(IV)). Transmission electron microscopic analyses revealed a striking accumulation of lipid droplets in the *stratum corneum* of K5Mα mice (**Figure 2 F**). Hemidesmosomes (black arrowheads), *lamina lucida* (yellow arrows) and *lamina densa* (black arrows) were detectable in the skin of both control and K5Mα mice indicating an intact epidermal basement membrane (**Figure 2 G**). Interestingly, the *lamina densa* appeared to be more electron dense in the skin of K5Mα than control mice.

**Figure 2:**
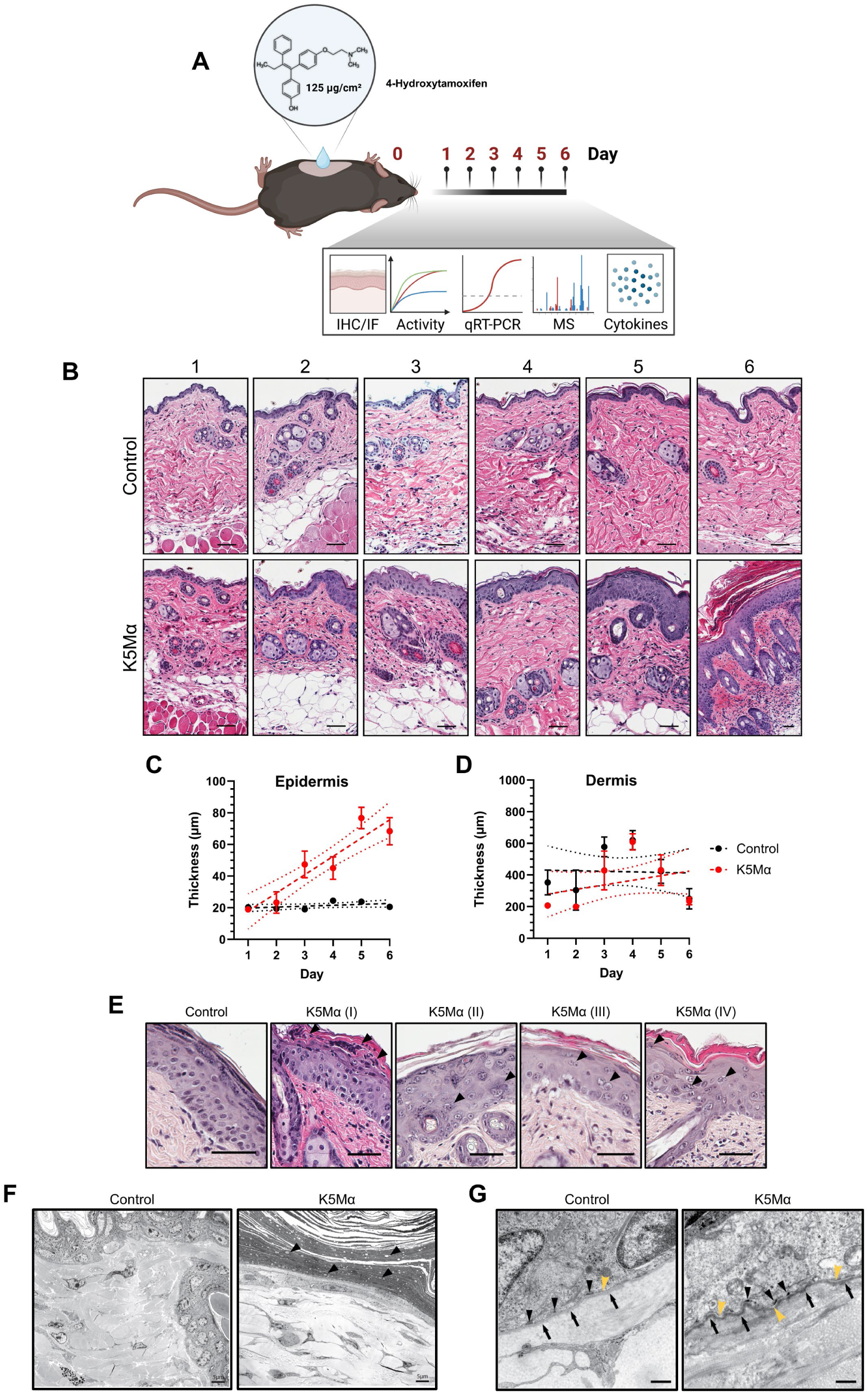
K5Mα mice develop a severe hyperkeratosis, acanthosis and parakeratosis with an intact epidermal basement membrane and lipid droplet accumulation in the *stratum corneum*. **(A)** Illustration of the topical tamoxifen treatment of control and K5Mα mice. Shaved skin areas were treated once on day zero with 125 µg/cm² 4-hydroxytamoxifen (TAM) in DMSO. The following six days, each day control and K5Mα mice were euthanized for sampling of the skin areas and subsequent analyses by immunohistochemistry (IHC), immunofluorescence (IF), real-time polymerase chain reaction (qRT-PCR), mass spectrometry (MS) and cytokine quantification. **(B)** Representative microscopical images of hematoxylin and eosin stained skin sections from control and K5Mα mice on day one to six after topical TAM treatment. Scale bar = 67 µm. **(C+D)** Image-based quantification of the **(C)** epidermal and **(D)** dermal thickness in skin sections from control (n=4 per day) and K5Mα (n=4 per day) mice at days one to six after TAM treatment. Data points and error bars display mean values and respective standard errors of the mean (SEM). Results from simple linear regression (dashed lines) and respective 95% confidence band of the best-fit line (dotted lines) are displayed. **(E)** Representative microscopical images of hematoxylin and eosin stained skin sections from control and K5Mα mice after topical TAM treatment. Black arrowheads (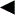) mark pathological findings in the epidermis of K5Mα mice including hyperkeratosis and parakeratosis (K5Mα (I)), highly granulated keratinocytes in the spinous layer accompanied by acanthosis (K5Mα (II)) as well as suprabasal proliferating keratinocytes (K5Mα (III + IV)). Scale bar = 50 µm. **(F+G)** Representative transmission electron microscope images of skin sections from TAM-treated control and K5Mα mice with a focus on **(F)** the upper spinous layer, *stratum granulosum* and *stratum corneum* or **(G)** the basal layer and the basement membrane. **(F)** Black arrowheads (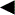) mark lipid droplets in the stratum corneum. Scale bars = 5 µm **(G)** Black arrowheads (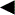) mark hemidesmosomes, yellow arrowheads (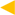) the *lamina lucida* and black arrows (→) the *lamina densa*. Scale bars = 500 nm. Graphical illustration in **(A)** was created with BioRender.com.

In an attempt to rescue the skin phenotype of K5Mα mice we repeated the topical induction of meprin α overexpression and treated the respective areas twice daily with actinonin (100 µg/cm² in DMSO), a potent hydroxamate-based inhibitor of meprin α^21^ (**Supplementary Figure 1 A**). However, actinonin treatment neither prevented the development of acanthosis, hyperkeratosis and parakeratosis (**Supplementary Figure 1 B**) nor led to reduced or delayed epidermal hyperplasia (**Supplementary Figure 1 C**). Moreover, meprin α activity in the skin of actinonin-treated K5Mα was only slightly but not significantly reduced compared to DMSO treated K5Mα mice (3.51 ±0.13 *vs.* 3.22 ±0.11) (**Supplementary Figure 1 D**). In order to examine whether development of the K5Mα skin phenotype was associated with an impaired barrier integrity, we measured daily the transepidermal water-loss (TEWL) on the DMSO and actinonin-treated skin areas of control and K5Mα mice. We detected an increasing TEWL on the skin areas of K5Mα mice treated with DMSO while TEWL in the respective control group was on a constant physiological level over the whole experimental course (**Supplementary Figure 1 E+G**). In comparison to DMSO-treated K5Mα mice, a similar increase in TEWL was measured on skin areas of K5Mα mice treated with actinonin (**Supplementary Figure 1 F+G**). Interestingly, TEWL values on skin areas of actinonin-treated control mice were significantly higher than in DMSO-treated control mice at day six (12.16 ±1.65 *vs*. 5.51 ±0.78; p=0.0045) (**Supplementary Figure 1 G**).

### Meprin α expression correlates with keratinocyte hyperproliferation in the skin of K5Mα mice

The rapidly increasing epidermal thickness as well as the evidence for suprabasal proliferation of keratinocytes indicated keratinocyte hyperproliferation as a potential cause of the acanthosis observed in K5Mα mice. Therefore, we assessed meprin α expression and activity in skin lysates as well as meprin α levels and Ki67 status in skin sections (**Figure 3 A**). In skin sections of control mice, we detected low but constant meprin α levels from day one to six (416.23 ±13.03 mean pixel intensity (MPI)), while high meprin α levels were observed already in small areas at day two (596.36 ±111.55 MPI) in K5Mα mice. Already three days after TAM treatment, meprin α levels were significantly higher in the skin of K5Mα mice than in control mice (1535.10 ±399.44 *vs*. 389.38 ±10.79 MPI) and further increased to a maximum at day five (2578.31 ±159.91 MPI) (**Figure 3 B**). In line with these observations, both *Mep1a* mRNA levels (**Figure 3 C**) and meprin α activity (**Figure 3 D**) significantly increased from day one to day three in the skin of K5Mα mice while *Mep1a* mRNA levels and meprin α activity in the skin of control mice remained at constant levels. Quantification of Ki67-positive keratinocytes in the skin of control mice revealed a mild gradual decrease from day one to six (**Figure 3 E**). The initially elevated proportion of Ki67- positive keratinocytes in the skin of control mice might be attributed to skin irritations in response to the shaving 24 h before TAM treatment. In contrast, the proportion of Ki67-positive keratinocytes in the skin of K5Mα mice significantly increased from day one to six (**Figure 3 E**). At day six, Ki67 was detectable in the nuclei of almost 50% of the keratinocytes in the skin of K5Mα mice. Of note, Ki67-positive keratinocytes were not exclusively located in the *stratum basale* but also found in the *stratum spinosum*. Moreover, we identified an accumulation of Ki67- positive cells in the dermis (**Figure 3 A**).

**Figure 3:**
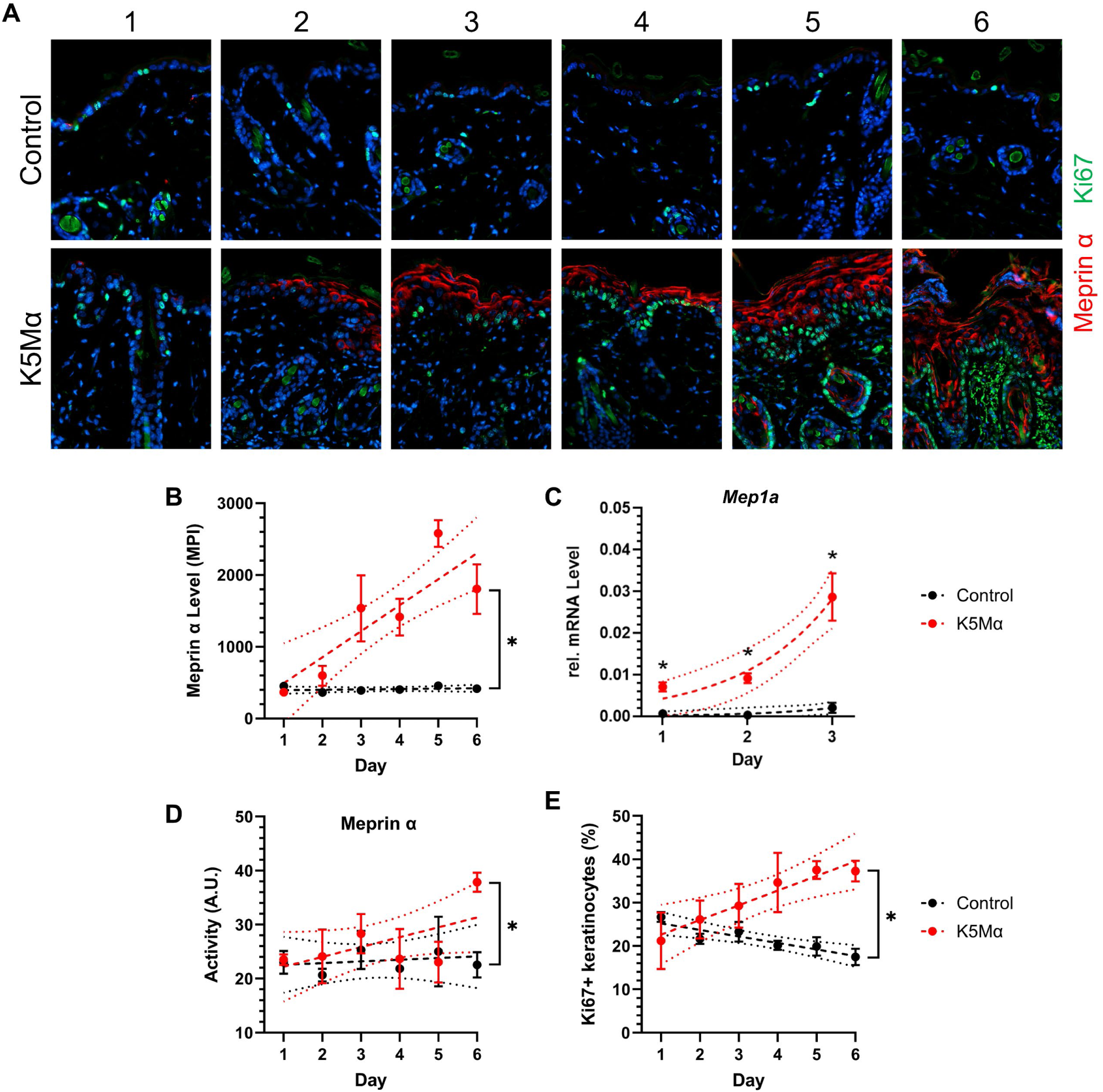
Ki67 status indicates initiation of keratinocyte hyperproliferation upon induction of meprin α overexpression in K5Mα mice. **(A)** Representative immunofluorescence images of skin sections from control and K5Mα mice on day one to six after topical TAM treatment. Meprin α (red) and Ki67 (green) were detected by indirect immunofluorescence staining. Nuclei were stained with 4’,6-diamino-2-phenylindol (DAPI). **(B)** Image-based quantification of the mean pixel intensity (MPI) from meprin α staining in skin sections from control (n=4 per day) and K5Mα (n=4 per day) mice at days one to six after TAM treatment. Data points and error bars display mean values and respective standard errors of the mean (SEM). Results from simple linear regression (dashed lines) and respective 95% confidence band of the best-fit line (dotted lines) are displayed. **(C)** Relative mRNA levels of *Mep1a* in skin samples from control (n=5 per day) and K5Mα (n=5 per day) mice at days one to three after TAM treatment. Mep1a levels were normalized to *Gapdh* levels in the respective sample by 2^-ΔΔCt^-method. Data points and error bars display mean values and respective SEM. Results from non-linear regression based on an exponential growth equation (dashed lines) and respective 95% confidence band of the best-fit line (dotted lines) are displayed. **(D)** Meprin α activity in skin lysates of control (n=4 per day) and K5Mα (n=4 per day) mice at days one to six after TAM treatment. Activity is plotted in arbitrary units (A.U.). Data points and error bars display mean values and respective SEM. Results from simple linear regression (dashed lines) and respective 95% confidence band of the best-fit line (dotted lines) are displayed. **(E)** Image-based quantification of the proportion of Ki67-positive keratinocytes in skin sections from control (n=4 per day) and K5Mα (n=4 per day) mice at days one to six after TAM treatment. Data points and error bars display mean values and respective SEM. Results from simple linear regression (dashed lines) and respective 95% confidence band of the best-fit line (dotted lines) are displayed. **(B-E)** Statistically significant differences (p < 0.05) are labeled by an asterisk (*).

In order to examine the proliferation and terminal differentiation of keratinocytes from K5Mα mice in more detail, we isolated keratinocytes from the tail skin of K5Mα mice (**Supplementary Figure 2 A**). Having validated meprin α expression and activity in isolated keratinocytes after treatment with TAM (**Supplementary Figure 2 B-F**), we found evidence for an increased proliferation of keratinocytes due to meprin α overexpression and activity (**Supplementary Figure 2 G-I)** but did not observe alterations in calcium-induced terminal differentiation (**Supplementary Figure 2 J-L)**.

In summary, meprin α overexpression was successfully induced in the skin of K5Mα mice by topical TAM treatment. Moreover, significantly increasing meprin α activity was detected in the skin from day one to six. K5Mα mice developed a severe pathological skin phenotype characterized by stepwise development of acanthosis, hyperkeratosis and parakeratosis accompanied by increasing TEWL, indicating epidermal barrier defects. Assessment of the Ki67 status and electron microscopic analyses *in situ* as well as *in vitro* analyses of isolated keratinocytes indicated that the skin phenotype development was initiated by keratinocyte hyperproliferation and not by defects in the basement membrane or terminal differentiation. Topical treatment with the meprin α inhibitor actinonin did neither delay nor rescue the phenotype progression.

### Alterations in the skin proteome of K5Mα mice support that keratinocyte hyperproliferation is initiated by epidermal barrier alarm mechanisms

In order to characterize changes in the skin proteome that go along with the hyperproliferative phenotype observed in K5Mα mice and to find evidence for the underlying molecular mechanisms, we performed proteomics analyses with skin samples from control and K5Mα mice sampled at day one to day 6 (**Figure 4 A**). On day one after TAM treatment, we identified ten significantly upregulated (≥ 2-fold) and six significantly downregulated (≤ 0.5-fold) proteins in the skin of K5Mα mice compared to control mice (**Figure 4 B**). Among the up- and downregulated proteins we found by gene ontology analysis several muscle-associated proteins (**Figure 4 C**; Cytoskeleton) as well as transcriptional or translational regulators and proteins associated with endosomal transport (**Supplementary Figure 3 A**).

**Figure 4:**
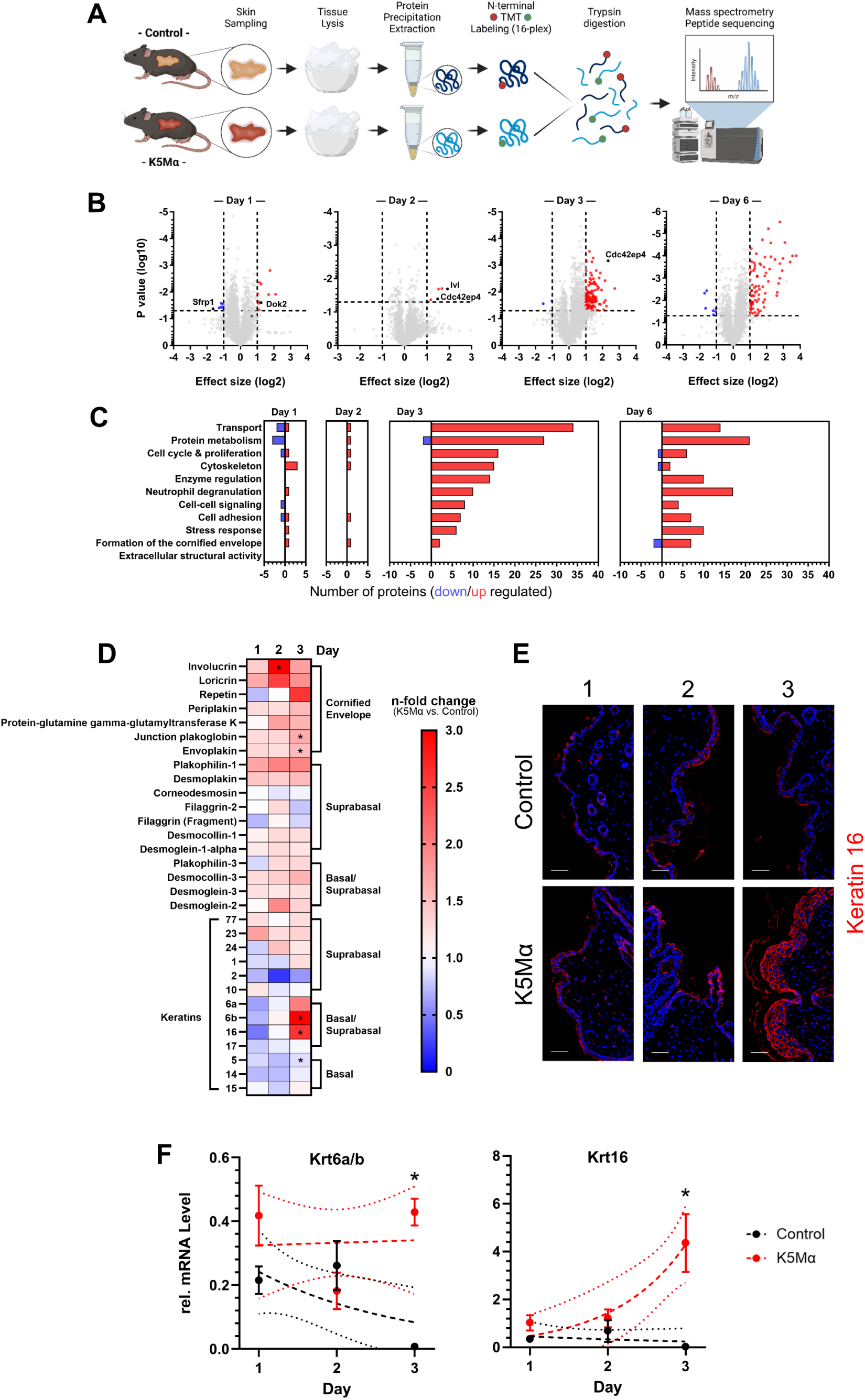
Keratinocyte hyperproliferation in the skin of K5Mα is accompanied by drastic alterations in the skin proteome indicating the initiation of epidermal barrier alarm mechanisms. **(A)** Schematic illustration of the sample preparation work flow for mass spectrometry-based proteomics analysis. Skin samples from control (n=3) and K5Mα (n=3) mice were excised after euthanasia on day one to six after topical TAM treatment, pulverized in a liquid nitrogen-cooled mortar and proteins isolated by methanol/chloroform precipitation. Proteins were TMT-labeled, trypsin digested and subjected to LC-MS/MS analysis. **(B)** Volcano plots display all detected proteins by LC-MS/MS analysis in skin samples of K5Mα mice compared to control mice on day one, two, three and six after TAM treatment. Dashed lines (- - -) mark 0.5- and 2.0-fold changes in effect size (x-axis; log2 transformed) and a p-value of 0.05 (y-axis; log10 transformed), the threshold for statistical significance. Proteins significantly higher (effect size ≥ 2.0-fold; red dots) or lower (effect size ≤ 0.5-fold; blue dots) abundant in the skin of K5Mα compared to control mice are highlighted. Other proteins are depicted as grey dots. **(C)** Bar charts display the number of significantly higher (red bars) and lower (blue bars) abundant proteins detected by LC-MS/MS in the skin of K5Mα compared to control mice on day one, two, three and six after TAM treatment grouped by gene ontology annotations. **(D)** Heatmap displays mean n-fold changes in the level of proteins associated with keratinocyte differentiation status detected by LC-MS/MS in the skin of K5Mα mice compared to control mice on day one, two and three after TAM treatment. Proteins higher (red) and lower (blue) abundant in the skin of K5Mα than in control mice are highlighted by color gradients. **(E)** Representative immunofluorescence images of skin sections from control and K5Mα mice on day one to three after topical TAM treatment. Keratin 16 (red) was detected by indirect immunofluorescence staining. Nuclei were stained with 4’,6-diamino-2-phenylindol (DAPI). Scale bars = 50 µm. **(F)** Relative mRNA levels of *Krt6a/b* and *Krt16* in skin samples from control (n=5 per day) and K5Mα (n=5 per day) mice at days one to three after TAM treatment. Relative messenger RNA levels were normalized to *Gapdh* levels in the respective sample by 2^-ΔΔCt^-method. Data points and error bars display mean values and respective standard errors of the mean (SEM). Results from simple linear regression and non-linear regression based on an exponential growth equation (dashed lines) as well as respective 95% confidence band of the best-fit line (dotted lines) are displayed. Statistically significant differences (p < 0.05) are labeled by an asterisk (*). Graphical illustration in **(A)** was created with BioRender.com.

Docking protein 2 (2.36-fold, p=0.03) and secreted frizzle-like protein 1 (0.32-fold, p=0.04) were of particular interest, since similarly altered levels of both proteins have been reported in atopic dermatitis (AD) and psoriatic lesion. *DOK2* has been identified as a susceptibility gene for AD in a genome-wide association study and proposed as a central hub for signaling processes via CD200 receptor 1 and interleukin-6 receptor followed by downstream activation of signal transducer and activator of transcription 3 (Stat3)^22^. In contrast, *SFRP1* encodes for a negative regulator of wnt-signaling and lower transcript levels were identified in psoriatic lesions^23^. On day two after TAM treatment we identified significantly upregulated levels of involucrin (3.79-fold, p=0.02), a scaffolding protein of the cornified envelope, and Cdc42 effector protein 4, a small GTPase involved in the regulation of epithelial cell polarization, proliferation and migration via interaction with Cdc42^24,25^ (**Figure 4 B; Supplementary Figure 3 B**). Notably, several studies reported a correlation of CDC42 levels with psoriasis severity as well as treatment response^26–28^ and Cdc42 levels were increased in the skin of K5Mα mice already on day one after TAM treatment (1.62-fold, p=0.08). On day three after TAM treatment we detected massive changes in the skin proteome of K5Mα compared to control mice (**Figure 4 B**). In total 130 proteins were significantly upregulated (≥ 2-fold) but only two proteins were significantly downregulated (≤ 0.5-fold). Gene ontology analysis revealed that most of the upregulated proteins were associated with transport processes (n=34) and protein metabolism (n=27) (**Figure 4 C**). Moreover, we also identified 16 significantly upregulated proteins associated with regulation of the cell cycle and proliferation, 15 proteins associated with regulation of the cytoskeleton and 14 proteins associated with the regulation of enzymes (**Figure 4 C**). In Supplementary Figure 3 C, the top 20 significantly higher abundant proteins as well as the two significantly lower abundant proteins are listed (**Supplementary Figure 3 C**). In the following three days until day six after TAM treatment, we still observed drastic differences in the skin proteome of K5Mα mice compared to respective control mice, but at the same time gene ontology analyses indicated changes in the functional focus. Most prominently, over 30% of the 98 significantly higher abundant proteins were associated with neutrophil degranulation (n=17), cellular stress response (n=10) and cell-cell signaling processes (n=4) (**Figure 4 C**). Taken together, our histological findings and proteomics data suggest that the K5Mα skin phenotype develops stepwise and the molecular mechanisms driving its initiation take place within the first three days after TAM treatment. Therefore, our subsequent analyses focused on the time period between day one and day three.

As a first step, we filtered our proteomics data for proteins associated with keratinocyte differentiation status and compared the levels detected in the skin of K5Mα and control mice (**Figure 4 D**). Proteomics data revealed a strong increase in the levels of proteins associated with the formation of the cornified envelope from day one to three. Involucrin (3.79-fold, p=0.02) and loricrin (2.49-fold, p=0.33) were particularly higher abundant on day two, while envoplakin (1.53-fold, p=0.02), junction plakoglobin (1.65-fold, p=0.03) and repetin (2.58-fold, p=0.38) were markedly higher abundant on day three in the skin of K5Mα compared to control mice (**Figure 4 D**). Overall, the levels of most suprabasal adhesion proteins were slightly higher, while the levels of basal and most suprabasal keratins were slightly lower in the skin of K5Mα than control mice. The drastic changes observed in the global proteome in the transition from day two to three are reflected by a significant increase of keratin 16 (2.58-fold, p=0.02) (**Figure 4 D**+**E**) and 6b (3.66-fold, p=0.002) as well as a markedly increase of keratin 6a (1.99-fold, p=0.08) and a slight but significantly decrease of keratin 5 (0.86-fold, p=0.02) levels in the skin of K5Mα mice (**Figure 4 D**). Subsequent qPCR analyses indicated that the observed differences were based on an elevated expression of *Krt6a*, *Krt6b* and *Krt16* (**Figure 4 F**). Notably, upregulation of *Krt6a*, *Krt6b* and *Krt16* expression by keratinocytes has been reported as a hallmark indicating disruption of the barrier integrity, e.g. after wounding^29,30^, associated with induction of hyperproliferation, alterations in adhesion and activation of innate immune responses^31,32^. Of note, upregulation of *KRT6* and *KRT16* expression has also been reported in psoriatic lesions^31,33,34^. In contrast, mRNA levels of other keratins, except for *Krt5*, as well as *Loricrin* and *Involucrin* mRNA levels (**Supplementary Figure 3 D**) did not correlate with the difference in respective protein levels observed between K5Mα and control mice (**Figure 4 F**).

In summary, our proteomics data emphasize that the pathological skin phenotype overserved in K5Mα mice develops stepwise and is initiated by keratinocyte hyperproliferation rapidly triggered within the first three days after induction of meprin α overexpression.

### Meprin α overexpression leads to the induction of a severe innate immune response characterized by the recruitment of neutrophilic granulocytes

The increasing levels of proteins associated with neutrophil degranulation identified in our proteomics data (**Supplementary Figure 3 C**) indicated that epidermal hyperproliferation was accompanied by an intensifying inflammatory response (**Figure 4 C**). In line, histological analyses of K5Mα skin sections suggested infiltration of immune cells into the dermis under hyperplastic epidermal areas. Moreover, we observed prominently enlarged vessels in the skin of K5Mα, indicative for an inflammation-associated activation of blood vessels (**Figure 5 A**). Detection of meprin α and CD45 validated a strong infiltration of leukocytes already on day two after TAM treatment, particularly into the dermis of skin areas characterized by prominent meprin α levels (**Figure 5 B**, white arrowheads). In order to further characterize the leukocyte infiltrate, we filtered our proteomics data for leukocyte population marker proteins. Thereby, we identified significantly elevated levels of CD45 (1.82-fold, p=0.015) and CD11b (1.70-fold, p=0.008) as well as markedly increased levels of Ly6c (1.40-fold, p=0.14) in the skin of K5Mα mice on day three after TAM treatment (**Figure 5 C**). Since the levels of macrophage-(F4/80; Adhesion G protein-coupled receptor E1) and langerhans cell-associated marker proteins (CD207, C-type lectin domain family 4 member K) showed only minor differences between K5Mα and control mice (**Figure 5 C**), we assumed that neutrophilic granulocytes represented the major leukocyte population within the dermal leukocyte infiltrates.

**Figure 5:**
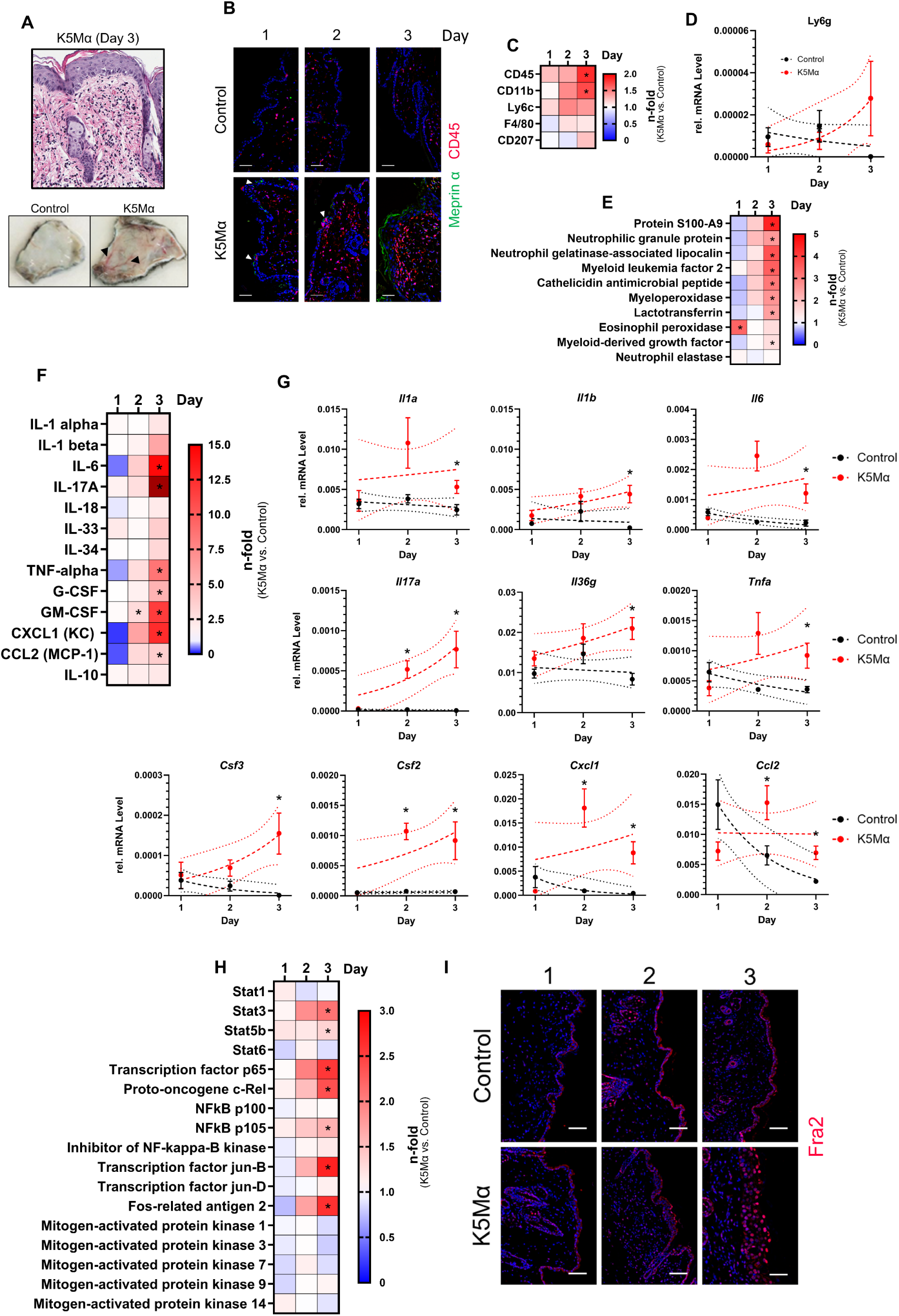
Meprin α overexpression leads to the induction of a severe innate immune response characterized by the recruitment of neutrophilic granulocytes in the skin of K5Mα mice. **(A)** Representative microscopical images of hematoxylin and eosin stained skin section from a K5Mα mouse showing dermal infiltrate as well as photos from the dermal side of the skin. Black arrowheads (◄) mark enlarged inflammatory vessels in the skin of a K5Mα mouse. **(B)** Representative immunofluorescence images of skin sections from control and K5Mα mice on day one to three after topical TAM treatment. Meprin α (green) and CD45 (red) were detected by indirect immunofluorescence staining. Nuclei were stained with 4’,6-diamino-2-phenylindol (DAPI). Scale bars = 50 µm. **(C)** Heatmap displays mean n-fold changes in the levels of leukocyte lineage marker proteins detected by LC- MS/MS in the skin of K5Mα mice compared to control mice on day one, two and three after TAM treatment. Proteins higher (red) and lower (blue) abundant in the skin of K5Mα than in control mice are highlighted by color gradients. **(D)** Relative mRNA levels of *Ly6g* in skin samples from control (n=5 per day) and K5Mα (n=5 per day) mice at days one to three after TAM treatment. *Ly6g* levels were normalized to *Gapdh* levels in the respective sample by 2^-ΔΔCt^-method. Data points and error bars display mean values and respective standard errors of the mean (SEM). Results from non-linear regression based on an exponential growth equation (dashed lines) and respective 95% confidence band of the best-fit line (dotted lines) are displayed. **(E)** Heatmap displays mean n-fold changes in the levels of leukocyte-associated effector proteins detected by LC-MS/MS in the skin of K5Mα mice compared to control mice on day one, two and three after TAM treatment. Proteins higher (red) and lower (blue) abundant in the skin of K5Mα than in control mice are highlighted by color gradients. **(F)** Heatmap displays mean n-fold changes in the level of cytokines and chemokines detected by flow cytometry-based multiplex in the skin of K5Mα mice compared to control mice on day one, two and three after TAM treatment. Proteins higher (red) and lower (blue) abundant in the skin of K5Mα than in control mice are highlighted by color gradients. **(G)** Relative mRNA levels of *Il1a*, *Il1b*, *Il6*, *Il17a*, *Il36g*, *Tnfa*, *Csf3*, *Csf2*, *Cxcl1* and *Ccl2* in skin samples from control (n=5 per day) and K5Mα (n=5 per day) mice at day one to three after TAM treatment. Relative messenger RNA levels were normalized to *Gapdh* levels in the respective sample by 2^-ΔΔCt^-method. Data points and error bars display mean values and respective SEM. Results from simple linear regression and non-linear regression based on an exponential growth equation (dashed lines) as well as respective 95% confidence band of the best-fit line (dotted lines) are displayed. **(H)** Heatmap displays mean n-fold changes in the level of proteins involved in intracellular signal transduction of inflammatory cytokines detected by flow cytometry-based multiplex in the skin of K5Mα mice compared to control mice on day one, two and three after TAM treatment. Proteins higher (red) and lower (blue) abundant in the skin of K5Mα than in control mice are highlighted by color gradients. **(I)** Representative immunofluorescence images of skin sections from control and K5Mα mice on day one to three after topical TAM treatment. Fos-related antigen 2 (Fra2; red) was detected by indirect immunofluorescence staining. Nuclei were stained with DAPI. Scale bars = 50 µm. Statistically significant differences (p < 0.05) are labeled by an asterisk (*).

Indeed, we detected markedly increasing *Ly6g* expression in the skin of K5Mα but not control mice (**Figure 5 D**) as well as significantly increasing levels of neutrophilic granulocyte-specific effector and regulator proteins, like S100A9, neutrophilic granule protein, neutrophil gelatinase-associated lipocalin, myeloid leukemia factor 2, cathelicidin antimicrobial peptide, myeloperoxidase and lactotransferrin (**Figure 5 E**). Notably, high S100A9 levels were detected in infiltrating leukocytes as well as keratinocytes (**Supplementary Figure 4 A**) and expression of both *S100a9* and *S100a8*, important mediators also in psoriasis^35^, was significantly elevated in the skin of K5Mα mice (**Supplementary Figure 4 B**). Supporting these data, induction of epidermal meprin α overexpression in K5Mα mice by tamoxifen-supplemented food led to a systemic leukocytosis characterized by a massive increase of neutrophilic granulocytes in the blood that correlates with the skin phenotype severity (data not shown).

Hyperproliferative, inflammatory skin diseases like psoriasis are driven by the excessive release of inflammatory cytokines like IL-17A and TNFα (reviewed in reference^36^). Hence, we asked whether these cytokines and chemokines might also play a role in the development of the pathological skin phenotype observed in K5Mα mice. Indeed, we detected increasing and significantly higher levels of IL-6 (14.4-fold (day 3)), IL-17A (22.0-fold (day 3)) and TNFα (8.5-fold (day 3)) in the skin of K5Mα compared to control mice (**Figure 5 F**, **Supplementary Figure 4 C**). Moreover, the levels of G-CSF (5.1-fold (day 3)) and GM-CSF (11.6-fold (day 3)), which specifically promote growth and differentiation of neutrophilic granulocytes, as well as CXCL1 (15.5-fold (day 3)) and CCL2 (3.6-fold (day 3)), chemokines that recruit and activate neutrophilic granulocytes, gradually and significantly increased in the skin of K5Mα but not control mice (**Figure 5 F**, **Supplementary Figure 4 C**). Additionally, we validated the increased expression of *Il1a*, *Il1b*, *Il6*, *Il17a*, *Il36g*, *Tnfa*, *Csf3*, *Csf2*, *Cxcl1* and *Ccl2* in the skin of K5Mα by qPCR analyses (**Figure 5 G**). Next, we filtered our proteomics data for key proteins that mediate intracellular signal transduction of these cytokines to examine whether there were temporal changes in the activity of signaling pathways and figure out which signaling pathways dominate during progression of the skin phenotype observed in K5Mα mice. In line with the changes detected for the cytokine and chemokine levels, our analyses revealed significantly higher levels of Stat3 (2.09-fold, p=0.02), Stat5b (1.33-fold, p=0.03), transcription factor p65 (2.56-fold, p=0.009), proto-oncogene c-Rel (2.37-fold, p=0.004), NFκB p105 (1.60-fold, p=0.01) transcription factor jun-B (2.73-fold, p=0.00005) and fos-related antigen 2 (2.61-fold, p=0.04) in the skin of K5Mα compared to control mice on day three after TAM treatment (**Figure 5 H**).

Detection of fos-related antigen 2 (Fra2) revealed a very high proportion of keratinocytes strongly positive for nuclear Fra2 in the skin of K5Mα mice on day three (**Figure 5 I**). Moreover, western blot analyses validated increasing levels of Stat3 phosphorylated on Y705, as an indicator of Stat3 activation, but not for p65/p-p65 (S536) (**Supplementary Figure 4 D**), Stat5/p-Stat5 (Y694) (data not shown) or IκBα/p-IκBα (S32) (data not shown). Overall, we did not find evidence that an adaptive immune response contributes to the skin phenotype development in K5Mα mice, neither by filtering our proteomics data nor by gene expression analyses or detection of lymphocytes and lymphocyte-related proteins in skin lysates or blood (data not shown). However, we did find evidence for the activation of the acute-phase system (**Supplementary Figure 4 E+F**), a finding that might be mechanistically linked to the elevated levels of S100A8 and S100A9, which have been shown to regulate the expression of complement factor C3 in PV^37^.

In summary, our data highlight that keratinocyte hyperproliferation initiated upon induction of meprin α overexpression in the skin of K5Mα mice is rapidly followed by activation of an innate immune response characterized by recruitment and activation of neutrophilic granulocytes. The composition of cytokines, chemokines and other effector molecules released within this inflammatory response strongly resembles the cytokine patterns reported in psoriatic lesions and we suggest that this innate immune response is a major driver for progression of the skin phenotype observed in K5Mα mice.

### Dominant chymotryptic activity accompanies massive alterations within the skin N-terminome of K5Mα mice

Since meprin α is part of the epidermal proteolytic network, we hypothesized that the development of the pathological skin phenotype in K5Mα mice is associated with dysregulation of epidermal protease activity. In order to characterize alterations in the epidermal degradome of skin samples on day one to three after TAM treatment, we performed Terminal Amine Isotopic Labeling of Substrates (TAILS) analyses, an N-terminomics approach that enables the specific identification and quantification of neo-N-termini generated by proteolytic events^38,39^ (**Figure 6 A**). Already on day one, we identified 60 significantly higher (≥ 2.0-fold) and 30 significantly lower (≤ 0.5-fold) abundant neo N-termini peptides in the skin of K5Mα mice compared to control mice (**Figure 6 B**). Gene ontology analysis revealed that most of the peptides with higher and lower abundance in the skin of K5Mα mice were derived from proteins associated with the cytoskeleton (41 *vs*. 6), protein metabolism (16 *vs*. 9) and stress response (14 *vs*. 10) (**Figure 6 B**). On day two after TAM treatment, we detected 25 significantly higher (≥ 2.0-fold) and only two significantly lower (≤ 0.5-fold) abundant peptides in the skin of K5Mα mice (**Figure 6 B**).

**Figure 6:**
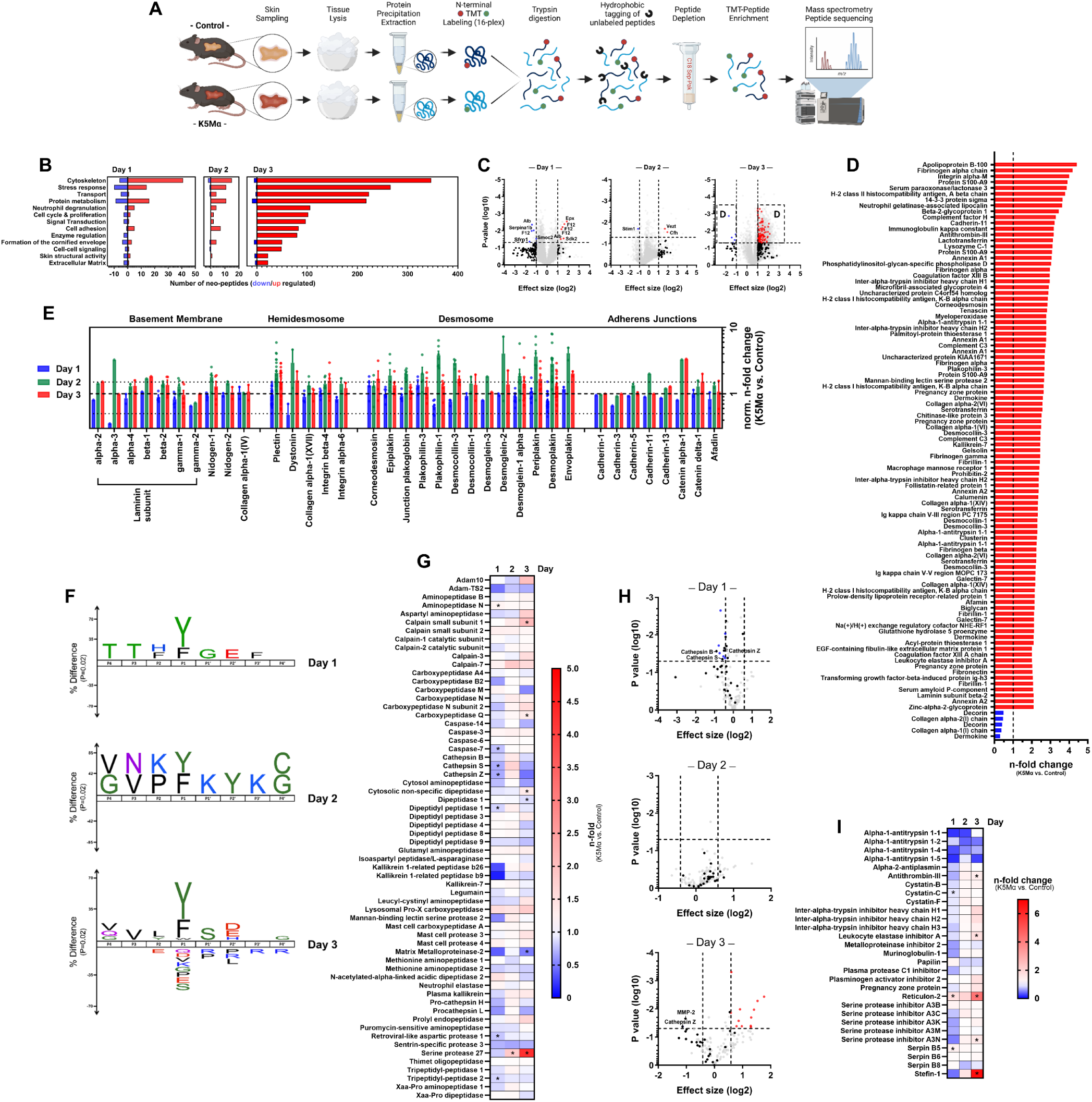
Dominant chymotryptic activity accompanies massive alterations within the skin N-terminome of K5Mα mice. **(A)** Schematic illustration of the sample preparation work flow for Terminal Amine Isotopic Labeling of Substrates (TAILS) analysis. Skin samples from control (n=3) and K5Mα (n=3) mice were excised after euthanasia on day one to six after topical TAM treatment, grinded in a liquid nitrogen-cooled mortar and proteins isolated by methanol/chloroform precipitation. Proteins were TMT-labeled and trypsin digested. Peptides with neo-N termini generated by trypsin digestion were undecanal-labeled, depleted by a C18 Sep-Pak column and enriched TMT-labeled peptides were subjected to LC-MS/MS analysis. **(B)** Bar charts display the number of significantly (p < 0.05) higher (red bars; ≥ 2.0-fold) and lower (blue bars; ≤ 0.5-fold) abundant peptides detected by TAILS in the skin of K5Mα compared to control mice on day one, two and three after TAM treatment grouped by gene ontology annotations. **(C)** Volcano plots display all N-terminally TMT-labeled peptides detected by TAILS analysis in skin samples of K5Mα mice compared to control mice on day one, two and three after TAM treatment. Dashed lines (- - -) mark thresholds for 0.5- and 2.0- fold changes in effect size (x-axis; log2 transformed) and a p-value of 0.05 (y-axis; log10 transformed), the threshold for statistical significance. Peptides derived from extracellular and transmembrane proteins significantly higher (effect size ≥ 2.0-fold; red dots) or lower (effect size ≤ 0.5-fold; blue dots) abundant in the skin of K5Mα compared to control mice are highlighted. Remaining peptides derived from extracellular and transmembrane proteins are depicted as black dots and peptides derived from intracellular proteins are depicted as grey dots. **(D)** Bar chart displays mean n-fold changes (sorted by highest to lowest) in the levels of significantly higher (red bars; ≥ 2.0-fold) and lower (blue bars; ≤ 0.5-fold) abundant peptides (annotated by corresponding proteins) derived from transmembrane and extracellular proteins detected by TAILS in the skin of K5Mα compared to control mice. **(E)** Bar chart displays mean n-fold changes in the levels of peptides (annotated by corresponding proteins) derived from basement membrane, hemidesmosomal, desmosomal and adherens junction proteins detected by TAILS on day one (blue), two (green) and three (red) after TAM treatment in the skin of K5Mα compared to control mice. Peptide levels were normalized to the total level of corresponding proteins detected by proteomics analysis. Equal normalized n-fold levels (n-fold = 1) are highlighted by a dashed line (- - -) and thresholds for 0.5-fold and 1.5-fold altered levels are marked by dotted line (· · ·). **(F)** Sequence logos display favored and disfavored amino acids in position P4 to P4’ of significantly higher (≥ 1.5-fold) abundant peptides derived from transmembrane and extracellular proteins detected by TAILS in the skin of K5Mα compared to control mice on day one, two and three after topical TAM treatment. Amino acids are displayed in one-letter code. Size of the letters indicate relative difference compared to the abundance in a precompiled Swiss-Prot data set comprising proteins from *mus musculus castaneus* at a significance level of p = 0.02. Amino acids were colored according to their chemical properties: polar (green/violet), basic (blue), acidic (red), hydrophobic (black). **(G)** Heatmap displays mean n-fold changes in the level of proteases and peptidases detected by LC-MS/MS in the skin of K5Mα mice compared to control mice on day one, two and three after TAM treatment. Proteins higher (red) and lower (blue) abundant in the skin of K5Mα than in control mice are highlighted by color gradients. **(H)** Volcano plots display N-terminally TMT-labeled peptides derived from proteases and peptidase detected by TAILS analysis in skin samples of K5Mα mice compared to control mice on day one, two and three after TAM treatment. Dashed lines (- - -) mark thresholds for 0.5- and 2.0-fold changes in effect size (x-axis; log2 transformed) and a p-value of 0.05 (y-axis; log10 transformed), the threshold for statistical significance. Peptides significantly higher (effect size ≥ 1.5-fold; red dots) or lower (effect size ≤ 0.75-fold; blue dots) abundant in the skin of K5Mα compared to control mice are highlighted. Peptides potentially generated by release of an N-terminal propeptide, indicating proteolytic activation of the respective protease or peptidase, are depicted as black dots and other peptides are depicted as grey dots. **(I)** Heatmap displays mean n-fold changes in the level of protease inhibitors detected by proteomics in the skin of K5Mα mice compared to control mice on day one, two and three after TAM treatment. Proteins higher (red) and lower (blue) abundant in the skin of K5Mα than in control mice are highlighted by color gradients. Statistically significant differences (p < 0.05) are labeled by an asterisk (*). Graphical illustration in **(A)** was created with BioRender.com.

The majority of neo N-termini peptides with higher abundance in the skin of K5Mα mice were derived from proteins associated with the cytoskeleton (15), stress response (11), protein metabolism (11) and cell adhesion (7) (**Figure 6 B**). As in our total proteome data, we observed massive changes in the skin N-terminome of K5Mα mice in the transition from day two to three after TAM treatment. In total, we identified 873 peptides with significantly higher abundance (≥ 2.0-fold), while only 18 peptides were significantly lower abundant (≤ 0.5-fold) in the skin of K5Mα compared to control mice (**Figure 6 B**). Gene ontology analyses showed that most peptides with significantly higher abundance in the skin of K5Mα mice originated from proteins associated with the cytoskeleton (347), stress response (266), transport processes (223) and protein metabolism (218) (**Figure 6 B**). Overall, these drastic differences in the skin N-terminome of K5Mα and control mice support the assumption that meprin α overexpression causes a dysregulation of the protease network characterized by an increased proteolytic activity.

Meprin α is either secreted to the extracellular space or forms a heterodimer complex with membrane-bound meprin β^40^. In both cases, meprin α cleaves its substrates extracellularly and intracellular activity of meprin α has not been reported so far. Therefore, we filtered our TAILS data in a first step for peptides with higher (≥ 2.0-fold) and lower (≤ 0.5-fold) abundance in the skin of K5Mα mice. In the second step, we identified by UniProt database search within this pool of peptides those that potentially could be directly generated by extracellular protease activity and categorized them into not significantly higher/lower abundant (**Figure 6 C**, black dots), significantly lower abundant (**Figure 6 C**, blue dots) and significantly higher abundant (**Figure 6 C**, red dots) in the skin of K5Mα mice. Applying these filter criteria, we identified on day one after TAM treatment five significantly lower and six significantly higher abundant peptides in the skin of K5Mα mice. (**Figure 6 C**). However, none of these peptides were significantly altered in the skin of K5Mα mice on day two after TAM treatment. (**Figure 6 C**). On day three after TAM treatment, we detected in total 94 significantly higher and five significantly lower abundant peptides that might be generated by extracellular proteolytic activity in the skin of K5Mα mice (**Figure 6 C**+**D**). Overall, the 94 enriched peptides were derived from proteins of various functions, but a major part was related to innate immune and acute phase response, as expected from our previous findings. Another large proportion of highly abundant peptides originated from cell adhesion and matrix proteins. Modulation of keratinocyte adhesion to the basement membrane and neighboring keratinocytes via proteolytic processing of hemidesmosomes, desmosomes and adherens junctions is a central regulatory mechanism for keratinocyte proliferation and differentiation^3,41,42^.

In order to examine if aberrant proteolytic processing of keratinocyte adhesion proteins takes place within the development of the skin phenotype in K5Mα mice, we filtered our TAILS data for peptides derived from basement membrane, hemidesmosomal, desmosomal and adherens junction-associated proteins (**Figure 6 E**). In our global proteomics data, we detected for most of these adhesion proteins higher levels in the skin of K5Mα compared to control mice, a finding that is very likely attributed to the epidermal hyperplasia. However, even under homeostatic conditions there is a constant turnover of these adhesion proteins within the epidermis. Therefore, higher baseline levels of these adhesion proteins would consequently be associated with higher peptide levels by homeostatic and not mandatory by pathologically increased turnover. Since our proteomics data were generated from the same sample set as our TAILS data, we normalized the signal intensity of each peptide from our TAILS data to the signal intensity of the respective protein from our proteomics data to compensate for potential effects by the homeostatic turnover. On day one after TAM treatment we did not observe overall higher levels of peptides derived from basement membrane, hemidesmosomal, desmosomal and adherens junction proteins in the skin of K5Mα mice but markedly lower levels of peptides originating from laminin subunit alpha-3 and dystonin (**Figure 6 E**). Interestingly, both play an important role in wound healing^43,44^ and, therefore, the observed alterations might be associated with elevated keratinocyte proliferation. Most prominent changes were detected on day two after TAM treatment. In the group of basement membrane associated proteins, we identified markedly higher peptide levels (≥ 1.5-fold) derived from laminin subunit alpha-3, beta-1, beta-2 and gamma-1 as well as nidogen-1 and -2 in the skin of K5Mα mice (**Figure 6 E**). Moreover, our data revealed clearly higher levels of peptides originating from all detected hemidesmosomal and desmosomal proteins as well as from adherens junction proteins cadherin-11 and catenin alpha-1 in the skin of K5Mα mice (**Figure 6 E**). Terminal differentiation of keratinocytes is accompanied by extensive proteolytic processing of desmosomal proteins (reviewed in reference^45^), therefore, our N-terminomics data match with hyperkeratosis observed in K5Mα mice. On day three further increased levels were detected for peptides derived from laminin subunit beta-1, integrin beta-4, corneodesmosin and catenin delta-1. In summary, these findings indicate a pathologically increased turnover of particular proteins in the basement membrane and adherens junctions as well as almost all detected proteins of the hemidesmosomal and desmosomal compartment on day two after TAM treatment in the skin of K5Mα mice concomitant to the initiation of keratinocyte hyperproliferation.

In order to characterize the extracellular proteolytic activity in the skin of K5Mα mice in more detail, we generated sequence logos for positions P4 to P4’ of the peptides potentially generated by extracellular proteolytic activity and identified as significantly higher abundant (≥ 1.5-fold, p < 0.05) in the skin of K5Mα compared to control mice. Sequence logos indicated a dominant chymotryptic activity in the skin of K5Mα mice that augmented from day one to three after TAM treatment (**Figure 6 F**). Interestingly, we also found a clear preference for aspartate and glutamate in P2’ position, which fits to the cleavage preference of meprin α^8^. This preference was still observed in the sequence logo after removal of all potential chymotryptic peptides from our experimental data set and re-analysis of the remaining peptides (data not shown). Filtering our global proteomics data set for proteases and peptidases, we found significantly altered levels for several proteases (**Figure 6 G**). However, only a few proteases gradually increased from day one to three in the skin of K5Mα mice in comparison to control mice. For example, we identified significantly higher levels of calpain small subunit-1 (2.06-fold, p=0.02), carboxypeptidase Q (1.30-fold, p=0.03) and cytosolic non-specific dipeptidase (1.30-fold, p=0.02) (**Figure 6 G**). The most prominently altered levels were observed for serine protease 27, also known as marapsin, a protease that is found at highly elevated levels in wound healing and psoriasis^46^. Serine protease 27 levels were already significantly higher on day two (1.82-fold, p=0.001) and further increased on day three (4.26-fold, p=0.02) in the skin of K5Mα mice (**Figure 6 G**). In line, we detected significantly increasing mRNA levels of *Prss27* in the skin of K5Mα but not control mice (**Supplementary Figure 5 A**). Notably, elevated *Prss27* expression has been reported in dependence of JAK1 activation and, therefore, might be related to increased Stat3 activation (**Supplementary Figure 4 D**). Since for serine protease 27 a tryptic cleavage specificity is reported^47^ and many proteases in the skin are expressed as zymogens, we assumed that the dominant chymotryptic activity in the skin of K5Mα might be related to the activation of chymotryptic proteases. Therefore, we filtered our TAILS data set for peptides derived from proteases and identified by database search those N-termini that indicate previous proteolytic activation by release of an N-terminal propeptide. However, at none of the three sample times we found direct evidence for an increased activation of any protease in the skin of K5Mα mice (**Figure 6 H**) even after normalization of our TAILS data set by our global proteomics data (**Supplementary Figure 5 B-D**). Nevertheless, on day three after TAM treatment, we found ten significantly higher abundant protease derived peptides in the skin of K5Mα mice (**Supplementary Figure 5 D**).

Although none of these peptides comprised an activation-related N-terminus, we found strong evidence for an elevated kallikrein-7 turnover (Kallikrein-7 [Y.S101] 2.59-fold, p=0.04) (**Supplementary Figure 5 D**) in the skin of K5Mα mice, a protease with chymotryptic cleavage preference^48,49^ that plays a central role in terminal differentiation^7^ and inflammatory skin diseases^50,51^. Supporting the assumption that kallikrein-7 activity causes the dominant chymotryptic cleavage profile in the skin of K5Mα mice, we detected significantly higher levels of Klk7 mRNA in the skin of K5Mα on day three after TAM treatment (**Supplementary Figure 5 A**) as well as significantly higher abundance of peptides derived from cathelicidin, an antimicrobial peptide whose proteolytic cleavage has been correlated with increased kallikrein-7 activity in acne rosacea^52^. Additionally, we found significantly increasing Klk8 mRNA levels in the skin of K5Mα but not control mice (**Supplementary Figure 5 A**).

Besides the proteolytic activation of proteases, inhibitors are the main regulators of the proteolytic network in the skin. Therefore, we also filtered our proteomics data for protease inhibitors and relatively quantified differences in the skin of K5Mα compared to control mice. Thereby, we found significantly increased levels of antithrombin-III (1.44-fold, p=0.001), leukocyte elastase inhibitor A (1.71-fold, p=0.0003) and serine protease inhibitor A3N (1.50-fold, p=0.017) (**Figure 6 I**). Most prominent changes were observed for reticulon-2 (4.23-fold, p=0.04), a potential modulator of bace-1 activity, and stefin-1 (6.68-fold, p=0.007), an intracellular thiol protease inhibitor (**Figure 6 I**). Notably, stefin-1 upregulation has already been reported in PV and murine PV models^53^.

In summary, our TAILS data highlight significant changes in the skin N-terminome of K5Mα mice. Our data provide evidence, that initiation of keratinocyte hyperproliferation is accompanied by drastic changes within the compartment of adhesion proteins, which precede the massively altered turnover of extracellular proteins observed on day three after TAM treatment. Moreover, our findings indicate that extracellular proteolytic activity is predominantly mediated by proteases exhibiting a chymotryptic cleavage preference. Within the protease compartment we observed a significant increase of serine protease 27 and markedly elevated processing of kallikrein-7, which might be causative for the alterations within the skin N-terminome observed in K5Mα mice.

### Filtering TAILS data for meprin α-specific selection criteria identified dermokine as a potential substrate

Our TAILS data indicated that keratinocyte hyperproliferation upon the overexpression of meprin α in the skin of our K5Mα mice comes with increased epidermal proteolytic activity characterized by dominant chymotryptic and not meprin α-specific activity. However, a dominant chymotryptic cleavage preference was also observed in sequence logos generated from all detected peptides that were almost equally abundant (≤ 1.1-fold and ≥ 0.9-fold) in the skin of K5Mα and control mice (data not shown), highlighting that prominent chymotryptic activity is also characteristic for the physiological proteolytic network. Therefore, we assumed that the onset of the skin phenotype observed in K5Mα mice is triggered by meprin α-mediated cleavage of a few decisive substrates rather than an overall excessive cleavage of many proteins. In order to identify those potential meprin α substrates, we applied a stepwise filtering scheme to our TAILS data set (**Figure 7 A**+**B**, for detailed description see **Materials and Methods** section). In total, we detected by TAILS 41,619 peptides on day one to three after TAM treatment in skin samples from K5Mα and control mice. Of note, we did not detect any peptide with our settings that was exclusively present in the data set of either control or K5Mα mice, indicating that all identified peptides are generated physiologically and not artificially by meprin α overexpression. Applying our filtering criteria, we identified six peptide candidates on day one, ten peptide candidates on day two and ten peptide candidates on day three, which were derived from ten different proteins (**Figure 7 A**). Substrate candidates included dermokine, syndecan-1, complement C3, aminopeptidase N, mast cell protease 4, fibrillin-1, collagen alpha-1 (XII), complement factor D, death domain-containing membrane protein NRADD and follistatin-related protein 1 (**Figure 7 C**). Subsequent normalization of the peptide candidate levels to the levels of the respective total proteins revealed that only the peptide D_298_EGYSVSR derived from dermokine was markedly higher abundant (2.02-fold, p=0.28) in the skin of K5Mα mice already on day one after TAM treatment (**Figure 7 D**). In order to assess a possible interaction of meprin α with our identified substrate candidates, we modelled each identified substrate candidate in complex with either monomeric or homodimeric meprin α using alphaFold 3^54^. Thereby, we identified a model of homodimeric meprin α in complex with dermokine that would result in cleavage of dermokine by one meprin α subunit between phenylalanine in position 297 and aspartate in position 298 (**Supplementary Figure 6 A**) as well as between arginine in position 439 and alanine in position 440 by the other subunit (**Supplementary Figure 6 B**). While the first putative cleavage site between F297 and D298 would exactly generate the dermokine-related peptide we identified by filtering our TAILS data set, also the second putative cleavage site between R439 and A440 would generate a dermokine-derived peptide we identified by our TAILS analysis ([R].A_440_GGADQFSKPEAR.[Q]; **Figure 8 A**). Therefore, we focused our further analyses on dermokine as a potential epidermal substrate of meprin α.

**Figure 7:**
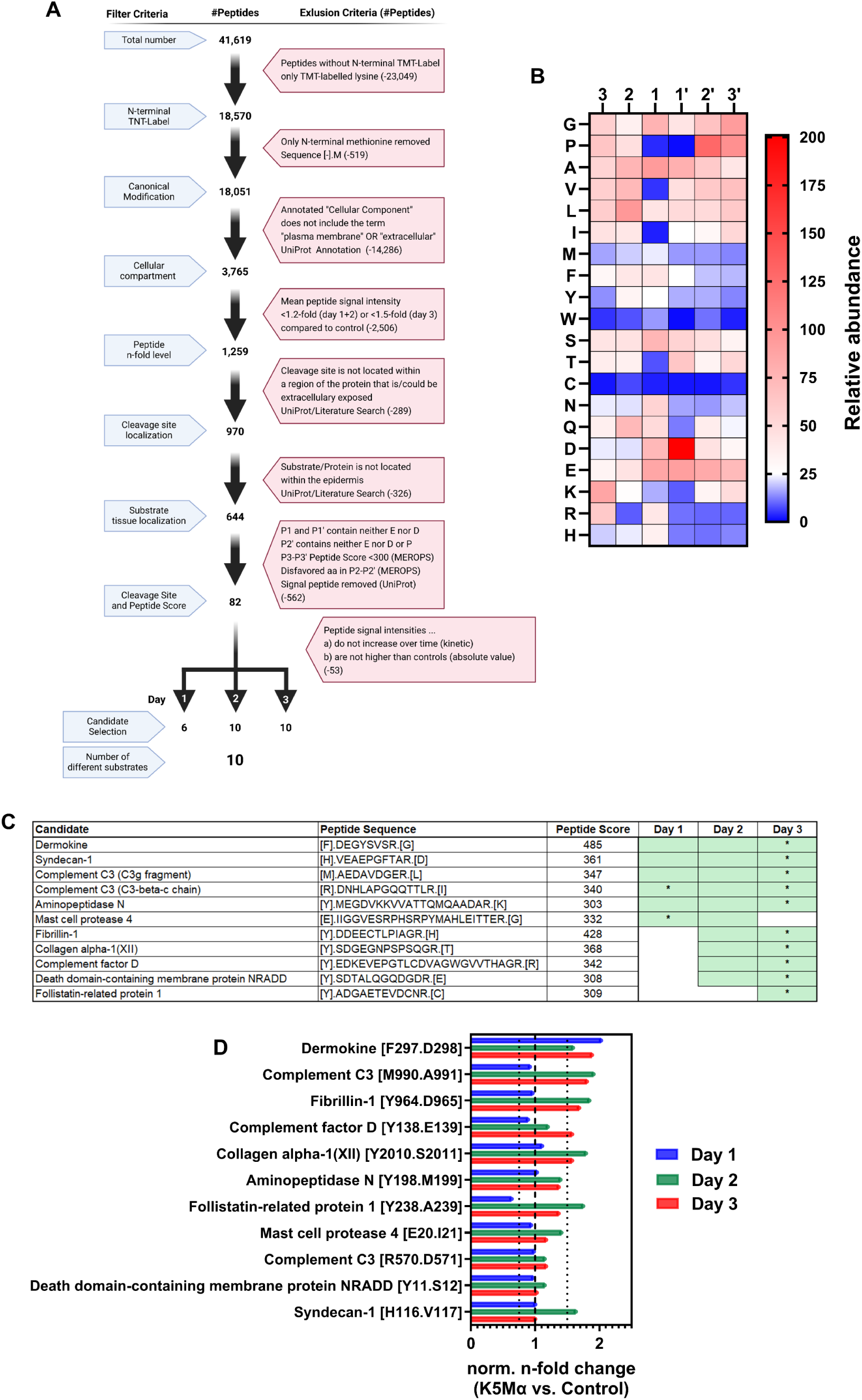
Filtering TAILS data for meprin α-specific selection criteria identified dermokine as a potential substrate. **(A)** Annotation of the filtering and exclusion criteria as well as illustration of the filtering process for identification of peptides potentially generated by meprin α cleavage within the TAILS data set. **(B)** Heatmap displays the amino acids in one-letter code preferred by human meprin α in position P3 to P3’ to the scissile bond. Likelihood scores in the range from 1 (very unlikely, blue) to 201 (very likely, red) were obtained from MEROPS database (https://www.ebi.ac.uk/merops/) and highlighted by color gradient. **(C)** Summary of the potential meprin α substrates identified by filtering of the TAILS data set. Listed are the corresponding proteins (“Candidate”), the respective peptides detected by TAILS in one-letter code and the Peptide Score (sum of likelihood scores for amino acids in position P3 to P3’). Start and end of the peptide sequences are marked by a full stop. Amino acids in squared brackets are annotated based on the protein sequence obtained from UniProt database. Boxes marked in green indicate that the peptide has been identified by filtering of the TAILS data set on the respective day. Statistically significant differences (p < 0.05) between K5Mα and control mice are labeled by an asterisk (*). **(D)** Bar chart displays mean n-fold changes in the levels of peptides (annotated by corresponding proteins and putative cleavage site in squared brackets) potentially generated by meprin α on day one (blue), two (green) and three (red) after TAM treatment in the skin of K5Mα compared to control mice. Peptide levels were normalized to the total level of corresponding proteins detected by proteomics analysis. Equal normalized n-fold levels (n-fold = 1) are highlighted by a dashed line (- - -) and thresholds for 0.75-fold and 1.5-fold altered levels are marked by dotted lines (· · ·). Graphical illustration in **(A)** was created with BioRender.com.

**Figure 8:**
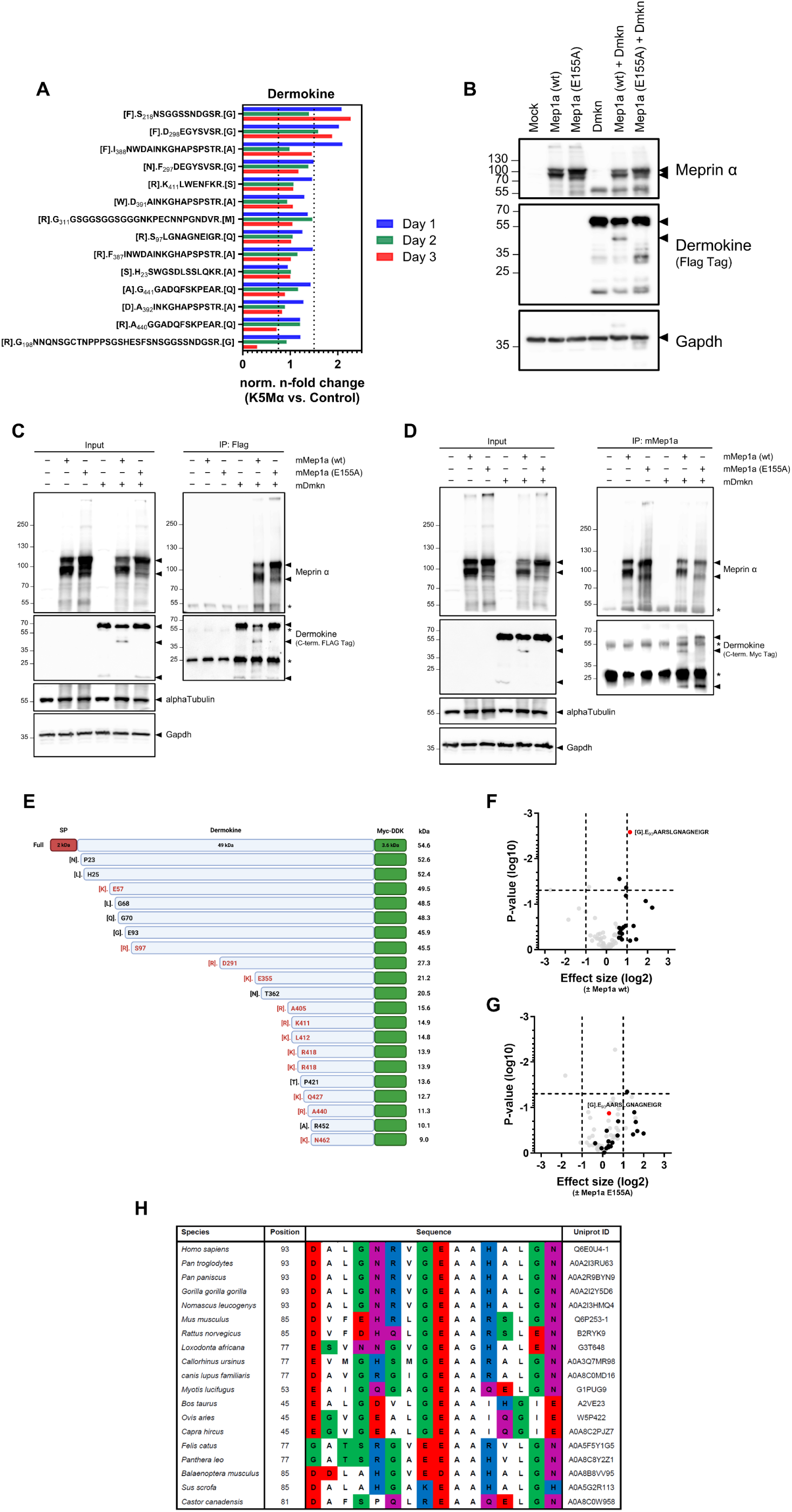
Validation of dermokine cleavage by meprin α. **(A)** Bar chart displays mean n-fold changes in the levels of dermokine derived peptides detected by TAILS on day one (blue), two (green) and three (red) after TAM treatment in the skin of K5Mα compared to control mice. Start and end of the peptide sequences are marked by a full stop. Amino acids in squared brackets are annotated based on the protein sequence obtained from UniProt database. Peptide levels were normalized to the total dermokine level detected by proteomics analysis. Thresholds for 0.75-fold and 1.5-fold altered levels are marked by dotted line (· · ·). **(B)** Western blot detection of meprin α and dermokine (via FLAG Tag) in lysates of HEK293T cells transiently transfected with an empty control plasmid (Mock) or plasmids encoding wildtype (wt) murine meprin α (*Mep1a*), C-terminally Myc-FLAG-tagged murine dermokine (*Dmkn*) and/or a catalytically inactive meprin α variant (*Mep1a* (E155A)). Gapdh was detected as a reference. **(C+D)** Western blot detection of meprin α and dermokine (via Myc or FLAG Tag) in lysates of HEK293T cells transiently transfected with an empty control plasmid or plasmids encoding wildtype (wt) murine meprin α (*mMep1a*), C-terminally Myc-FLAG-tagged murine dermokine (*mDmkn*) and/or a catalytically inactive meprin α variant (*mMep1a* (E155A)) before (Input) and after co- immunoprecipitation (IP) of **(C)** dermokine (IP:Flag) or **(D)** meprin α (IP: mMep1a). Gapdh and alpha Tubulin were detected as references in input lysates. Asterisks (*) mark detected light and heavy chains of the antibodies used for immunoprecipitation **(G)** Schematic illustration of the full-length C-terminally Myc-FLAG (Myc-DDK)-tagged murine dermokine (*Dmkn*) encoded in the plasmid used for overexpression as well as putative C-terminal fragments (CTFs) corresponding to the dermokine derived peptides identified by mass spectrometry after co-immunoprecipitation. Annotated are the signal peptide (SP), the N-terminal amino acids in one-letter codes including position in the protein and the putative size of full-length dermokine and CTFs in kilodalton (kDa). Amino acids in squared brackets are annotated based on the protein sequence obtained from UniProt database. Cleavage sites marked in red are potentially generated by trypsin digestion during sample preparation **(F+G)** Volcano plots display dermokine peptides detected by mass spectrometry in lysates of transiently transfected HEK293T cells after immunoprecipication of dermokine. Dermokine peptide levels detected in immunoprecipitates from HEK293T cells transfected only with a plasmid encoding C-terminally Myc-FLAG-tagged *Dmkn* were compared to levels detected in immunoprecipitates from HEK293T cells co-transfected with **(F)** *Mep1a* wt or **(G)** catalytically inactive *Mep1a* E155A. Dashed lines (- - -) mark thresholds for 0.5- and 2.0-fold changes in effect size (x-axis; log2 transformed) and a p-value of 0.05 (y-axis; log10 transformed), the threshold for statistical significance. Peptides higher abundant (≥ 1.5-fold) in immunoprecipitates from cells co-expressing *Mep1a* wt and *Dmkn* than in cells only expressing *Dmkn* are marked as black dots. The putative dermokine peptide derived by meprin α cleavage is annotated and marked (red dot). All other detected dermokine-derived peptides are presented as grey dots (n = 3 *vs*. 3). **(H)** Alignment of the protein sequences around the identified cleavage site in dermokine from various species. Amino acids were color coded according to their chemical properties: polar (green/violet), basic (blue), acidic (red), hydrophobic (white). Graphical illustration in **(E)** was created with BioRender.com.

### Validation of dermokine as a substrate of meprin α

Dermokine is a secreted protein highly expressed in the spinous layer of the epidermis, identified by Matsui and colleagues in 2004^55^. In total, we identified by TAILS 14 different peptides derived from dermokine isoforms (**Figure 8 A**). In the UniProt database four murine isoforms generated by alternative splicing are listed for dermokine. Thirteen of the identified peptides were covered by isoform 1 (dermokine β) and isoform 4. Eleven peptides were covered by isoform 2 (dermokine γ) and isoform 1. Only one peptide (H_22_SWGSDLSSLQKR) was exclusively covered by isoform 3 (dermokine α), a truncated isoform of about 11.5 kDa. Besides the dermokine peptide we identified by filtering our TAILS data for potential meprin α cleavage sites (D_298_EGYSVSR), we detected two additional dermokine-derived peptides that were markedly higher abundant (≥ 2.0-fold) in the skin of K5Mα than control mice already on day one after TAM treatment (S_218_NSGGSSNDGSR, 2.07-fold, p=0.24; I_388_NWDAINKGHAPSPSTR, 2.09-fold, p=0.36) and four more peptides that were almost 1.5-fold increased (**Figure 8 A**). In contrast, the abundance of two dermokine-derived peptides markedly or even significantly decreased from day one to day three after TAM treatment in the skin of K5Mα mice (G_198_NNQNSGCTNPPPSGSHESFS NSGGSSNDGSR, 0.30-fold, p=0.001; A_440_GGADQFSKPEAR, 0.71-fold, p=0.27) (**Figure 8 A**). These striking differences support our hypothesis of an altered dermokine processing in the skin of K5Mα mice. In line with this assumption, we detected similar *Dmkn* mRNA levels (**Supplementary Figure 6 C**) but markedly lower dermokine protein levels on day one after TAM treatment in the skin of K5Mα mice (**Supplementary Figure 6 D**). In contrast, dermokine levels detected in the skin of control mice were constant from day one to three after TAM treatment (**Supplementary Figure 6 D**).

In order to examine if meprin α is capable of cleaving dermokine, we performed transient co-expression of meprin α and dermokine *in vitro*. For this purpose, we transfected HEK293T cells with constructs encoding wildtype (wt) *Mep1a*, C-terminal Myc-Flag-tagged *Dmkn* and a *Mep1a* variant (E155A) that leads to the synthesis of catalytically inactive meprin α (**Supplementary Figure 6 E**). The *Dmkn* sequence in the plasmid we used encoded dermokine β (UniProt ID: Q6P253-4), which comprises all of the potential cleavage sites identified by TAILS except for the one between S22 and H23, which is exclusively present in dermokine α (UniProt ID: Q6P253-3). Subsequent western blot analyses revealed a prominent C-terminal dermokine fragment with a size of about 45 kDa exclusively present in the lysates of HEK293T cells expressing both meprin α wt and dermokine. Moreover, we observed lower levels of full-length dermokine (∼55 kDa) in these lysates (**Figure 8 B**). We validated the interaction of both meprin α wt and inactive meprin α (E155A) with dermokine by co-immunoprecipitation after pull-down of dermokine via the FLAG-tag (**Figure 8 C**) as well as after pull-down of meprin α (**Figure 8 D**), indicating that the molecular interaction of meprin α with dermokine is independent of its proteolytic activity. Notably, we also detected the C-terminal fragment (CTF) of dermokine after pull-down of meprin α wt, supporting our assumption that this fragment is directly generated by meprin α and not by any other protease, which might be activated directly or indirectly by meprin α (**Figure 8 D**). However, the molecular weight of the CTF detected *in vitro* did not match with the expected molecular weights of the CTFs that would be generated by the three most prominent potential cleavage events identified by TAILS, which would result in CTFs of about 34, 27 and 18 kDa, respectively. The only peptide identified by TAILS that would match with a CTF molecular weight of about 45 kDa was neither altered with regard to its abundance in the skin of K5Mα and control mice nor comprised a potential cleavage site that would fit to the specificity of meprin α ([R]S_97_LGNAGNEIGR.[Q]; (**Figure 8 A**). We hypothesized that meprin α can cleave dermokine at multiple sites and the peptide corresponding to the CTF detected *in vitro* might be rapidly further processed *in vivo* and, therefore, was not detectable by TAILS. In order to identify the meprin α cleavage site that leads to formation of the dermokine CTF observed *in vitro*, we performed a proteomics analysis after co-immunoprecipitation of dermokine from lysates of HEK293T cells transfected only with the plasmid encoding *Dmkn β* or *Dmkn β* in combination with either *Mep1a* wt or *Mep1a* E155A. First of all, we identified both dermokine and meprin α among the highest abundant proteins in the co-immunoprecipitates of co-transfected HEK293T cells, validating the interaction of dermokine and meprin α (**Supplementary Figure 6 F**).

In total, we detected 86 dermokine-derived peptides covering 72% of its total sequence. Among these 86 peptides, we identified 20 peptides that were higher abundant (≥ 1.5-fold) in the immunoprecipitates from HEK293T cells expressing both *Mep1a* wt and *Dmkn* than in those from cells only expressing *Dmkn* (**Figure 8 E**). Of these 20 peptides, we identified 12 peptides that were likely derived by trypsin instead of meprin α cleavage (**Figure 8 E**, red). Additionally, two peptides were likely derived by release of the N-terminal signal peptide. Within the remaining six peptides, only three peptides were identified that could correspond to the N-terminus of a dermokine CTF of about 45 kDa ([L].G68, [Q].G70 and [G].E93; **Figure 8 E**). Here, we identified significantly higher levels of [G].E_93_AARSLGNAGNEIGR in the immunoprecipitates of cells expressing both *Dmkn* and *Mep1a* wt (**Figure 8 F**), but not in cells expressing *Dmkn* and catalytically inactive *Mep1a* E155A (**Figure 8 G**). Therefore, our data suggest that meprin α cleaves dermokine predominantly between G92 and E93.

In order to estimate potential functional consequences of dermokine cleavage by meprin α at this site, we modelled the structure of dermokine using alphaFold 3 (**Supplementary Figure 6 G**). Notably, for the major part of dermokine alphaFold 3 calculated very low (pIDDT <50) and low prediction scores (70 > pIDDT > 50) as well as a very low predicted template modelling score of only about 0.2, which is reflected by a high proportion of unstructured areas in all models. The highest prediction scores were calculated for the dermokine sequence from N22 to S125, which was reproducibly predicted to fold into two alpha helices, with the meprin α cleavage site located in the first alpha helix (**Supplementary Figure 6 G**). Lastly, we aligned the dermokine sequence around the identified meprin α cleavage site from various mammals (**Figure 8 H**). Strikingly, our analysis showed that this region is highly conserved in mammal species indicating that proteolytic regulation of dermokine function by meprin α might represent an evolutionary conserved mechanism.

## Discussion

The expression and spatial separation of meprin α and meprin β in the skin, localized in the *stratum basale* and the *stratum granulosum*, respectively, suggested that both proteases regulate different aspects of epidermal homeostasis^18^. Based on first experiments with HaCaT cells exposed to meprin activity and detection of meprins in psoriatic lesions, it was suggested that meprin β rather regulates terminal keratinocyte differentiation while meprin α might modulate keratinocyte proliferation^18^. Therefore, we hypothesized that if meprin α regulates keratinocyte proliferation then epidermal mislocalization and/or elevated meprin α activity might be a driver of keratinocyte hyperproliferation in skin diseases like PV. In order to test this hypothesis, we generated the K5Mα mouse model, which enables epidermal overexpression of meprin α to mimic pathological meprin α levels observed in PV^18^. Here we report, that K5Mα mice develop a severe inflammatory skin phenotype within only six days after induction of meprin α overexpression initiated by keratinocyte hyperproliferation and identified dermokine cleavage by meprin α as a potential pathological mechanism.

Dermokine was identified and first described in 2004 by Matsui and colleagues as a protein secreted by differentiating keratinocytes in the spinous layer of mouse and human epidermis^55^. Back then, *in situ* hybridization screenings revealed two splice variants of dermokine, designated as alpha and beta. However, subsequent studies indicated that human *DMKN* encodes more than 15 isoforms and more than 18 computationally mapped isoforms^56^. For murine dermokine there are three major secreted isoforms annotated, namely α, β and γ^57^. A global transcriptomics study by Bazzi and colleagues identified *Dmkn* as one of the most highly differently expressed genes during development of the murine epidermis^58^. Attempting to unravel the function of dermokine, Leclerc and colleagues generated *Dmkn* βγ-deficient mice^57^ while Utsonomiya and colleagues generated both *Dmkn* βγ- and *Dmkn* αβγ-deficient mice^59^. Both groups reported a transient scaly skin phenotype in *Dmkn* βγ-deficient mice that disappeared one week after birth. In contrast, *Dmkn* αβγ-deficient mice developed a progressively increasing severe ichthyosis^59^. Based on the findings by Higashi and colleagues who provided evidence that dermokine acts as a negative regulator of ERK signaling^60^, Utsunomiya and colleagues proposed dermokine as a negative regulator of keratinocyte proliferation that balances the differentiation status of keratinocytes^59^. In direct comparison, the skin phenotype we described here for K5Mα mice represents a striking phenocopy of *Dmkn* αβγ-deficient mice reported by Utsunomiya and colleagues. Therefore, we propose that cleavage by meprin α causes the functional inactivation of dermokine leading to keratinocyte hyperproliferation.

Our data strongly support that initiation of the skin phenotype in K5Mα mice is triggered by keratinocyte hyperproliferation. Besides a rapidly increasing proportion of Ki67-positive keratinocytes, we detected markedly higher levels of the proliferating cell nuclear antigen, a protein that is also elevated in malignant and non-malignant hyperproliferative skin diseases^61^, as one of the earliest events in the skin of K5Mα mice followed by the increase of 15 other proteins associated with regulation of the cell cycle and proliferation. Moreover, our histological analyses indicated that keratinocyte hyperproliferation is not restricted to the *stratum basale* but also takes place in suprabasal layers of the skin. Similar findings were reported in the context of wound re-epithelialization^62^ and PV^63^. Supporting a mechanistic link between meprin α, dermokine and keratinocyte proliferation, meprin α is highly abundant particularly in epidermal tongues during wound re-epithelialization^19^. In line, both Leclerc as well as Hasegawa and colleagues reported that dermokine expression is induced in epidermal tongues during early stages of wound healing in mice^57^. However, having a closer look at the dermokine staining in both publications, we noted rather a markedly lower staining intensity particularly at the wound edges compared to epidermal areas more distal to the wound edge. Moreover, Hasegawa and colleagues reported that particularly early wound healing is significantly delayed by topical treatment with recombinant dermokine β^64^. In the context of the hyperproliferative skin disease PV, we reported that meprin α in lesional skin areas is highly abundant in suprabasal layers and not in the basal layer as in healthy skin^18^. Notably, the same epidermal localization was reported for dermokine in PV by Hasegawa and colleagues^65^. Here again, dermokine staining intensity in the representative PV skin section appeared to us markedly lower compared to representative sections from other skin pathologies. However, it has to be further investigated if these spatial correlations between meprin α and dermokine levels particularly in wound healing and PV are due to dermokine cleavage by meprin α. Nevertheless, in case of the mechanistic validation, this would implicate that re-activation of the proliferative potential in keratinocytes having already entered terminal differentiation is both a physiological and pathophysiological process regulated by meprin α.

Similar to *Dmkn* αβγ-deficient mice^59^, our K5Mα mice developed an impaired barrier integrity characterized by an increased epidermal water loss. Transepidermal water evaporation is controlled by the architecture of the *stratum corneum*, where corneocytes and extracellular lamellar lipid membranes form a dense hydrophobic barrier (reviewed by references^66,67^). In the *stratum corneum* of K5Mα mice we noticed a strong accumulation of lipid droplets, a characteristic finding also reported in *Dmkn* αβγ-deficient mice^59^.

We assume that these lipid droplets are derived from incomplete processing of lamellar bodies in the *stratum granulosum* due to the excessive number of keratinocytes transitioning from the *stratum granulosum* into the *stratum corneum*. A compromised release of lipids from lamellar bodies would cause a reduced synthesis of extracellular lamellar lipid membranes and consequently lead to an impaired evaporation barrier. Supporting the hypothesis that K5Mα mice acquire an impaired *stratum corneum* differentiation, we observed incomplete differentiation of corneocytes and significantly increased levels of protein-glutamine gamma-glutamyl transferase 5 (TGM5), an enzyme that is responsible for the release and extracellular crosslinking of contents from lamellar bodies^68^. De Koning and colleagues reported that the levels of TGM5, TGM1 and TGM3 are also upregulated in PV^69^, suggesting a link between elevated transglutaminase levels and an impaired formation of the *stratum corneum*.

With regard to cytoskeletal alterations in keratinocytes, we observed highly increased levels of Keratin 6a, 6b and 16 in the skin of K5Mα mice on day three after induction of meprin α overexpression. McGowan and colleagues as well as several other studies (reviewed in reference^31^) showed that the expression of *Krt6*, *Krt16* in keratinocytes is rapidly induced within a few hours after wounding^70^, a very characteristic event, since these keratins are almost absent in keratinocytes of the interfollicular epidermis under homeostatic conditions. Therefore, keratin 6 and 16 serve as biomarkers for keratinocytes that acquire a highly proliferative phenotype in response to an impaired epidermal barrier function.

Both keratinocyte hyperproliferation and epidermal barrier defects either due to wounding or skin diseases are accompanied by inflammatory responses. Interestingly, Hasegawa and colleagues showed that dermokine also acts as a negative regulator of ELR^+^CXC chemokine expression^64^. In brief, *Cxcl1* expression in wounds of mice as well as CXCL1 and CXCL8 levels in human keratinocyte cultures stimulated with IL-1β were significantly reduced after treatment with recombinant dermokine^64^. Our data show that particularly CXCL1, CCL2, G-CSF and GM-CSF are highly elevated in the skin of K5Mα mice already on day two. Hence, we suggest that these mediators are the drivers of the subsequently aggravating inflammatory response characterized by an increasing infiltration of neutrophilic granulocytes specifically into areas of high meprin α levels, followed by an accumulation of inflammatory cytokines like IL-17A, IL-6, IL-36 and TNFα as well as acute phase proteins. With regard to the mechanistic insights provided by Hasegawa and colleagues, proteolytic inactivation of dermokine by meprin α might be causative for the excessive release of CXCL1 and, thereby, initiate the subsequent inflammatory cascade observed in K5Mα mice.

Supporting this hypothesis, there is a striking overlap between dysregulated gene expression patterns reported in *Dmkn* αβγ-deficient mice and proteomic changes observed in the skin of K5Mα mice. For example, *Krt6b*, *S100a8* and *S100a9* are among the highest upregulated genes in *Dmkn* αβγ-deficient mice as well as K5Mα mice^59^. Moreover, we provide longitudinal multi-omics data sets that comprise an extensive content of information on various aspects of the K5Mα phenotype development that we have not discussed here in detail, e.g. changes in the cell-cell/matrix adhesion proteins, the cytoskeleton, the protease web, the inflammatory cascade and many more.

Interestingly, Utsunomiya and colleagues reported that 15% of *Dmkn* αβγ-deficient mice developed a very severe skin phenotype and died within 21 days after birth^59^. In K5Mα mice systemically treated with tamoxifen, we even observed 100% mortality within 18 days after switching to tamoxifen-supplemented food (data not shown). Such an escalating phenotype is not observed in *Dmkn* βγ-deficient mice because loss of dermokine β and γ might be compensated by increased expression of dermokine α^57,59^. This would be a reasonable explanation for the transient skin phenotype observed in *Dmkn* βγ-deficient mice, which in contrast persists and even progresses in *Dmkn* αβγ-deficient mice. In K5Mα mice we detected by mass spectrometry markedly reduced levels of dermokine on day one, which increased to similar levels detected in control mice on day three. These changes within the first three days could indicate an initial compensatory phase in response to dermokine cleavage by meprin α. Adaptation to dermokine loss in *Dmkn* αβγ-deficient mice could also be a reasonable explanation for the differences in mortality compared to K5Mα mice. While viable born *Dmkn* αβγ-deficient mice compensated the loss of dermokine already from early embryonic development on, K5Mα mice experience an acute and rapid degradation of dermokine at an age of 8 to 15 weeks through tamoxifen-induced meprin α expression. Overall, there is increasing evidence for major differences between the molecular and cellular responses initiated by a gene not being expressed, e.g. due to knock-out, in comparison to the encoded protein being post-translational functionally inactivated, e.g. by proteolysis^71^.

In this context, our TAILS data indicate increased cleavage of dermokine isoforms β and γ in the skin of K5Mα mice. Notably, both isoforms contain the meprin α cleavage site identified by TAILS. Having a closer look at the proteolytic processing of dermokine, most of the highly abundant peptides identified by TAILS are located near the C-terminal globular domain 2, which has been described by Higashi and Hasegawa as the functional domain of dermokine with regard to modulation of ERK signaling and wound healing^60,64^.

In contrast, the prominent CTF generated by meprin α results from cleavage between G92 and E93. Therefore, we assume that the initial cleavage by meprin α makes dermokine prone to additional subsequent proteolytic events executed by meprin α or other proteases. The amino acid sequence in position P3 to P3’ to the identified cleavage site is highly favored by meprin α, indicated by peptide score of 429. Additionally, meprin α cleavage of dermokine particularly at this site would explain why we did not detect the corresponding peptide by TAILS. Cleavage between R96 and S97, which we identified by TAILS and that is probably caused by trypsin digestion during sample preparation, causes a loss of the peptide comprising this neo N-terminus ([G].E_93_AAR_96_S_97_LGNAGNEIGR). We assume that we were only able to detect this neo N-terminus due to the enrichment of dermokine and its CTFs by immunoprecipitation. AlphaFold 3 models of dermokine and sequence alignment analyses suggested that the meprin α cleavage site is located within a highly conserved, potentially alpha helical folded region of dermokine. While AlphaFold 3-based structural predictions for most of the dermokine sequence were very uncertain, particularly this alpha helix was reproducible modeled, which might indicate a relevance of this domain for the folding, stability and/or signaling of dermokine. Since the cleavage site we identified is located within the center of the second helix, we speculate that proteolytic processing by meprin α can compromise the function of dermokine.

In summary, we propose that meprin α functionally counteracts dermokine function by its degradation and, thereby, meprin α indirectly regulates keratinocytes proliferation and immune cell recruitment (**Figure 9**). Moreover, available data on their epidermal localization and expression levels indicate that meprin α-mediated cleavage of dermokine might represent a relevant physiological mechanism in wound healing and a pathophysiological mechanism in skin diseases like PV. Interestingly, we found that catalytically inactive meprin α E155A also interacts with dermokine. Hence, complex formation of dermokine and meprin α already on the secretory pathway is a potential alternative mechanism that could limit the amount of functionally active dermokine. Of note, this would we reasonable explanation why topical treatment with actinonin for inhibition of meprin α did not prevent or delay phenotype development in K5Mα mice. However, it has to be noted that actinonin is not a meprin α-specific inhibitor, but acts via its hydroxamate group as a chelating agent for zinc ions of also other metalloprotease in the skin^72^, which might explain skin irritations observed by TEWL measurements in control mice. Therefore, further mechanistic analyses including e.g. transgenic mice for inducible overexpression of a catalytically inactive meprin α mutant should be investigated in the future.

**Figure 9:**
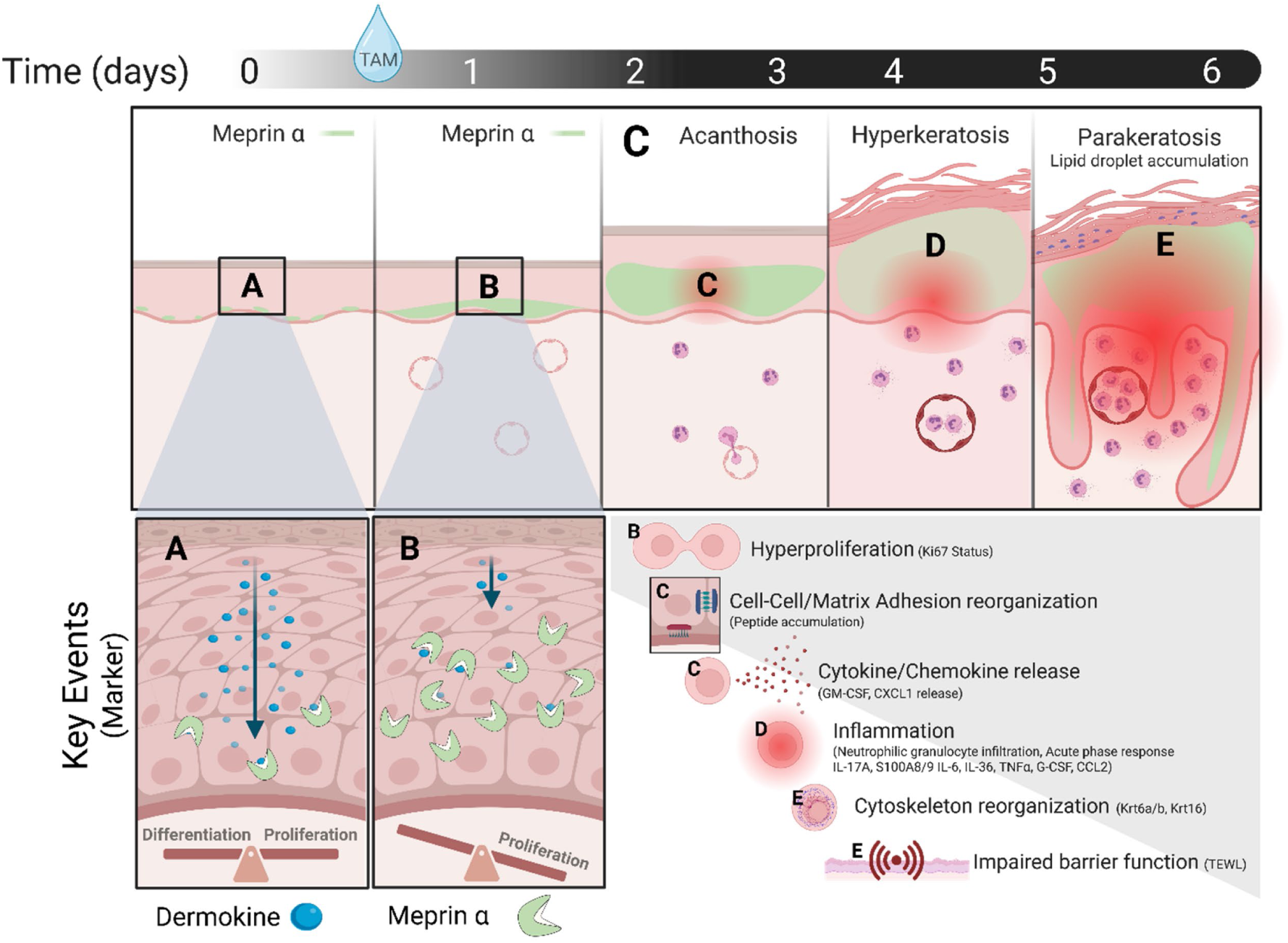
Summary of the skin phenotype development in K5Mα mice. **(A)** Under homeostatic conditions dermokine is secreted by differentiating suprabasal keratinocytes to suppress keratinocyte proliferation and release of proinflammatory cytokines/chemokines. Meprin α synthesized at low levels by basal keratinocytes partially degrades dermokine to balance proliferation and initiation of terminal differentiation. **(B)** Topical 4-hydroxytamoxifen (TAM) treatment in K5Mα mice initiates overexpression of meprin α. Increased dermokine degradation by meprin α shifts the balance towards keratinocyte proliferation and elevated secretion of proinflammatory cytokines/chemokines. **(C)** Keratinocyte hyperproliferation leads to epidermal hyperplasia (acanthosis) and is associated with reorganization proteins involved in cell-cell/matrix adhesion. Increased release of GM-CSF and CXCL1 recruits **(D)** neutrophilic granulocytes to hyperplastic epidermal areas, which foster a local inflammatory response characterized by high levels of IL-17A, S100A8, S100A9, IL-6, TNFα, G-CSF, CCL2 and acute phase proteins. Due to hyperproliferation an excessive number of keratinocytes enter terminal differentiation and form a thickened *stratum corneum* (hyperkeratosis). **(E)** Continuously increasing infiltration of neutrophilic granulocytes boosts the local inflammatory response. The excessive number of keratinocytes rapidly transitioning into the *stratum granulosum* leads to an incomplete differentiation of corneocytes characterized by partial degradation of the nucleus (parakeratosis) and impaired processing of lamellar bodies and keratohyalin granules in the *stratum granulosum*. Defective differentiation of the stratum corneum indicated by increased expression of barrier alarm keratins 6a, 6b and 16 is associated with an increased transepidermal water loss. Graphical illustration was created with BioRender.com.

In addition to the identification of dermokine as a novel epidermal meprin α substrate, we provide a novel disease mouse model. K5Mα mice develop a highly reproducible, genetically inducible psoriasis-like phenotype based on pathologically elevated meprin α expression as observed in PV. Therefore, the K5Mα mouse model might be a good alternative or complement, e.g. to commonly used model systems like the imiquimod model, for the community to study the molecular mechanisms underlying hyperproliferative and inflammatory skin diseases.

## Materials and Methods

### Chemicals

All chemicals were purchased of analytical grade from Merck, Carl Roth, Thermo Fisher Scientific or Roche.

### Animal Care

Animal acquisition, breeding, accommodation, care and usage in experimental procedures was performed in accordance with the guidelines of the German National Committee for the Protection of Animals Used for Scientific Purposes. Animals were housed under specific pathogen-free conditions in an IVC (individually ventilated cage) system at constant environmental conditions (12 h light-dark cycle, 22 ±2°C, 55 ±10% relative humidity). A maximum of five male or female siblings were housed in one cage with food and drinking water supply *ad libitum*. Cages were enriched with aspen litter, houses and tubes. Daily animal care was assured by professional animal care attendants.

### Generation of C57BL/6J-KRT5^tm1.1-CreERT2^;ROSA26^Stop-mMep1a^ mice

Meprin α knock-in mice (C57BL/6J-ROSA26^Stop-mMep1a^) were generated at the Transgenic Core Facility of the Max Planck Institute for Molecular Cell Biology and Genetics in Dresden, Germany. In advance, murine meprin α cDNA (Transcript ID: ENSMUST00000117137.8; A95-G2335) was cloned into pCAG-ROSA-IRES-EGFP targeting vector. Murine JM8A1.N3 embryonic stem cells^73^ were transfected by electroporation and clones selected that incorporated the transgenic construct (CAG-loxP-Neo^R^-STOP-loxP-mMep1a-HA-IRES-GFP) by homologous recombination into the Rosa26 locus of their genome. Suitable ES clones were transplanted by laser-assisted microinjection into C57BL6/NCrl morula stages 2.5 days post coitum^74^ for fertilization of recipient female mice. Sperms of chimeric male offspring were screened by short tandem repeat analysis for transgenic proportion and suitable samples were selected for *in vitro* fertilization. Heterozygous offspring was crossed with B6N.129S6(Cg)-Krt5^tm1.1(cre/ERT2)Blh^/J mice^75^ (Jackson Laboratory, stock #029155) to generate C57BL/6J-KRT5^tm1.1-CreERT2^;ROSA26^Stop-mMep1a^ mice (patent application EP20160475.8/W0 2021/175738 A1). Mice that carried homozygous the mMep1a construct within the Rosa26 locus and either heterozygous the information for the Cre recombinase/modified estrogen receptor fusion protein (CreER^T2^+/-; K5Mα) or not (CreER^T2^-/-; Control) were used for subsequent experiments. In order to breed only mice with a heterozygous and not homozygous CreER^T2^ status, CreER^T2^-positive mice were only crossed with CreER^T2^- negative mice.

### Genotyping

For genotyping of C57BL/6J-KRT5^tm1.1-CreERT2^;ROSA26^Stop-mMep1a^ mice genomic DNA was extracted from ear punches using the DirectPCR^®^ Lysis Reagent Tail (Peqlab, Germany). Ear punches were incubated overnight at 55°C and 300 rpm in 200 µl DirectPCR^®^-Tail buffer diluted with 200 µl nuclease-free water and supplemented with 9 IU/ml (0.3 mg/ml) recombinant proteinase K (Thermo Fisher, Germany). Afterwards, proteinase K was heat-inactivated by incubation at 85°C for 45 min. For subsequent PCR reaction 2 µl lysate was added to a total reaction volume of 30 µl containing DreamTaq DNA polymerase (80 IU/ml), 3 µl DreamTaq Green Buffer (10X), 2 mM dNTPs and respective primers at a concentration of 0.4 µM. Analysis of the Rosa26 locus zygosity for wildtype (wt) sequence and transgenic (tg) sequence was performed using the following primer set: forward (wt/tg) 5’-aaagtcgctctgagttgttatc-3’; reverse (wt) 5’- gatatgaagtactgggctctt-3’; reverse (tg) 5’- tgtcgcaaattaactgtgaatc-3’. PCR products at a size of 570 bp and 380 bp indicated wt and tg Rosa26 locus, respectively. Analysis of the CreER^T2^ status was performed using the following primer set: forward 5’- ggttcgcaagaacctgatggacat-3’; reverse 5’- gctagagcctgttttgcacgttca-3’. A PCR product at a size of 342 bp indicated insertion of the CreER^T2^ construct downstream of the keratin 5 promotor. Following PCR programs were run for genotyping of Rosa26 and CreER^T2^ status: i) Rosa26: 95°C for 3 min, 35 cycles (95°C for 30 s, 56°C for 30 s, 72°C for 50 s), 72°C for 10 min; ii) CreER^T2^: 95°C for 5 min, 30 cycles (95°C for 30 s, 56°C for 30 s, 72°C for 15 s), 72°C for 10 min.

### Tamoxifen treatment

Mice were either fed tamoxifen-supplemented food or topically treated with 4-hydroxytamoxifen (TAM) for local induction of meprin α overexpression. Food pellets supplemented with 400 mg/kg tamoxifen citrate (product ID: TD.130860) were purchased from Inotiv and fed weekly *ad libitum* from Monday to Friday. Saturday and Sunday mice were fed *ad libitum* with standard mouse keeping food pellets (Ssniff Spezialdiäten GmbH, Germany). For topical treatment (Z)-4-hydroxy tamoxifen (product ID: H7904; Merck, Germany) was solved in DMSO at a concentration of 5 mg/ml. In preparation, 24 h before the topical treatment a 2 cm² area at the flank of the mice was shaved. A total volume of 40 µl TAM/cm² (125 µg TAM/cm²) was directly applied to the skin and air-dried.

### Isolation and cultivation of primary murine keratinocytes

For the isolation of primary murine keratinocytes CreER^T2^+/- K5Mα mice were sacrificed by cervical dislocation. Tails were dissected, disinfected by incubation for 5 min in povidone iodine (Betaisodona, Mundipharma) and washed thrice in 70% ethanol. Under a clean bench tail skin was removed from the bone and cut into pieces of 0.25 to 0.5 cm². Skin pieces were placed with the dermal site facing a sterile filter paper soaked with DMEM supplemented with 1% bovine trypsin and incubated at 37°C and 85% relative humidity for 1 h. During incubation, culture dishes were coated with 5 µg/cm² rat tail collagen I (product ID: sc-136157; Santa Cruz Biotechnology). Collagen I was diluted in 20 mM acetic acid and dishes were incubated for 1 h at room temperature. Afterwards, dishes were washed thrice with sterile PBS. Epidermis of the tail skin was separated from the dermis using forceps, rinsed with 10% FCS/PBS and grinded through a 40 µm cell strainer. Cell suspension was centrifuged at 200 g for 5 min, cells were resuspended in CnT-57 medium (CELLnTEC, Switzerland) supplemented with 10 µg/ml gentamycin, 100 U/ml penicillin and 100 µg/ml streptomycin and seeded into collagen-coated culture dishes. Keratinocytes were cultured at 37°C, 5% CO_2_ and 85% relative humidity. Culture medium was replaced by fresh CnT-57 medium twice per week until keratinocyte cultures reached 20-30% confluency. Then, keratinocytes were immortalized by stable transfection with a plasmid encoding the large T antigen of the simian virus 40. TurboFect^TM^ (Thermo Fisher Scientific) was used as transfection reagent according to the manufacturer’s instructions and transfection medium was replaced by fresh CnT-57 medium 6 h after transfection. Within the following six weeks immortalized keratinocyte clones formed colonies while non-immortalized cells underwent cell death. Culture medium was replaced twice per week by fresh CnT-57 medium and keratinocyte cultures were passaged at 90% confluency. For this purpose, culture medium was removed, keratinocytes were washed with PBS and treated with 0.5% trypsin-EDTA (Gibco, Thermo Fisher Scientific) diluted in PBS for 10-15 min at 37°C. Reaction was stopped by addition of an equal amount of 10% FCS/PBS, keratinocytes were centrifuged at 200 g for 5 min, resuspended in pre-warmed fresh CnT-57 medium and seeded into culture dishes. For *in vitro* induction of meprin α overexpression, 1 µM TAM/DMSO was added to the culture medium of a culture at 50% confluency and keratinocytes were cultured for four days before the medium was replaced. Before TAM-treated keratinocytes were used for subsequent experiments, they were cultured at least for seven days in CnT-57 medium without TAM.

### Calcium differentiation

For the *in vitro* induction of terminal differentiation, murine keratinocytes were cultured to a confluency of 80%. Then, culture medium was replaced by CnT-57 medium supplemented with 540 µM CaCl_2_ and keratinocytes were cultured for up to 7 days at 37°C, 5% CO_2_ and 85% relative humidity. Differentiation process was monitored by light microscopy and detection of changes in differentiation associated proteins by western blotting.

### Preparation of conditioned medium

For the preparation of conditioned medium, supernatants from murine keratinocytes cultures that reached >95% confluency within three days were used. Supernatants were transferred to sterile tubes and centrifuged for 30 min at 10,000 g. Then, supernatants were transferred to fresh sterile tubes, supplemented with 5 µg/ml sterile-filtered bovine trypsin/PBS and incubated for 30 min at 37°C. Afterwards, 10 µg/ml sterile-filtered ovomucoid/PBS were added to the supernatant and incubated for 30 min at 37°C to inhibit trypsin activity. These conditioned supernatants were directly used for subsequent culture of murine keratinocytes.

### Skin lysate preparation

After mice were sacrificed, respective skin areas were shaved, excised, minced, snap frozen in liquid nitrogen and 100-150 mg grinded in a liquid nitrogen-cooled mortar. Grinded skin samples were transferred to a 1.5 ml tube with a screw cap. Next, 400 µl lysis buffer (50 mM Tris-HCl, 150 mM NaCl, 1% (v/v) Triton X-100, 1% (w/v) SDS, 2 mM EDTA, Complete^®^ Protease Inhibitor Cocktail, PhosSTOP Phosphatase Inhibitor Cocktail, pH 6.8) were added to the skin. Notably, skin lysates intended for meprin α activity measurements were generated in lysis buffer without EDTA. Afterwards, lysates were treated in three cycles with an ultrasonic homogenizer (Micro Ultrasonic Cell Disruptor KT 50 with CV18 Converter and 1/8” probe, Kontes) at 20% power 5x2 s/cycle with a cooling phase of 5 min between each cycle. Then, ceramic beads were added and lysates were subjected to a Precellys 24 tissue homogenizer equipped with Cryolys adapter (bertin technologies). Liquid nitrogen-cooled lysis was performed in three cycles each at 6,500 rpm for 30 s with a break of 30 s between cycles. Finally, lysates were incubated for 1 h on ice, centrifuged at 20,000 g and 4°C for 30 min and supernatants were transferred to fresh tubes. Protein concentrations were determined by BCA assay according to the manufacturer’s instructions (Pierce^TM^ BCA Assay Kit, Thermo Fisher Scientific) and lysates were stored at −20°C until subsequent analysis.

### Transient transfection

In preparation, 3 x 10^6^ HEK293T cells were seeded in 10 ml DMEM + GlutaMAX (Gibco, Thermo Fisher Scientific) supplemented with 10% (v/v) heat-inactivated FCS (Thermo Fisher Scientific), 100 U/ml penicillin, 100 µg/ml streptomycin (Gibco, Thermo Fisher Scientific) into a 10 cm (58 cm²) culture dish. HEK293T cells were cultured for 24 h at 37°C, 5% CO_2_ and 85% relative humidity. For transient transfection 3 µl polyethylenimine (Merck) per microgram DNA were mixed in DMEM with respective plasmids. After incubation for 20 min at room temperature, transfection mix was added dropwise to the culture supernatant under constant agitation of the culture dish. Twenty-four hours after transfection, culture medium was removed, cells were washed with PBS and 5 ml DMEM + GlutaMAX supplemented with 100 U/ml penicillin and 100 µg/ml streptomycin were added to the cells. Additional 24 h later, supernatants were collected and cells were subjected to lysis.

The following plasmids were used for overexpression of target proteins by transient transfection of HEK293T cells:

| Vector | Transgen/Tag (species) | Transcript ID (ORF 5'-3') | Mutation |
| --- | --- | --- | --- |
| pCMV6 | *Mep1a* (*Mus musculus*) | ENSMUST00000117137.8<br>(A95-A2338) | wildtype |
| pCMV6 | *Mep1a* E155A<br>(*Mus musculus*) | ENSMUST00000117137.8<br>(A95-A2338) | A558C |
| pCMV6 | *Dmkn*/Myc-DDK [C-term]<br>(*Mus musculus*) | ENSMUST00000085691.11<br>(A162-G1685) | wildtype |

### Cell lysis

Culture medium of HEK293T cells or murine keratinocytes was removed and cells were washed twice with PBS. Cells were taken up in lysis buffer (50 mM Tris-HCl, 150 mM NaCl, 1% (v/v) Triton X-100, 1% (w/v) SDS, 2 mM EDTA, Complete^®^ Protease Inhibitor Cocktail, PhosSTOP Phosphatase Inhibitor Cocktail, pH 6.8) and treated in three cycles with an ultrasonic homogenizer (Micro Ultrasonic Cell Disruptor KT 50 with CV18 Converter and 1/8” probe, Kontes) at 20% power 3x2 s/cycle with a cooling phase of 5 min between each cycle. Cell lysates intended for meprin α activity measurements were generated in lysis buffer without EDTA. Finally, lysates were incubated for 1 h on ice, centrifuged at 20,000 g and 4°C for 30 min and supernatants were transferred to fresh tubes. Protein concentrations were determined by BCA assay according to the manufacturer’s instructions (Pierce^TM^ BCA Assay Kit, Thermo Fisher Scientific) and lysates were stored at −20°C until subsequent analysis.

### Trichloroacetic acid precipitation

Isolated culture supernatants were centrifuged at 186,000 g (Optima^TM^ MAX-XP ultracentrifuge, Beckman Coulter), 4 microns and 4°C for 2 h to remove cell debris and extracellular vesicles. Afterwards, 900 µl supernatant were mixed with 100 µl trichloroacetic acid (TCA; >99%) and incubated one ice for 30 min. Then, supernatants were centrifuged at 20,800 g and 4°C for 20 min. The supernatant was discarded and the sediment was washed with −20°C cold acetone. Again, the sediment was centrifuged at 20,800 g and 4°C for 15 min, the acetone was decanted, followed by two additional washing steps with −20°C cold acetone to remove the excess TCA from the precipitated proteins. Then, acetone was removed from the sediment and sediment was dried overnight to remove excess acetone. Finally, the sediment was resuspended in 50 µl SDS-PAGE sample buffer (50 mM Tris-HCl, 70 mM SDS, 1.5 mM bromophenol blue, 10% (v/v) glycerol, 20 mM DTT in ddH_2_O, pH 6.8) and incubated for 10 min at 95°C and 650 rpm until the sediment was completely dissolved. In case the color of the sample buffer turned from blue to yellow, indicating acidification due to excess TCA in sediment, pH value was adjusted by titration with 1 M sodium hydroxide before incubation.

### (Co-)immunoprecipitation

(Co-)immunoprecipitation was performed with the Dynabeads^TM^ Protein G Immunoprecipitation Kit (Invitrogen, Thermo Fisher Scientific) according to the manufacturer’s instructions. Transfected HEK293T cells were washed three-times with ice-cold PBS and lysed with EBC lysis buffer (120 mM NaCl, 50 mM Tris-HCl, 0.5% (v/v) NP-40, Complete^®^ Protease Inhibitor Cocktail with EDTA in ddH_2_O, pH 7.4). Lysates were incubated twice for 10 s in an ultrasonic bath, incubated on ice for 30 min and centrifuged at 13,500 g and 4°C for 30 min. Supernatants were transferred to fresh tubes and protein concentrations were determined by BCA assay according to the manufacturer’s instructions (Pierce^TM^ BCA Assay Kit, Thermo Fisher Scientific). For lysate controls, a lysate volume containing 250 µg protein was separated. Afterwards, a lysate volume F3165, Merck) or 10 µl of a meprin α anti-serum isolated from an immunized rabbit (custom production by Pineda Antibody Service, Germany) overnight on a roller mixer at 4°C. Dynabeads^TM^ magnetic beads were vortexed for 30 s and for each immunoprecipitation sample 50 µl bead solution were transferred to a fresh tube. Beads were separated from the supernatant with a magnet, supernatant was discarded, beads were washed with the provided Binding & Washing Buffer and incubated overnight on a roller mixer at 4°C. The following day, Binding & Washing Buffer was removed from the beads, beads were mixed with the lysates that have been incubated with the antibodies and samples were incubated for 15 min on a roller mixer at room temperature. Next, samples were placed in the magnet, supernatant was transferred to a fresh tube and remaining beads were washed thrice with the provided Washing Buffer. Finally, beads were resuspended in 100 µl Washing Buffer, transferred to a fresh tube and Washing Buffer was removed. Afterwards, beads were resuspended in 70 µl SDS-PAGE sample buffer (50 mM Tris-HCl, 70 mM SDS, 1.5 mM bromophenol blue, 10% (v/v) glycerol, 20 mM DTT in ddH_2_O, pH 6.8) and incubated for 10 min at 95°C. Supernatant was separated from the beads with a magnet and subjected to SDS-PAGE.

### Sodium dodecyl sulfate polyacrylamide gel electrophoresis

SDS-PAGE was performed with the MINI-Protean Tetra Cell system from Bio-Rad. In preparation, lysates were adjusted to an equal protein concentration with respective lysis buffer and diluted 5:1 with 5X SDS-PAGE sample buffer (250 mM Tris-HCl, 350 mM SDS, 7.5 mM bromophenol blue, 50% (v/v) glycerol, 100 mM DTT in ddH_2_O, pH 6.8). Samples were denatured by incubation for 10 min at 95°C. Discontinuous SDS-PAGE was carried out with a polyacrylamide content of 7.5%, 10%, 12%, 14% or 18% in the separating gel dependent on the desired resolution. Electrophoresis was run at constant voltage of 90 V in the collecting gel and 120 V in the separating gel. PageRuler^TM^ Prestained Protein Ladder (Thermo Fisher Scientific) was used as a molecular weight marker.

### Western blotting

Western blotting was performed with the Criterion^TM^ tank blotting system from Bio-Rad. For the transfer onto a polyvinylidene fluoride membrane in tank blot buffer (25 mM Tris-HCl, 200 mM glycine, 20% (v/v) methanol in ddH_2_O, pH 8.3) a constant current of 2.5 mA/cm² was applied for 2 h at 4°C. Afterwards, membranes were washed for 5 min in TBS-T (25 mM Tris-HCl, 150 mM powder in TBS-T for 1 h at room temperature. Then, membranes were incubated overnight on a roller mixer at 4°C with respective primary antibodies diluted in either 5% (w/v) skim milk powder/TBS-T or 5% (w/v) bovine serum albumin/TBS-T. The following day, membranes were washed twice in TBS-T, once in TBS and incubated for 1 h on a roller mixer at room temperature with horse radish-conjugated secondary antibodies in 5% (w/v) skim milk powder/TBS-T. After further washing steps, membranes were incubated for 2 min in WesternBright ECL HRP substrate (Advansta) and chemiluminescence was detected with the Amersham^TM^ ImageQuant^TM^ 800 biomolecular imager (Cytiva). Image analysis was performed with the ImageQuant TL software version 8.2 (GE Healthcare).

### RNA isolation

RNA was isolated from skin samples with the RNeasy^®^ Fibrous Tissue Mini Kit (Qiagen) according to the manufacturer’s instructions. Briefly, mice were sacrificed, and respective skin area was shaved. Then, skin area was excised and 50 mg skin sample was minced in buffer RLT supplemented with 1% (v/v) β-mercaptoethanol. Minced skin was snap frozen in liquid nitrogen and grinded in a liquid nitrogen-cooled mortar. Grinded skin sample was transferred to a fresh tube and solved in 300 µl buffer RLT supplemented with 1% (v/v) β-mercaptoethanol. Then, 590 µl nuclease-free water and 10 µl proteinase K (ready-to-use provided by the kit) were added and incubated for 10 min at 55°C. Afterwards, sample was centrifuged at 10,000 g for 3 min, supernatant was transferred to a fresh tube, mixed with 0.5 ml 96% ethanol and loaded onto a RNeasy Mini column. Column was centrifuged for 15 s at 8,000 g and flow-through was discarded. Then, 350 µl buffer RW1 were applied onto the column, centrifuged for 15 s at 8,000 g and flow-through was discarded. Subsequently, 10 µl DNase I (18.2 units) were diluted in 70 µl buffer RDD, added to the RNeasy membrane and incubated for 15 min at room temperature. Following this, RNeasy membrane was washed once with buffer RW1, twice with buffer RPE and centrifuged empty at 20,000 g for 1 min. Finally, elution was performed by adding 50 µl pre-warmed RNase-free water onto the RNeasy membrane, followed by incubation for 5 min and centrifugation at 8,000 g for 1 min. RNA concentration was measured with a NanoDrop ND-1000 spectrophotometer. Isolated RNA was stored at −80°C.

### Reverse transcription

Reverse transcription was performed with the RevertAid First Strand cDNA Synthesis Kit (Thermo Fisher Scientific) according to the manufacturer’s instructions. Briefly, 500 ng isolated RNA were mixed with 1 µl oligo dT primer and nuclease-free water to a volume of 12.5 µl and incubated for 5 min at 65°C. Then, the reaction mixture was cooled on ice for 5 min, followed by addition of 7.5 µl master mix comprising 0.5 µl RiboLock Rnase inhibitor (20 U/µl), 2 µl dNTP mix (10 mM), 4 µl Reaction Buffer (5X) and 1 µl RevertAid M-MuLV reverse transcriptase (200 U/µl). Reaction mix was incubated for 1 h at 42°C. Finally, reaction was stopped by incubation for 5 min at 70°C. Synthesized cDNA was stored at −20°C.

### Real-time polymerase chain reaction

Real-time polymerase chain reaction analyses were carried out with a qTower^3^ 384G Real-time Thermocycler (Analytik Jena). Each reaction was conducted in technical triplicates. The reaction volume of 10 µl comprised 5 µl Luna^®^ Universal qPCR Master Mix (New England Biolabs), 3 µl cDNA, 0.25 µM forward/reverse primer and RT-PCR grade water (Thermo Fisher Scientific). Following PCR program was run: 95°C for 1 min, 40 cycles (95°C for 15 s, 60°C for 30 s). After the last cycle, a melting curve analysis was run by increasing the temperature from 60°C to 95°C in increments of 0.5°C. Data analysis was performed with qPCRsoft384 software version 1.2 (Analytik Jena). Relative mRNA levels were calculated by 2^-ΔΔCt^-method with Gapdh mRNA levels as reference.

### Paraffin embedding and tissue sectioning

Excised skin samples were fixed by incubation in 4% (w/v) paraformaldehyde/PBS for 48 h at room temperature. Afterwards, skin samples were watered in tap water for 5 h. Tissue dehydration by ascending alcohol series started with 50% ethanol overnight and proceeded with 70% ethanol for 1 h, 95% ethanol for 30 min, 95% ethanol for 1 h, 100% ethanol for 1.5 h, 100% ethanol for 2 h, xylene for 1.5 h and ended by incubation in xylene for 1.5 h. Dehydrated skin samples were soaked with paraffin at 60°C overnight. Embedded skin samples were cut with a rotary microtome HistoCore BIOCUT (Leica Biosystems) into 5 µm sections and fixed on slides for subsequent staining.

### Hematoxylin and eosin staining

Skin sections were stained with Mayer’s hemalum (Merck) and 1% aqueous Eosin-G (Roth). Therefore, skin sections were deparaffinized in xylene and rehydrated by a descending alcohol series (2x 100%, 95%, 70%, 50% ethanol). Then, skin sections were rinsed in distilled water and incubated for 10 min in filtered Mayer’s hemalum solution. Afterwards, skin sections were rinsed twice with distilled water, rinsed once with distilled water acidified with HCl, rinsed twice with distilled water and incubated for 5 min in lukewarm tap water for hemalum development. Then, skin sections were rinsed twice with distilled water and incubated for 5 min in filtered aqueous Eosin-G solution acidified with acetic acid. After skin sections were rinsed once with distilled water, sections were dehydrated by an ascending alcohol series (70%, 90%, 96%, 2x 100% ethanol, 2x xylene), embedded in Entellan^TM^ mounting medium (Merck) and sealed with a coverslip. Imaging was performed with the Aperio CS2 brightfield scanner (Leica Biosystems) at 40x magnification. Aperio ImageScope software version 12.4.6.5003 (Leica Biosystems) was used for visualization and image analyses.

### Immunofluorescence staining

Skin sections were deparaffinized in xylene, rehydrated by a descending alcohol series (2x 100%, 95%, 70%, 50% ethanol) rinsed with distilled water and incubated for 10 min in PBS. Heat-induced antigen retrieval was conducted in pre-heated citric acid buffer (2 mM citric acid, 8 mM trisodium citrate trihydrate, 0.05% Tween-20, pH 6.8) for 20 min in a steamer. Afterwards, tissue sections were cooled to room temperature, three-times washed in PBS for 5 min each and blocked by incubation in 5% (w/v) IgG-free BSA/PBS for 1 h at room temperature in a wet chamber. Then, skin sections were incubated overnight at 4°C with respective primary antibodies diluted in 1% (w/v) IgG-free bovine serum albumin/PBS in a wet chamber. Afterwards, skin sections were washed thrice and incubated in for 1 h at room temperature in a wet chamber with fluorochrome-conjugated secondary antibodies in 1% (w/v) IgG-free BSA/PBS supplemented with 1.5 µg/ml 4’,6-diamidino-2-phenylindole (DAPI). Finally, skin sections were washed three-times, rinsed with distilled water, embedded in Fluorescence Mounting Medium (Dako) and sealed with a cover slip. Image acquisition was performed with the MICA microhub fluorescence microscope (Leica Biosystems). Noise reduction was conducted by instant computational clearing in THUNDER mode^76^. For visualization and image analysis Leica Application Suite X software (Leica Biosystems) and ImageJ (NIH) were used.

### Antibodies

| Target (-Conjugate)<br>[Clone] | Host<br>[Isotype] | Concentration/Dilution<br>(Application) | Manufacturer<br>(Catalog ID) |
| --- | --- | --- | --- |
| Meprin $\alpha$<br>anti-serum | rabbit | 1:5000 (WB)<br>10 μl/800 μg protein (IP) | Pineda Antibody Service |
| Cytokeratin 14<br>[LL002] | mouse<br>[IgG<sub>3</sub>] | 10 μg/ml (WB) | Abcam (ab7800) |
| Ki67 [D3B5] | rabbit<br>[IgG] | 1:200 (IF) | Cell Signaling Technology<br>(112202) |
| Meprin $\alpha$<br>[polyclonal] | goat | 1 μg/ml (IF) | Bio-Techne (AF4007) |
| Cytokeratin 16<br>[polyclonal] | rabbit | 1 μg/ml (IF) | Proteintech<br>(17265-1-AP) |
| S100A9 [D3U8M] | rabbit<br>[IgG] | 1:200 (IF) | Cell Signaling Technology<br>(73425) |
| CD45 [30-F11] | rat<br>[IgG<sub>2b</sub>] | 1 μg/ml (IF) | Santa Cruz Biotechnology<br>(sc-53665) |
| Rabbit IgG H+L<br>(HRP) [polyclonal] | goat | 1:5000 (WB) | Jackson ImmunoResearch<br>(111-035-003) |
| Mouse IgG H+L<br>(HRP) [polyclonal] | sheep | 1:5000 (WB) | Jackson ImmunoResearch<br>(515-035-003) |
| Goat IgG H+L<br>(AlexaFluor488)<br>[polyclonal] | donkey | 2 μg/ml (IF) | Thermo Fisher Scientific<br>(A-11055) |
| Rabbit IgG H+L<br>(AlexaFluor647)<br>[polyclonal] | donkey | 2 μg/ml (IF) | Thermo Fisher Scientific<br>(A-31573) |
| Phospho-Stat3 Y705<br>[D3A7] | rabbit<br>[IgG] | 1:2500 (WB) | Cell Signaling Technology<br>(9145) |
| Stat3<br>[124H6] | mouse<br>[IgG<sub>2a</sub>, $\kappa$] | 1:1000 (WB) | Cell Signaling Technology<br>(9139) |
| Phospho-NF-κB p65<br>S536 [93H1] | rabbit<br>[IgG] | 1:1000 (WB) | Cell Signaling Technology<br>(3033) |
| NF-κB p65<br>[C22B4] | rabbit<br>[IgG] | 1:1000 (WB) | Cell Signaling Technology<br>(4764) |
| Phospho-IκBα Ser32<br>[14D4] | rabbit<br>[IgG] | 1:1000 (WB) | Cell Signaling Technology<br>(2859) |
| IκBα<br>[44D4] | rabbit<br>[IgG] | 1:1000 (WB) | Cell Signaling Technology<br>(4812) |
| Stat5<br>[D2O6Y] | rabbit<br>[IgG] | 1:1000 (WB) | Cell Signaling Technology<br>(94205) |
| Phospho-Stat5<br>Tyr694 [C71E5] | rabbit<br>[IgG] | 1:1000 (WB) | Cell Signaling Technology<br>(9314) |
| Gapdh<br>[14C10] | rabbit<br>[IgG] | 1:5000 (WB) | Cell Signaling Technology<br>(2118) |
| FLAG-Tag [M2] | mouse<br>[IgG<sub>1</sub>] | 1:1000 (WB)<br>1 µg/800 µg protein (IP) | Merck (F3165) |
| Myc-Tag [9B11] | mouse<br>[IgG<sub>2a</sub>, κ] | 1:2500 (WB) | Cell Signaling Technology<br>(2276) |
| Cytokeratin 5<br>[polyclonal] | rabbit | 1:1000 (WB) | GeneTex (GTX113219) |
| E-Cadherin [24E10] | rabbit<br>[IgG] | 1:1000 (WB) | Cell Signaling Technology<br>(3195) |
| Filaggrin [polyclonal] | rabbit | 1:500 (WB) | Biolegend (Poly19058) |
| Caspase 14 [32] | mouse<br>[IgG<sub>1</sub>, κ] | 1:500 (WB) | Santa Cruz Biotechnology<br>(sc-136351) |
| Desmoglein 1<br>[EPR6766(B)] | rabbit<br>[IgG] | 1:1000 (WB) | Abcam (ab124798) |
| Gasdermin A<br>[EPR19858-104] | rabbit<br>[IgG] | 1:100 (WB) | Abcam (ab230768) |
| Fra-2<br>[polyclonal] | rabbit | 1 µg/ml (IF) | Santa Cruz Biotechnology<br>(sc-13017) |

### Primer

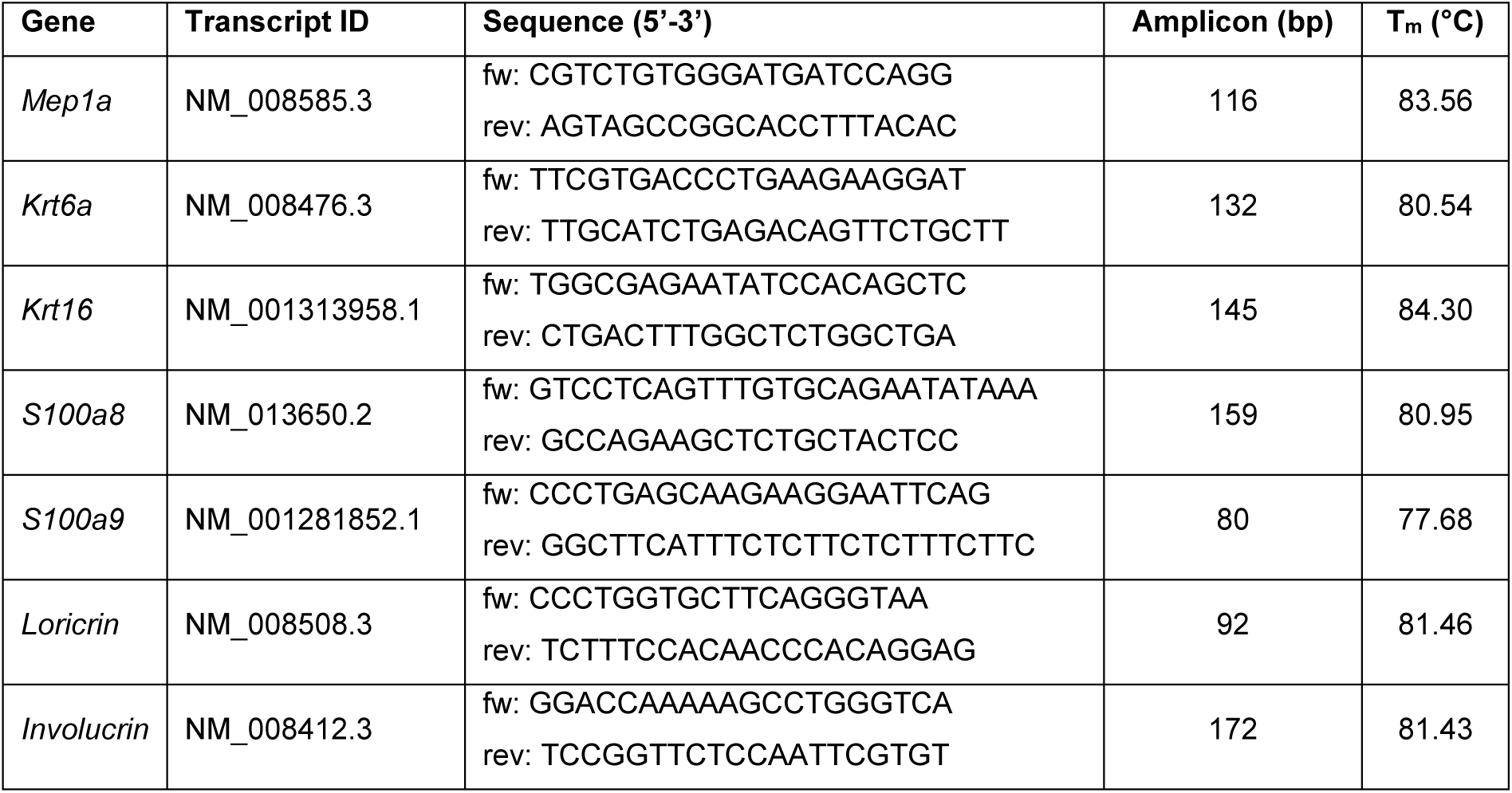
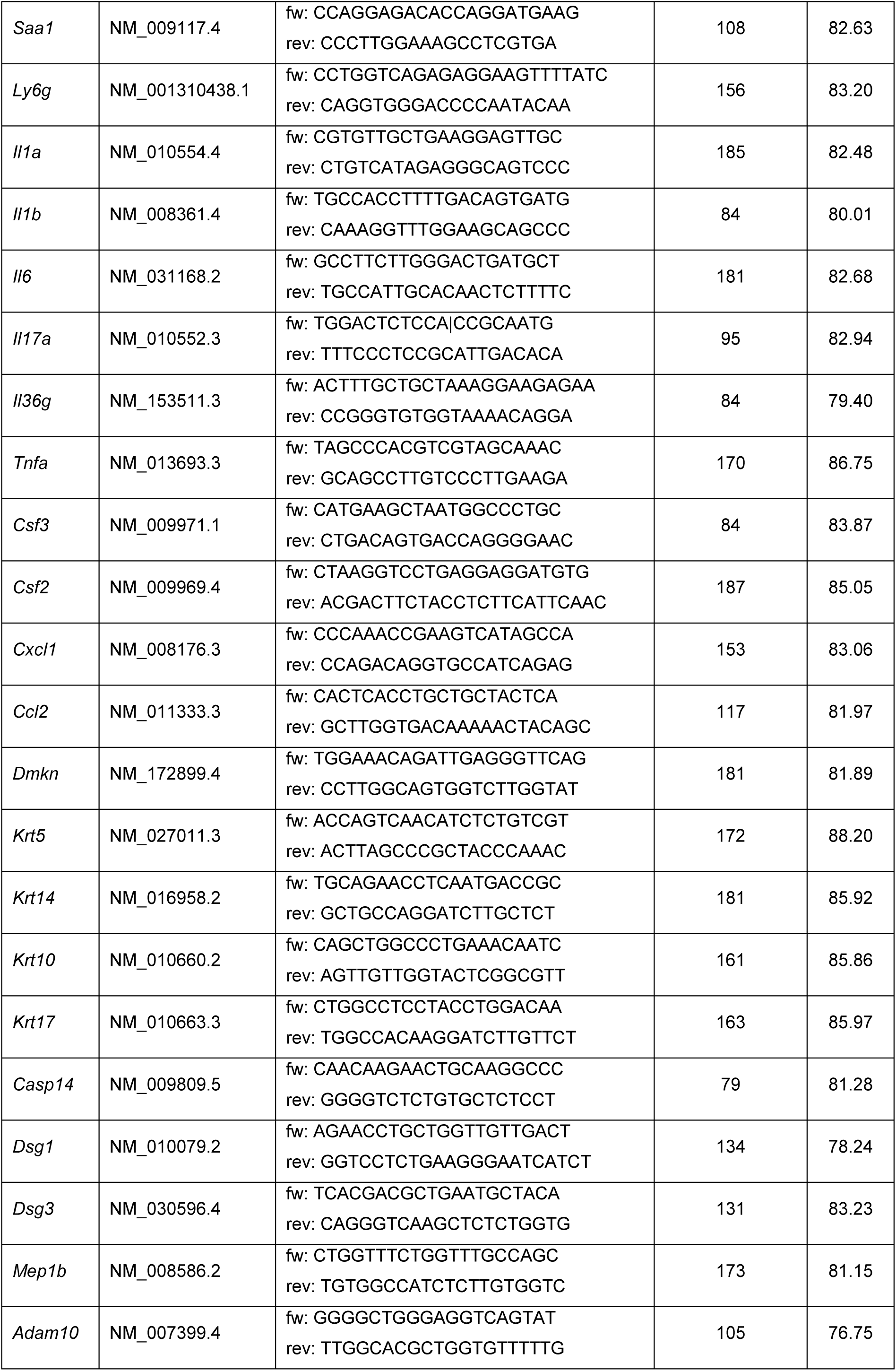
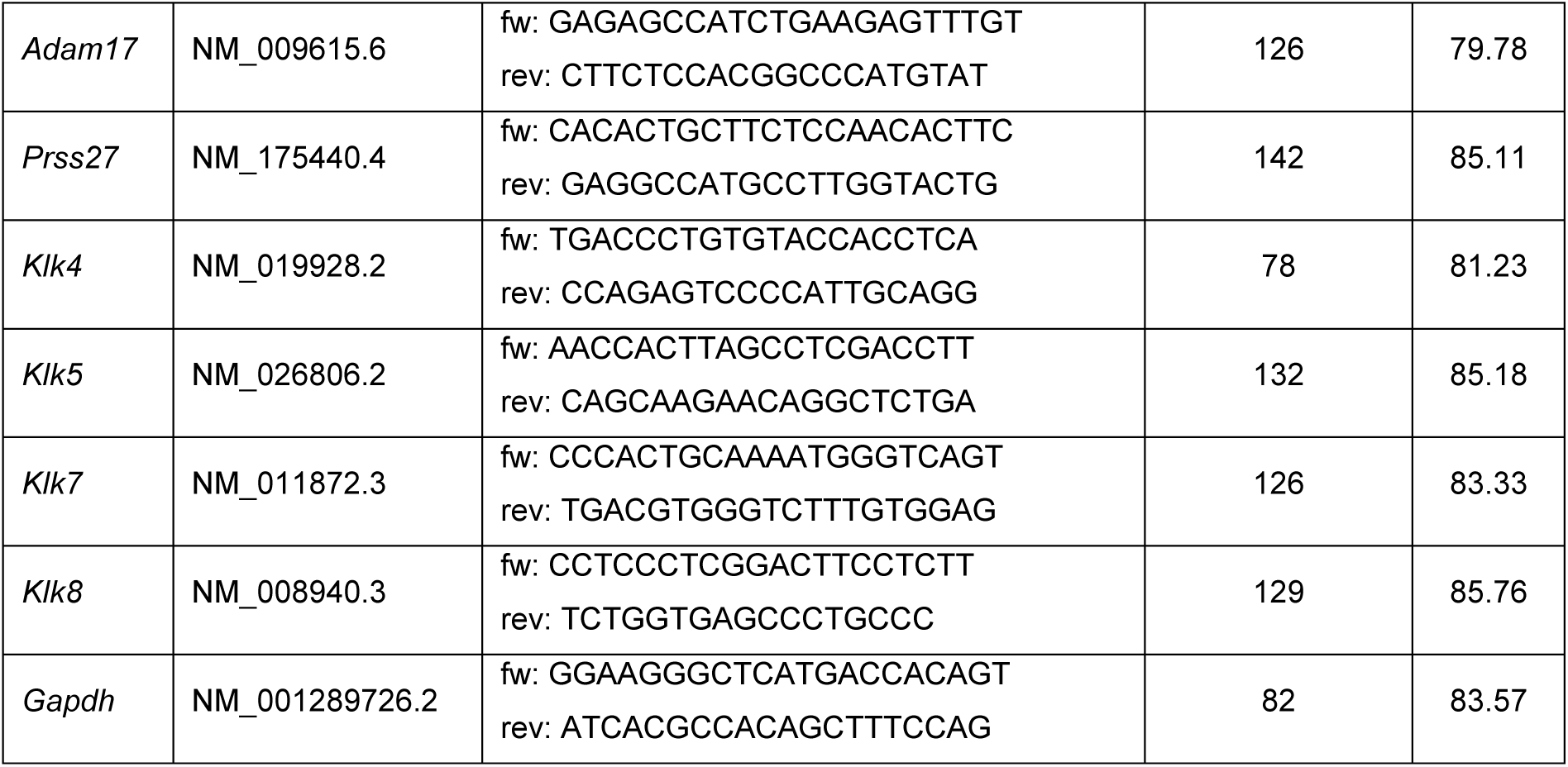

### Fluorogenic peptide-based activity assay

Meprin α activity in skin lysates and keratinocyte culture supernatants was measured by *in vitro* cleavage of a fluorogenic peptide substrate (mca-HVANDPIW-K-ε-dnp; Genosphere Biotechnologies). The peptide is specifically cleaved by meprin α^40^, N-terminal conjugated with a fluorophore (7-methyloxycoumarin-4-yl) and C-terminal conjugated with a quencher (K-ε-2,4- dinitrophenyl). Briefly, 100 µg skin lysate or 100 µl ultracentrifuged culture supernatant were mixed with 5 µM peptide substrate in a white 96-well flat bottom plate to reaction volume of 100 µl with PBS. For activation of meprin α in keratinocyte culture supernatants, 50 µg/ml bovine trypsin were added to the reaction volume. For inhibition of meprin α in keratinocyte culture supernatants, 10 µM actinonin was added to the reaction volume. Fluorescence intensity was measured for 2 h every 30 s at 37°C with a Tecan Spark spectrofluorophotometer (Tecan) with an excitation wavelength of 320 nm at an emission wavelength of 405 nm. All samples were measured in technical duplicates. Mean temporal changes in relative fluorescence intensities were calculated by linear regression in equal timeframes. Relative fluorescence intensity change per minute was defined as meprin α activity.

### Terminal amine isotopic labeling of substrates and mass spectrometry

In preparation for mass spectrometry analyses, proteins in skin lysates were precipitated by methanol and chloroform. In brief, methanol was added to skin lysates in a ratio of 4:1 and thoroughly mixed. Then, chloroform was added in a ratio of 1:5 and thoroughly mixed. Afterwards, distilled water was added in a ratio of 1:2, lysates were mixed and centrifuged for 2 min at 14,000 g. Top layer was removed, interface layer was separated and mixed with methanol at a ratio of 1:4. Finally, samples were centrifuged for 3 min at 14,000 g and methanol was discarded. Remaining sediment was dried and stored at −80°C until further processing.

Sample processing and N-terminal enrichment were performed according to the N-terminomics HUNTER protocol^77^, modified for TMT multiplexing^78^ with minor adaptations. Briefly, 50 µg protein per sample was resuspended in 100 µl TAILS buffer (250 mM HEPES pH 7.8, 2.5 M GuHCl) in protein low-bind tubes. Reduction and alkylation were performed by adding TCEP and freshly prepared CAA to final concentrations of 10 mM and 40 mM, respectively, followed by incubation at 95 °C for 10 min at 600 rpm. Each vial of 16-plex TMTpro (Thermo Fisher Scientific) label (200 µg) was dissolved in 55 µl anhydrous DMSO, mixed with an equal volume of sample, and incubated at room temperature for approximately 1 hour. The labeling reaction was quenched by adding 1 M ammonium bicarbonate (final concentration 100 mM) and incubating for 30 min at room temperature. Labeled samples were combined, aliquoted into 1.5 ml protein low-bind tubes (approx. 250 µl each), and magnetic beads (Sera-Mag^TM^ hydrophilic and hydrophobic carboxylate-modified SpeedBeads, Cytiva) were added at a 1:10 (protein:beads, w/w) ratio. Samples were incubated in 80% ethanol on a shaker (800 rpm) at room temperature for 15 min, followed by washing with 90% ethanol. Beads were then resuspended in 100 mM HEPES (pH 7.8) for final protein concentration of 1 µg/µl. Trypsin digestion (1:50 trypsin:protein, w/w) occurred overnight at 37 °C, 350 rpm. Following digestion, samples were mixed with equal volumes of 100 mM HEPES containing 2 M GuHCl, incubated at 85 °C for 5 min at 1000 rpm and centrifuged briefly. The supernatant was transferred, and 10% was set aside as a non-pullout (NPO) control stored at −20 °C. The remaining peptides underwent depletion of tryptic peptides by adding undecanal (50:1 undecanal:peptide, w/w) and NaBH3CN (50 mM final concentration), adjusting pH to 6-7, and incubating at 50 °C for 1.5 hours at 450 rpm. The reaction was acidified to pH 1.5-2.5 with Buffer H’ (40% ethanol, 5% TFA), adjusted to 600 µl with Buffer H (40% ethanol, 1% TFA), and loaded onto activated SepPak columns to isolate enriched N-terminal peptides (pullout, PO).

Both NPO and PO samples were desalted using activated SepPak columns (Waters), equilibrated with Buffer A’ (3% ACN, 1% TFA), washed with Buffer A (0.1% FA), and eluted with Buffer B (80% ACN, 0.1% FA). The eluates were dried in a speed vacuum concentrator and resuspended in Buffer A* (2% ACN, 1% TFA). Peptide concentrations were measured, and 500 ng peptides per sample were loaded onto EvoTips (EvoSep) for subsequent LC-MS/MS analysis.

### TAILS data acquisition

Samples were measured using the EvoSep One platform (Evosep, Denmark) coupled to an Orbitrap Exploris 480 mass spectrometer with a FAIMSpro interface (Thermo Fisher Scientific). Peptides were separated on an EV1106 C18 column (15 cm × 150 µm, 1.9 µm diameter, Evosep) using the Whisper100 nanoflow system with a 15 SPD (samples per day) method, comprising an 88-minute gradient. Eluting peptides were injected into the mass spectrometer through a 20 µm fused silica emitter (EV1087, Evosep) at a static voltage of 2300 V, with a carrier gas flow of 3.6 L/min and an ion transfer tube temperature of 240°C. FAIMS was operated at two CVs, −50 and − 70, with cycle time of 2 seconds each. MS1 scans were acquired at 120,000 resolution across a mass range of 375–1500 m/z, maximum injection time of 246 ms, normalized AGC target of 1.2e6, RF Lens at 40%. For data dependent scans, filters included charged ions (charge states 2–7), dynamic exclusion for 60 seconds with a ±10 ppm tolerance, minimum intensity threshold of 5e3, and precursor fit threshold of 0.7. Data-dependent MS2 scans were acquired in centroid mode at 60,000 resolution, with an isolation window of 0.7 m/z, first mass at 110 m/z, HCD fragmentation at 34% NCE, AGC target of 375,000, and a maximum injection time of 118 ms. Raw data were analyzed using Proteome Discoverer v2.4, searching against the mouse proteome database from Uniprot. Search parameters included semi-tryptic specificity, two maximum missed cleavages, dynamic modifications (methionine oxidation [+15.995], asparagine deamidation [+0.984], TMTpro labeling [+304.207], and protein N-terminal acetylation [+42.011]), and static modifications (cysteine carbamidomethylation [+57.021] and lysine TMTpro labeling). Peptide-spectrum matching utilized Sequest HT, with FDR controlled by Percolator at 1% strict and 5% relaxed thresholds. Peptides and proteins were quantified using the Reporter Ion Quantifier node and normalized to the total peptide amount per channel. Exported peptide group and protein reports underwent downstream analysis with CLIPPER 2.0^79^.

### Proteomics and TAILS data availability

The mass spectrometry proteomics data have been deposited to the ProteomeXchange Consortium via the PRIDE^80^ partner repository with the dataset identifier PXD063928.

### TAILS data filtering for identification of potential meprin α substrates

For the identification of peptides potentially generated by meprin α TAILS data were filtered according to the following scheme. First, all peptides without an N-terminal TMT-label and those that only did not contain the canonical N-terminal methionine were filtered out. Next, peptides derived from proteins annotated as plasma membrane or extracellular localized were filtered by Gene Ontology annotations. Subsequently, peptides that were higher abundant in the skin of K5Mα than control mice were filtered by cutoff-values. For this purpose, n-fold differences of the mean were calculated. In order not to overlook proteolytic events that might cause only minor changes in relative peptide abundance but have a relevant impact, a lower threshold on day one and two (≥ 1.2-fold) than on day three (≥ 1.5-fold) was defined. These peptides were filtered for the extracellular proteolytic accessibility of their N-terminus by the data deposited in the UniProt data base to ensure that these peptides could be potentially generated by meprin α. Afterwards, peptides derived from proteins that are potentially localized in the epidermis were identified by literature search. Here, only peptides derived from proteins that are not localized in the epidermis by high certainty, e.g. muscle specific proteins, were excluded from further analyses. Next, the peptides’ putative cleavage site was examined with regard to the cleavage specificity of meprin α. Therefore, peptides that contained an aspartate or glutamate residue in P1, P1’ or P2’ position or a proline residue in P2’ were filtered. Then, the amino acid sequence from P3 to P3’ position of the peptides was rated with regard to amino acids preferred by human meprin α. Here, the relative abundance of the respective amino acids annotated in the MEROPS database (https://www.ebi.ac.uk/merops/) was used. As readout, a Peptide Score was calculated which represents the sum of the likelihood-values for each amino acid in position P3 to P3’. A Peptide Score ≥ 300 was defined as a cutoff by calculating the median value of all analyzed peptides. In the same step, peptides that contained a highly disfavored amino acid (likelihood-value < 10) in positions P2 to P2’ and those that were derived by removal of a signal peptide were excluded. Finally, the raw signal intensities and changes in the signal intensities detected by TAILS were defined as selection criteria. Selection criteria included that signal intensities of candidate peptides have to increase in skin samples from K5Mα mice from day one to two or three and absolute intensity value must be higher than in skin samples from control mice on day one, two or three after TAM treatment.

### Mass spectrometry-based proteomics sample preparation and analysis for cleavage site identification

Dynabeads^TM^ magnetic beads from co-immunoprecipitation were resuspended in 4 M digestion buffer (4 M GuHCl and HEPES (pH 8.0)) and an on-bead digestion was carried out. Cysteine reduction was carried out using Tris-(2-carboxyethyl)-phosphine (TCEP) (5 mM final concentration) and alkylation with chloroacetamide (CAA) (20 mM final concentration) both carried out at 400 rpm, 65 ^◦^C for 30 minutes. After reduction and alkylation, samples were diluted 8-fold to reduce the GuHCl concentration to 0.5 M using 100 mM HEPES (pH 8.0) and LysC (Wako, Osaka, Japan) (1:100) was added for 4 hours followed by overnight digestion with sequencing-grade modified trypsin (1:50) (Sigma). The reaction was quenched using 1% TFA. The beads were centrifuged at 14,000 *x g* for 10 minutes and the supernatant was transferred to a fresh tube. Peptides were purified using Solaµ plates (Thermo Fisher Scientific, Waltham, MA), according to the manufacturer’s protocol.

Purified peptides were analyzed by LC-MS/MS using an Evosep One LC system (Evosep, Odense, DK) coupled to an Eclipse mass spectrometer (Thermo Fisher Scientific, Waltham, MA). Peptides were separated using the pre-installed 40 Sample-per-Day Whisper method. Chromatographic separation was performed using an Aurora Series Generation 3 15 cm × 75 µm inner diameter, 1.7 µm C18 analytical column (Ion Opticks, Victoria, AU). Mobile phase A consisted of 0.1% formic acid (FA) in water, and mobile phase B contained 0.1% FA in 80% acetonitrile (ACN).

Mass spectrometry data were acquired over 31 minutes in positive ion mode utilizing a data-independent acquisition (DIA) strategy. MS1 Orbitrap resolution was set to 120,000 within a scan range of 400-1000 m/z, with an AGC target of 300% or 1.2×10^6^ and the maximum fill time set to auto. For MS2, Orbitrap resolution was 60,000 with an AGC target of 1000% or 5×10^5^ and an auto maximum fill time, within a precursor scan range of 400-1000 m/z and a scan range of 200-1200 m/z. Automatic DIA window type was enabled, normalized collision energy was set to 33%, isolation windows were placed at 8 m/z with a window overlap of 1 m/z and optimal window placement enabled. Loop control was enabled and set to 25 number of spectra. FAIMS voltages were set to −50 compensation voltage for both experiments.

Raw data were analyzed via Spectronaut’s (version 19.7.250203.62635; Biognosys, Schlieren, CH) pulsar search engine, using the reviewed mouse fasta file (UP000000589). Mass spectra were matched using default settings and peptides were searched setting the Pulsar search to semi-specific, cleavage rules to Trypsin/P and LysC with a maximum and minimum peptide length of 48 and 6 amino acids. No imputation was performed and the MS1 area of proteotypic peptides was considered for quantification.

### Bead-based cytokine and chemokine multiplex

Cytokine and chemokine concentrations in culture supernatants and skin lysates were measured with a customized LEGENDplex^TM^ kit (Biolegend) according to the manufacturer’s instructions. The custom panels included beads, antibodies and standards for the detection and quantification of CXCL1, IL-34, IL-1α, IL-18, TNFα, IL-33, CCL2, IL-10, IL-6, G-CSF, IL-17A, GM-CSF (panel 1) and IL-1β or IL-1β IL-6, IL-10, IL-12p70, IL-17A, IL-23, IL-27, TNFα, CCL2, IFNβ, IFNγ and GM-CSF (panel 2). In preparation, all reagents and samples were pre-warmed at room temperature, supplied beads and antibodies were mixed with a vortex mixer for 5 min prior to use and standards were diluted according to the manufacturer’s instructions. Briefly, diluted standards and 25 µl ultracentrifuged culture supernatants or 25 µl skin lysate were mixed in a 96-V-bottom well plate with assay buffer and pre-mixed beads. After incubation for 2 h at 800 rpm and room temperature on a plate shaker, samples were centrifuged for 5 min at 250 g and washed with the provided Washing buffer. Premixed detection antibody mix was added to the samples and incubated for 1 h at 800 rpm and room temperature on a plate shaker. Then, PE-conjugated streptavidin was added to the samples and incubated for 30 min at 800 rpm and room temperature on a plate shaker. After two washing steps, samples were measured with a FACS Symphony A1 flow cytometer. Data analysis was conducted with the cloud-based LEGENDplex^TM^ Data Analysis Software Suite (https://legendplex.qognit.com/user/login?next=home) provided by Biolegend. Cytokine and chemokine levels quantified in skin lysates were normalized to the respective protein concentrations of the lysates determined by BCA assay.

### Transepidermal water loss measurement

Transepidermal waterloss was measured with the Multy Display Device MDD4 equipped with a Tewameter^®^ TM Nano probe (both Courage Khazaka). Each measurement was performed in technical triplicates for at least 30 s each. Mean transepidermal waterloss in g/h/m² was calculated from technical replicates for each time point for data analysis. For topical actinonin treatment, 100 µg/cm² actinonin (Merck) in DMSO (2 mg/ml) were applied twice daily onto the shaved skin of mice. An equal volume of DMSO was applied in the control group.

### Transmission electron microscopy

Skin samples were dissected and immersion fixed in a mixture of 4% paraformaldehyde and 1% glutaraldehyde in 0.1 M PBS at pH 7.4 overnight. Samples were rinsed three times in 0.1 M sodium cacodylate buffer (pH 7.2–7.4) and osmicated using 1% osmium tetroxide in cacodylate buffer. Following osmication, the tissue was dehydrated using ascending ethyl alcohol concentration steps, followed by two rinses in propylene oxide. Infiltration of the embedding medium was performed by immersing the pieces in a 1:1 mixture of propylene oxide and Epon and finally in neat Epon and hardened at 60°C. Vertical semithin sections (0.5 µm) from the skin were prepared for light microscopy, mounted on glass slides, and stained for 1 minute with 1% Toluidine blue. Consecutive ultrathin sections (60 nm) were examined in a EM902 (Zeiss, Germany). Pictures were taken with a MegaViewIII digital camera (A. Tröndle, Moorenweis, Germany).

### Sequence Logo

Sequence logos were generated with iceLogo web application (https://iomics.ugent.be/icelogoserver)^81^. Experimental sets comprised one-letter code of amino acids in P4 to P4’ position of peptides identified by mass spectrometry. A precompiled Swiss-Prot composition from *mus musculus castaneus* was used as reference set. Scoring system was set to “Percentage”, Start Position was set to “-4” and P value was set to “0.02”. “IceLogo” was selected as visualization type. Frequency difference of amino acids in P4 to P4’ position (x-axis) in the experimental data set compared to the reference data set was plotted on the y-axis in percentage. Amino acids in one-letter code were colored according to their chemical properties: polar (green/violet), basic (blue), acidic (red), hydrophobic (black).

### AlphaFold 3 Modelling

The structure of protein complexes was predict using AlphaFold 3^54^ via the official AlphaFold 3 server (https://alphafoldserver.com/about). The option “Seed” was set to auto and following amino acid sequences were used for modelling:

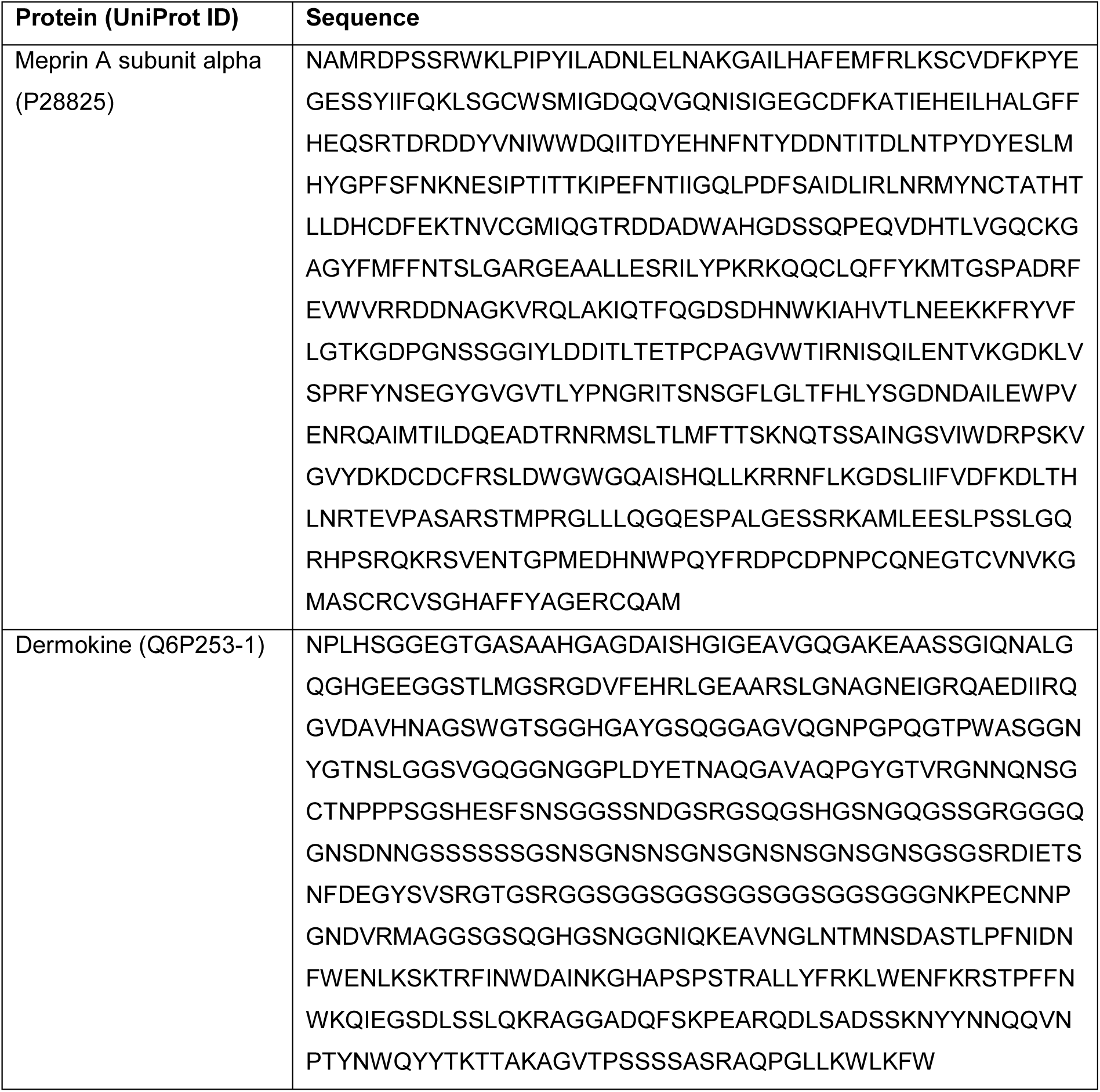

AlphaFold 3 Model visualization and analysis was performed with PyMOL^TM^ (Schrodinger, LCC).

### Statistics

Statistical analyses were performed using GraphPad Prism version 10.4.1 (GraphPad Software, LCC). Data sets were tested for Gaussian distribution and equal variance by Shapiro-Wilk and Equal Variance test, respectively. For comparison of two-groups comprising parametrically distributed data, two-tailed unpaired t-test was conducted. In case Shapiro-Wilk test was passed but datasets failed Equal Variance test, Welch’s correction was performed. Datasets that failed normality test were analyzed by two-tailed unpaired Mann-Whitney Rank Sum test. In all cases, confidences level was set to 95% assuming statistically significant differences for p-values < 0.05. Parametric data sets comprising multiple groups were compared by one-way analysis of variance (one-way ANOVA) for statistical significance. Correction for multiple comparisons was conducted by Tukey’s test. Non-parametrical datasets comprising multiple groups were analyzed with Kruskal-Wallis one-way ANOVA on ranks. Correction for multiple comparisons was conducted by Dunn’s test. Curve fitting was performed by simple linear regression or non-linear regression tool. Outliers were not eliminated for regression analysis and least squares regression was chosen as fitting method. For non-linear regression, convergence criteria were set to “medium” with automatic switch to “strict” when recommended. Maximum number of iterations was set to 1000. No weighting was performed. Regression start and end were set to “automatic” and 95% confidence bands were plotted for both linear and non-linear regressions. R and R square (R²) values for correlation between parametric data sets were computed by Pearson correlation. R and R² values for correlation between non-parametric data sets were computed by Spearman correlation. Two-tailed p-values were calculated for a confidence levels of 95% and statistically significant differences between groups were assumed for p < 0.05 indicated by an asterisk (*). Differences between groups that did not reach significance level were not marked.

### Ethical statement and study approval

All animal experiments were carried out in accordance with the European guidelines for care and use of laboratory animals and approved by the Ministry for Agriculture, Rural Areas, Europe and Consumer Protection of the state Schleswig-Holstein, Germany (reference V242 - 12659/2018 (30-4/18)).

## Acknowledgements

We thank Lennard Arp (former technical assistant at the Institute of Biochemistry, University of Kiel) and Alex Bitter (technical assistant at the Institute of Biochemistry, University of Kiel) for excellent technical assistance as well as Hila Emmert (former junior professor at the Clinic for Dermatology, Venerology and Allergology, UK-SH Kiel) for productive scientific exchange.

## Author contributions

VK, FP, CBP and SR designed the experiments. VK, FP, IG, EF, NB, FA, SB, CB, VC, MM, KB, MR, KK, MS and SR performed the experiments. VK, SR and CBP interpreted the data. MH, RN, FP, VK, IG, SB and SR generated the transgenic mice and bred the mice. IG, FA, VC, KK, UadK and SR performed sample preparation for the mass spectrometry analyses, measurements and data analyses. MS performed the electron microscopy analyses. IH and SR performed the TEWL analyses. SR wrote the manuscript. All authors proof-read the manuscript.

## Funding

This study was supported by the Deutsche Forschungsgemeinschaft (DFG) via the Collaborative Research Centre (CRC) 877 (Proteolysis as a Regulatory Event in Pathophysiology, Project A9) and Project 5098655290 (Knock-in mouse models for the characterization of meprin metalloproteases in hyperkeratosis, inflammation and systemic sclerosis) (both to CBP) as well as the Christian-Albrechts-University Kiel by intramural funding K126444 (Meprin α – A novel molecular driver and potential therapeutic target in sepsis) and the Cluster of Excellence EXC 2167 (Precision Medicine in Chronic Inflammation) for the mini proposal “Characterization of a genetic mouse model for systemic inflammation: A novel tool to unravel molecular hallmarks and therapeutic targets in sepsis?” (both to SR).

## Conflict of interests

The authors declare no conflict of interests.

## Supplementary Data

**Supplementary Figure 1:**
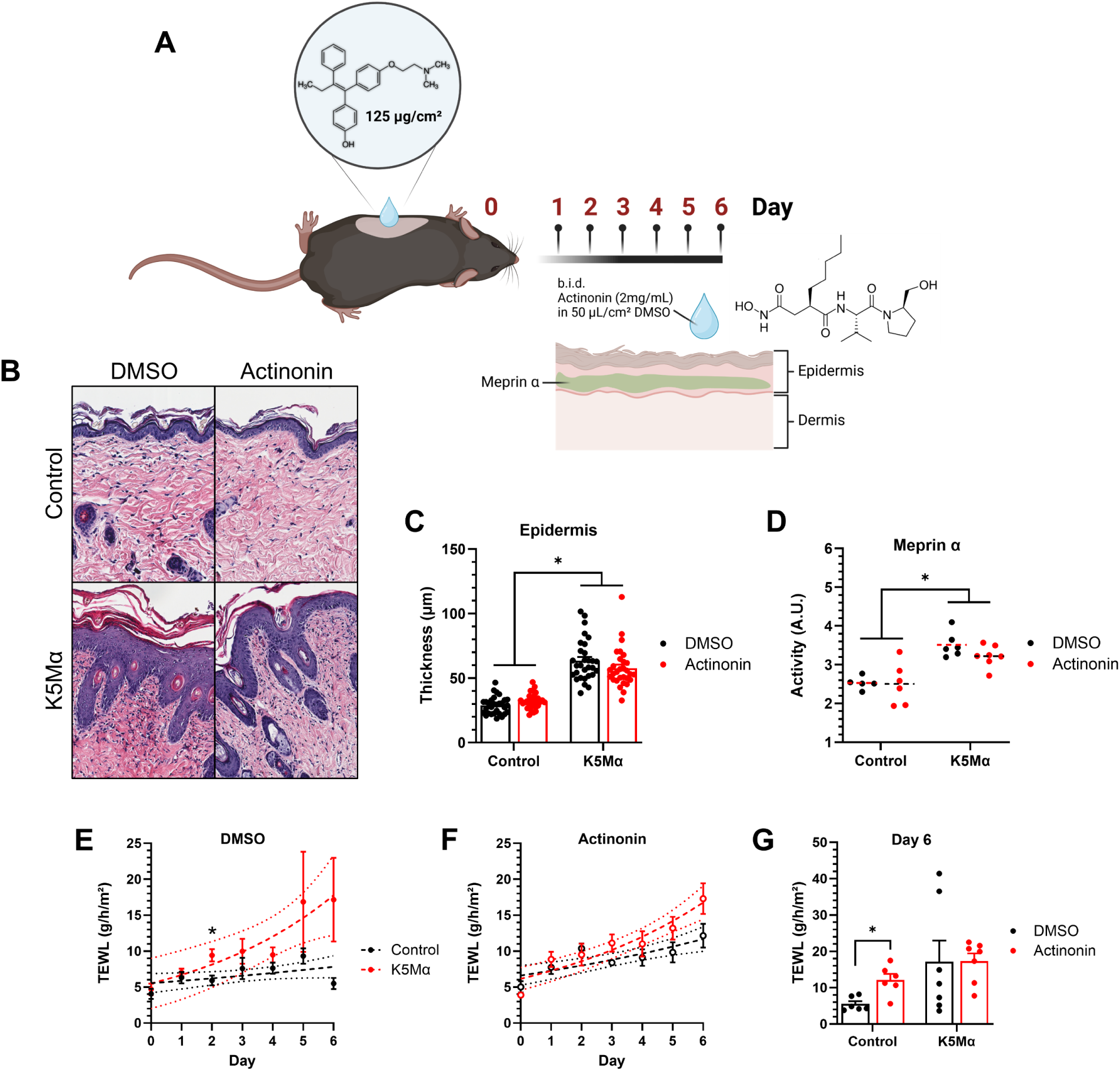
Topical actinonin treatment neither prevents nor delays development of pathological skin phenotype and barrier defect in K5Mα mice. **(A)** Illustration of the topical 4-hydroxytamoxifen treatment of control and K5Mα mice followed by actinonin treatment. Shaved skin areas were treated once on day zero with 125 µg/cm² 4-hydroxytamoxifen (TAM) in DMSO. The following six days, skin areas were treated twice daily (*bis in die* (b.i.d.)) with 50 µl/cm² actinonin in DMSO (2mg/ml). **(B)** Representative microscopical images of hematoxylin and eosin stained skin sections from control and K5Mα mice on day six after topical TAM treatment and daily treatment with actinonin. **(C)** Image-based quantification of the epidermal thickness in skin sections from control (n=6) and K5Mα (n=6) mice topically treated with DMSO (n=3 per group) or actinonin (n=3 per group) six days after TAM treatment. Bars and error bars display mean values and respective standard error of the mean (SEM). Data points display quantifications of ten areas per skin section (n=30 per group). **(D)** Meprin α activity in skin lysates of control (n=11) and K5Mα (n=12) mice topically treated with DMSO (n=5-6 per group) or actinonin (n=6 per group) six days after TAM treatment. Activity is plotted in arbitrary units (A.U.). Dashed lines (- - -) mark mean values of each group. **(E+F)** Transepidermal water-loss (TEWL) via the skin areas of control (n=11) and K5Mα (n=12) mice treated on day zero with TAM and the following days with **(E)** DMSO (n=5-6 per group) or **(F)** actinonin (n=6 per group) twice daily for six days. Data points and error bars display mean values and respective SEM. Results from simple linear regression (dashed lines) and respective 95% confidence band of the best-fit line (dotted lines) are displayed. **(G)** TEWL via the skin areas of control (n=11) and K5Mα (n=12) mice treated with DMSO or Actinonin on day six after TAM treatment. Bars and error bars display mean values and respective SEM. Statistically significant differences (p < 0.05) are labeled by an asterisk (*). Graphical illustration in **(A)** was created with BioRender.com.

**Supplementary Figure 2:**
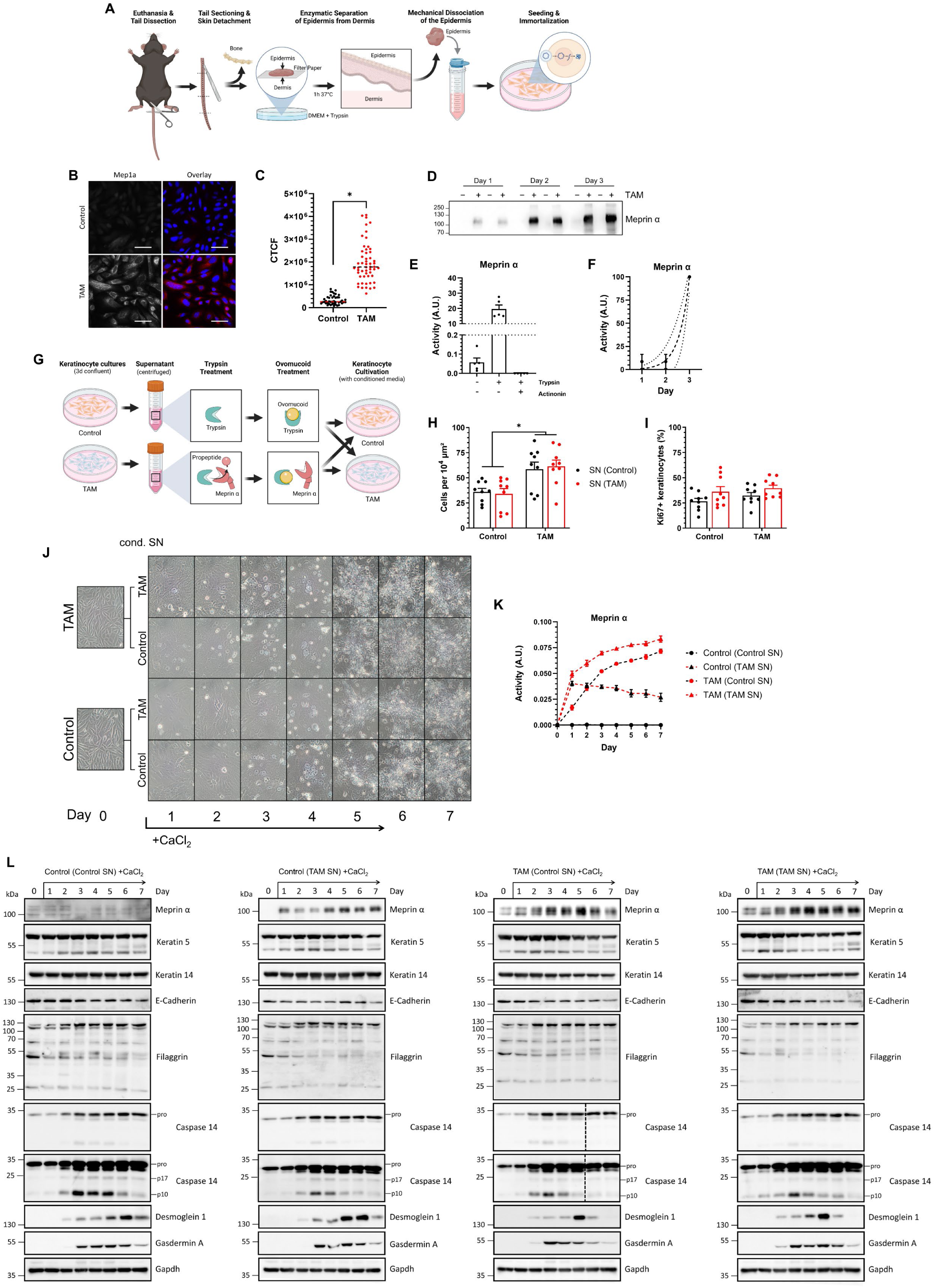
Meprin α overexpressing keratinocytes isolated from K5Mα mice show an increased proliferation but no defects in calcium-induced terminal differentiation. **(A)** Illustration of the keratinocyte isolation from tail epidermis of K5Mα mice. Mice were euthanized, tails were dissected and skin removed. Epidermis was separated from the dermis by treatment with trypsin, keratinocytes were dissociated in a cell strainer. Keratinocytes were immortalized with a plasmid encoding the large T antigen of the simian virus 40. **(B)** Representative immunofluorescence images of K5Mα keratinocytes treated with DMSO (Control) or 1 µM 4-hydroxytamoxifen in DMSO (TAM). Meprin α (grey levels or red in overlay) was detected by indirect immunofluorescence staining. Nuclei were stained with 4’,6-diamino-2-phenylindol (DAPI). Scale bars = 50 µm. **(C)** Image-based quantification of meprin α staining in K5Mα keratinocytes treated with DMSO (Control) or TAM. Data is displayed as corrected total cell fluorescence (CTCF) and each data point represents a single cell. Dashed lines (- - -) mark the mean values of each group. **(D)** Western blot detection of meprin α in the culture supernatant of K5Mα keratinocytes treated with DMSO (-) or TAM (+) after one, two and three days. Western blot analyses revealed that TAM-treated keratinocytes secreted meprin α, which accumulates within three days in the culture supernatant **(E)** Meprin α activity in the culture supernatant of K5Mα keratinocytes treated with TAM for three days. Culture supernatants (n=5 per group) were either left untreated, treated with trypsin (50 µg/ml) for activation of meprin α or treated with trypsin and actinonin (10 µM) for inhibition of activated meprin α. Meprin α is secreted as a zymogen, can be activated by trypsin and inhibited by actinonin. Activity is plotted in arbitrary units (A.U.). Bar charts and error bars display mean and standard error of the mean (SEM). **(F)** Meprin α activity in the culture supernatant of K5Mα keratinocytes on day one, two and three after TAM treatment (n=3 per day). Culture supernatants were treated with trypsin (50 µg/ml) for activation of meprin α. Data points and error bars display mean values and SEM. Results from non-linear regression based on an exponential growth equation (dashed line) as well as respective 95% confidence band of the best-fit line (dotted lines) are displayed. **(G)** Illustration of the preparation of conditioned culture medium. Culture supernatant from DMSO (Control) or TAM-treated keratinocytes was isolated, centrifuged and treated with trypsin for activation of meprin α. Afterwards, trypsin was inhibited by addition of ovomucoid. Resulting conditioned culture media were used for subsequent cultivation of K5Mα control or TAM-treated keratinocytes. **(H)** Cell count in control and TAM-treated keratinocyte cultures after cultivation in conditioned media (SN) for 48 h. Bar charts and error bars display mean and SEM. After 48 h of culturing control and TAM-treated keratinocytes in conditioned media, significantly higher cell numbers were found in the group of TAM-treated keratinocytes independent of the culture medium **(I)** Proportion of Ki67-positive keratinocytes in control and TAM-treated keratinocyte cultures after cultivation in conditioned media (SN) for 48 h. The proportion of Ki67-positive keratinocytes was higher in control and TAM-treated keratinocytes cultured in conditioned medium from TAM-treated keratinocytes than in those cultured in conditioned medium from control keratinocytes. Bar charts and error bars display mean and SEM. **(J)** Representative light microscopic images of control and TAM-treated keratinocytes cultured in conditioned media (cond. SN) supplemented with 540 µM CaCl_2_ for seven days. Light microscopical analyses revealed that keratinocytes in each condition underwent terminal differentiation without prominent temporary differences. **(K)** Meprin α activity in the culture supernatant of control and TAM-treated K5Mα keratinocytes cultured in conditioned media (SN) supplemented with 540 µM CaCl_2_ over seven days. No meprin α activity was detected in the supernatant of control keratinocytes treated with conditioned medium from control keratinocytes during terminal differentiation. In supernatants of control keratinocytes cultured in conditioned medium from TAM-treated keratinocytes, highest meprin α activity was observed on day one after addition of the conditioned medium and slightly decreasing activity on the following six days. Continuously increasing meprin α activity was detected in the supernatant of TAM-treated keratinocytes cultured in conditioned medium from control keratinocytes indicating that either keratinocytes endogenously synthesize an activator of meprin α upon CaCl_2_-induced terminal differentiation, or trypsin treatment for the production of conditioned media activated endogenous activators present in the supernatants. Data points and error bars display mean and SEM (n=3 per group per day). **(L)** Western blot detection of meprin α, keratin 14, E-cadherin, filaggrin, caspase 14, desmoglein 1 and gasdermin A in lysates of control and TAM-treated K5Mα keratinocytes cultured for seven days in conditioned media supplemented with 540 µM CaCl_2_ for induction of terminal differentiation. Western blot analyses did not provide any evidence that increased meprin α activity affects terminal differentiation. Within the seven days of terminal differentiation a decrease in the levels of basal keratinocyte markers keratin 5 and E-cadherin was observed. Proteins that are associated with and processed during terminal differentiation (filaggrin levels, caspase 14 levels and activation, desmoglein 1 and gasdermin A levels) did not reveal any prominent differences between the four groups but showed temporal changes as expected during terminal keratinocyte differentiation. Gapdh was detected as a reference. Western blot for the detection of caspase 14 in the lysate of TAM-treated keratinocytes cultured in conditioned medium from control keratinocytes was cut (indictated by dashed line) and rearranged for reasons of consistency. Statistically significant differences (p < 0.05) are labeled by an asterisk (*). Graphical illustrations in **(A)** and **(G)** were created with BioRender.com.

**Supplementary Figure 3:**
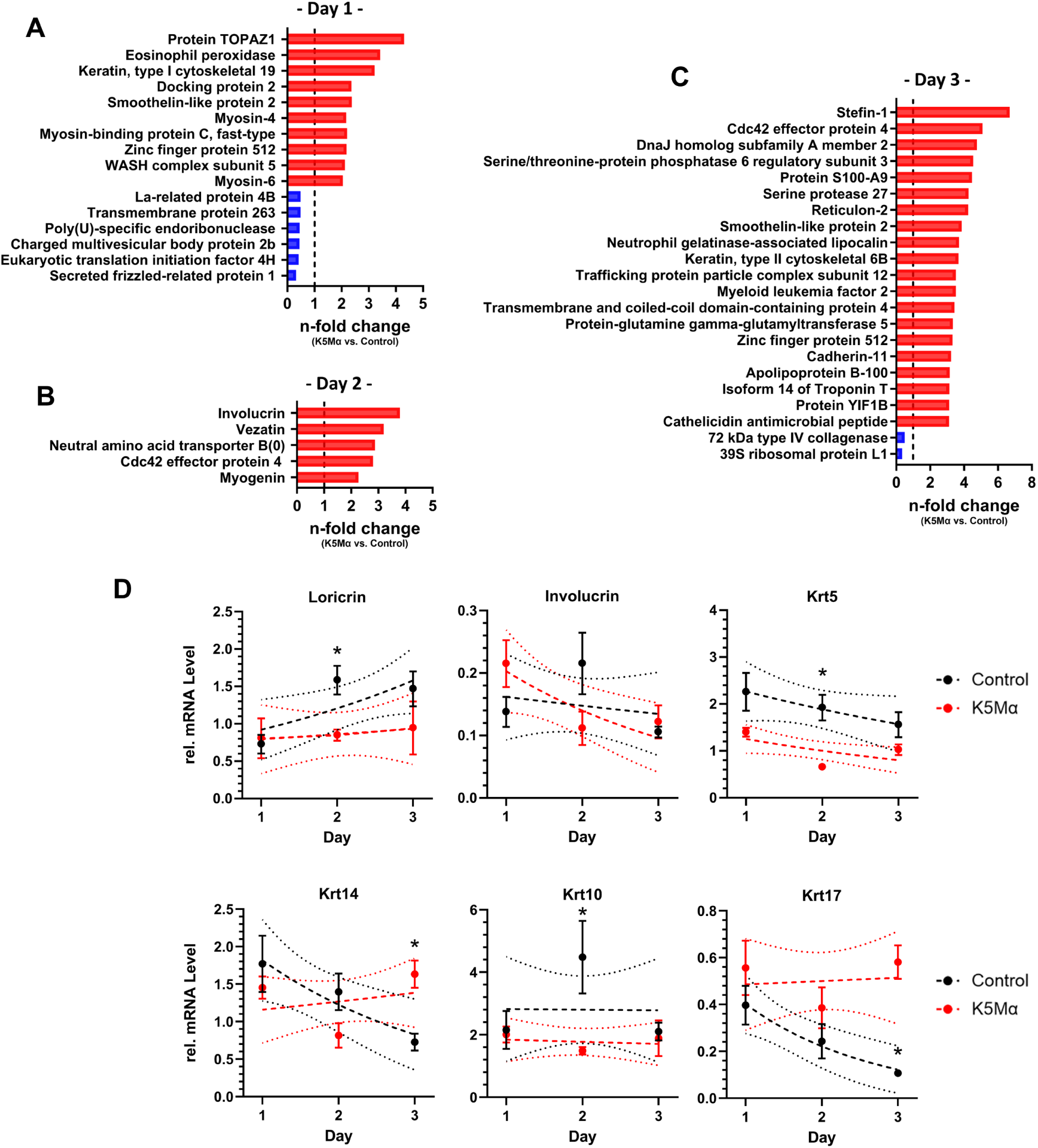
Alterations in the skin proteome of K5Mα mice. (A-C) Bar charts display n-fold changes (sorted by highest to lowest) in the levels of significantly higher (red bars; ≥ 2.0-fold) and lower (blue bars; ≤ 0.5-fold) abundant proteins detected by proteomics in the skin of K5Mα compared to control mice on day **(A)** one, **(B)** two and **(C)** three (only top 20 higher abundant proteins) after topical treatment with 4-hydroxytamoxifen (TAM). **(D)** Relative mRNA levels of *Loricrin*, *Involucrin*, *Krt5*, *Krt14*, *Krt10* and *Krt17* in skin samples from control (n=5 per day) and K5Mα (n=5 per day) mice at days one to three after TAM treatment. Relative messenger RNA levels were normalized to *Gapdh* levels in the respective sample by 2^-ΔΔCt^-method. Data points and error bars display mean values and respective standard errors of the mean (SEM). Results from simple linear regression and non-linear regression based on an exponential growth equation (dashed lines) as well as respective 95% confidence band of the best-fit line (dotted lines) are displayed. Statistically significant differences (p < 0.05) are labeled by an asterisk (*).

**Supplementary Figure 4:**
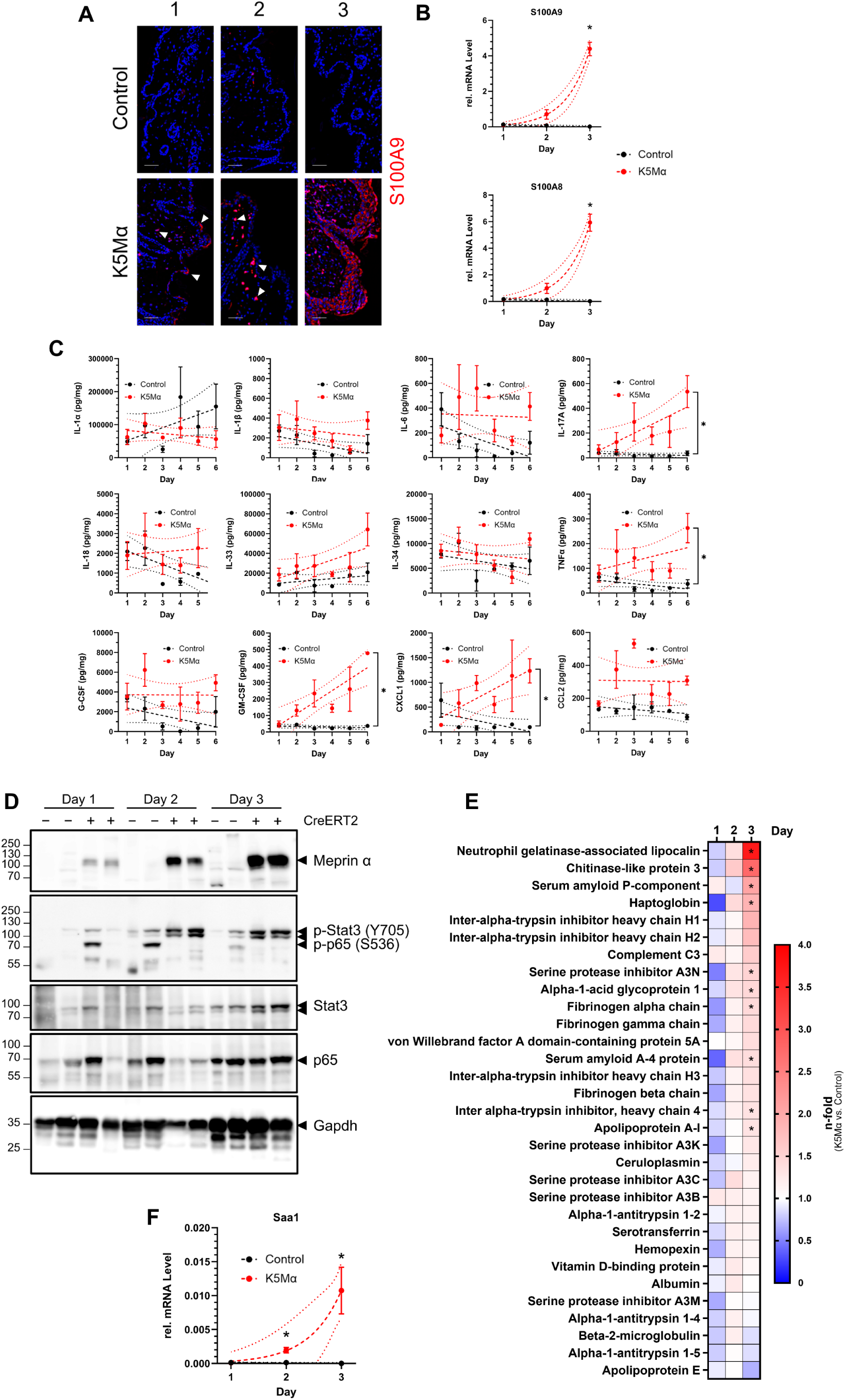
Inflammatory cytokines and acute phase response proteins in the skin of K5Mα mice. **(A)** Representative immunofluorescence images of skin sections from control and K5Mα mice on day one to three after topical TAM treatment. S100A9 (red) was detected by indirect immunofluorescence staining. Nuclei were stained with 4’,6-diamino-2- phenylindol (DAPI). Scale bars = 50 µm. **(B)** Relative mRNA levels of *S100a9* and *S100a8* in skin samples from control (n=5 per day) and K5Mα (n=5 per day) mice at days one to three after TAM treatment. Relative messenger RNA levels were normalized to *Gapdh* levels in the respective sample by 2^-ΔΔCt^-method. Data points and error bars display mean values and respective standard errors of the mean (SEM). Results from simple linear regression and non-linear regression based on an exponential growth equation (dashed lines) as well as respective 95% confidence band of the best-fit line (dotted lines) are displayed. **(C)** Concentrations of IL-1α, IL-1β, IL-6, IL-17A, IL-18, IL-33, IL-34, TNFα, G-CSF, GM-CSF, CXCL1 and CCL2 in skin lysates of control (n=3) and K5Mα (n=3) mice on day one to six after topical treatment with 4-hydroxytamoxifen (TAM). Analyte concentrations were normalized to the respective protein concentration in the lysates. Data points and error bars display mean values and respective standard errors of the mean (SEM). Results from simple linear regression (dashed lines) and respective 95% confidence band of the best-fit line (dotted lines) are displayed. **(D)** Western blot detection of meprin α, phospho-Stat3 (Y705), phospho-p65 (S536), total Stat3 and total p65 in skin lysates of control (CreER^T2^ -; n=2) and K5Mα (CreER^T2^ +; n=2) mice on day one, two and three after TAM treatment. Gapdh was detected as a reference. **(E)** Heatmap displays the mean n-fold changes in the level of acute phase response proteins detected by LC-MS/MS in the skin of K5Mα mice compared to control mice on day one, two and three after TAM treatment. Proteins higher (red) and lower (blue) abundant in the skin of K5Mα than in control mice are highlighted by color gradients. **(F)** Relative mRNA level of *Saa1* in skin samples from control (n=5 per day) and K5Mα (n=5 per day) mice at days one to three after TAM treatment. *Saa1* mRNA levels were normalized to *Gapdh* levels in the respective sample by 2^-ΔΔCt^-method. Data points and error bars display mean values and respective standard errors of the mean (SEM). Results from non-linear regression based on an exponential growth equation (dashed lines) and respective 95% confidence band of the best-fit line (dotted lines) are displayed. Statistically significant differences (p < 0.05) are labeled by an asterisk (*).

**Supplementary Figure 5:**
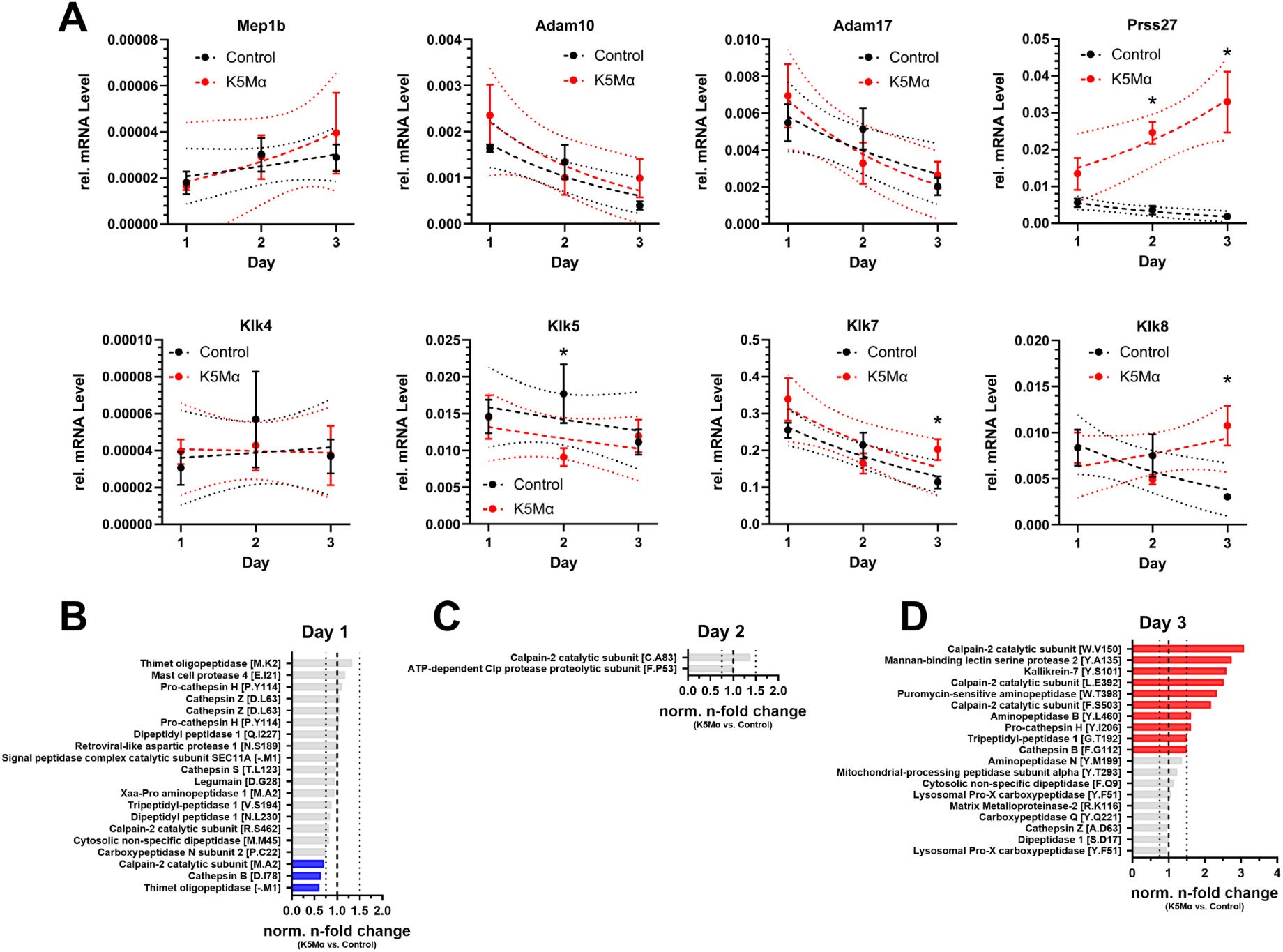
Protease and peptidase levels in the skin of K5Mα mice. **(A)** Relative mRNA levels of *Mep1b*, *Adam10*, *Adam17*, *Prss27*, *Klk4*, *Klk5*, *Klk7* and *Klk8* in skin samples from control (n=5 per day) and K5Mα (n=5 per day) mice at days one to three after topical treatment with 4-hydroxytamoxifen (TAM). Relative messenger RNA levels were normalized to *Gapdh* levels in the respective sample by 2^-ΔΔCt^-method. Data points and error bars display mean values and respective standard errors of the mean (SEM). Results from simple linear regression (dashed lines) and respective 95% confidence band of the best-fit line (dotted lines) are displayed. **(B-D)** Bar charts display mean n-fold changes in the levels of peptides (annotated by corresponding proteins and putative cleavage site in squared brackets) derived from proteases and peptidases detected by TAILS in the skin of K5Mα compared to control mice on day **(B)** one, **(C)** two and **(D)** three after topical TAM treatment. Peptide levels were normalized to the total level of corresponding proteins detected by proteomics analysis. Equal normalized n-fold levels (n-fold=1) are highlighted by a dashed line (- - -) and thresholds for 0.75-fold and 1.5-fold altered levels are marked by dotted line (· · ·). Bar colors highlight higher (> 1.5-fold; red) and lower (< 0.5-fold; blue) abundant peptides. Peptides detected at similar levels (0.5-fold < n-fold < 1.5-fold) in the skin of control and K5Mα mice are presented as grey bars. Statistically significant differences (p < 0.05) are labeled by an asterisk (*).

**Supplementary Figure 6:**
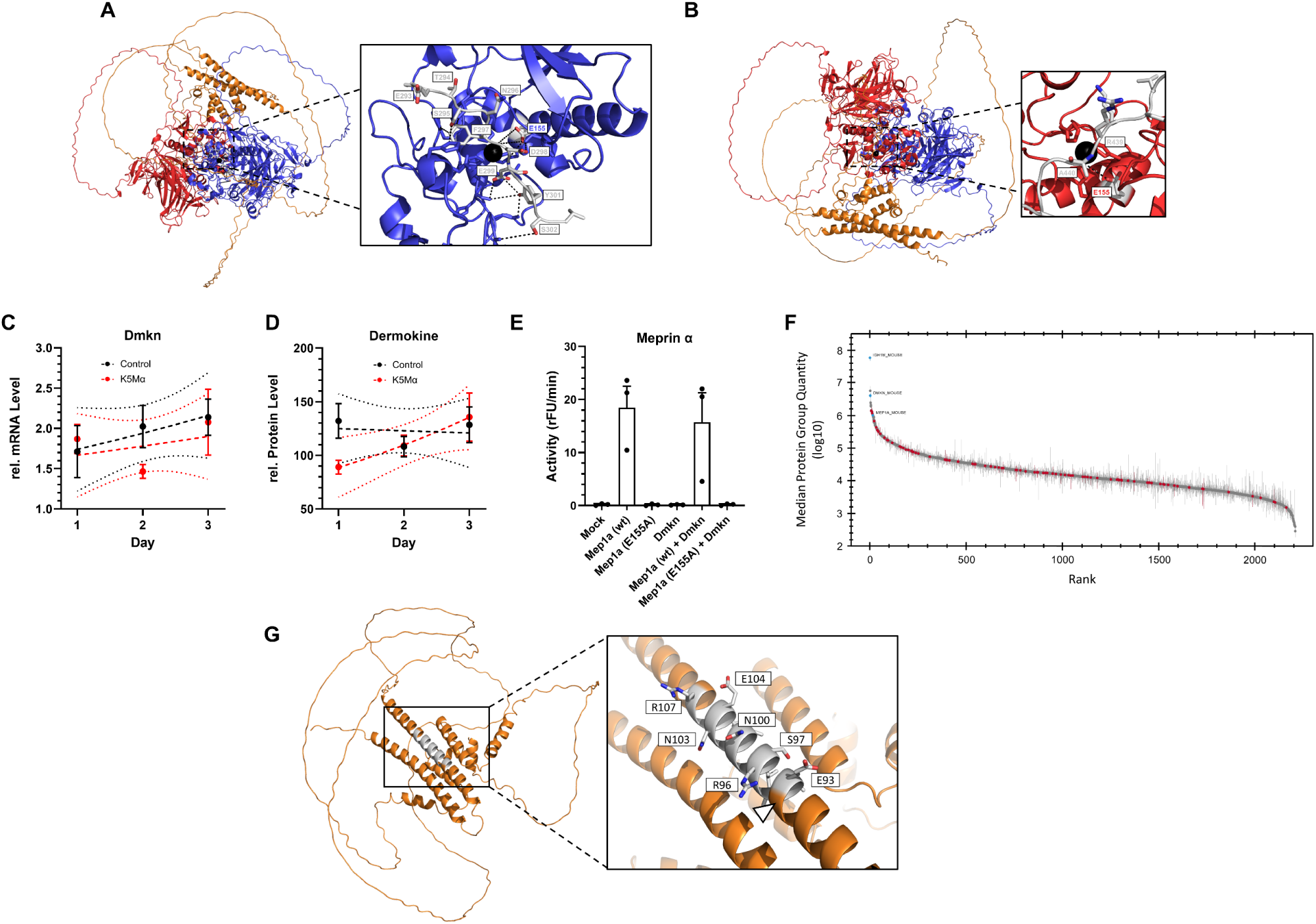
Structural prediction of dermokine in complex with meprin α. (A+B) AlphaFold 3 model of murine dermokine β (UniProt ID: Q6P253-1, without N-terminal signal peptide sequence) in complex with the homodimeric ectodomain of murine meprin α (P28825, ectodomain without N-terminal signal and propeptide sequences (N65 to M713). Dermokine (orange) and meprin α subunits (red and blue) are presented as ribbon diagrams. Zoom-ins show the orientation and interaction of dermokine in the active site of meprin α. Side chains of dermokine are color coded by elements (carbon (grey), oxygen (red) and nitrogen (blue)) and amino acids in one-letter code as well as respective position are annotated (grey). Side chain of the catalytic glutamate (E155) within the active site of meprin α is also presented, color coded and annotated (**(A)** blue or **(B)** red). Zinc ion is presented as black sphere. Predicted non-covalent interactions between dermokine and meprin α are highlighted by dashed lines (- - -): **(A)** Hydrogen bonds between S295-H155, D298-C127, E299-H212, Y301-G239, Y301-H212, Y301-L210 and S302-F216. Binding of dermokine F297 into a pocket formed by meprin α F163 and I131. Distance of meprin α E155 side chain to the dermokine peptide bonds between F297-D298 and D298-E299. **(B)** Distance of meprin α E155 side chain to the dermokine peptide bond between R439 and A440. **(C)** Relative mRNA levels of *Dmkn* in skin samples from control (n=5 per day) and K5Mα (n=5 per day) mice at days one to three after TAM treatment. *Dmkn* levels were normalized to *Gapdh* levels in the respective sample by 2^-ΔΔCt^-method. Data points and error bars display mean values and respective standard errors of the mean (SEM). Results from simple linear regression (dashed lines) and respective 95% confidence band of the best-fit line (dotted lines) are displayed. **(D)** Relative dermokine protein levels detected by LC-MS/MS in skin samples from control (n=5 per day) and K5Mα (n=5 per day) mice at days one to three after TAM treatment. Data points and error bars display mean values and respective SEM. Results from simple linear regression (dashed lines) and respective 95% confidence band of the best-fit line (dotted lines) are displayed. **(E)** Meprin α activity in the culture supernatant of HEK293T cells transiently transfected with an empty control plasmid (Mock) or plasmids encoding wildtype (wt) murine meprin α (*Mep1a*), C-terminally Myc-FLAG-tagged murine dermokine (*Dmkn*) and/or a catalytically inactive meprin α variant (*Mep1a* (E155A)). Activity is plotted as change in relative fluorescence units (rFU) per minute. Bar charts and error bars display mean and standard error of the mean (SEM) (n = 3 per group). **(F)** Median quantity of proteins detected by mass spectrometry in lysates of HEK293T cells transiently transfected with plasmids encoding *Mep1a* wt and *Dmkn* after co-immunoprecipitation (co-IP) of dermokine. Proteins are grouped and ranked according to their abundance. Highlighted are murine Ig-gamma chain C region (IGH1M_MOUSE; antibody used for co-IP), murine dermokine (DMKN_MOUSE) and murine meprin α (MEP1A_MOUSE). **(G)** AlphaFold 3 model of murine dermokine β (UniProt ID: Q6P253-1, without N-terminal signal peptide sequence) presented as ribbon diagram. Zoom-in highlights the cleavage site by meprin α (Δ) as well as the peptide identified by mass spectrometry after co-IP. Side chains of the peptide are color coded by elements (carbon (grey), oxygen (red) and nitrogen (blue)) and amino acids in one-letter code as well as respective position are annotated.

## Notes

https://www.ebi.ac.uk/pride/archive

### Competing Interest Statement

The authors have declared no competing interest.

